# Immune-mediated Tubule Atrophy Promotes Acute Kidney Injury to Chronic Kidney Disease Transition

**DOI:** 10.1101/2022.06.01.494328

**Authors:** Leyuan Xu, Jiankan Guo, Dennis G. Moledina, Lloyd G. Cantley

**Affiliations:** Department of Internal Medicine/Section of Nephrology, Yale University School of Medicine

**Keywords:** Ischemia/reperfusion, acute kidney injury, chronic kidney disease, immune response, inflammation

## Abstract

Incomplete repair after acute kidney injury (AKI) is associated with progressive loss of tubular cell function and development of chronic kidney disease (CKD). Here, we compared mice subjected to either unilateral ischemia-reperfusion kidney injury with contralateral nephrectomy (IRI/CL-NX, in which tubule repair predominates) or unilateral IRI with contralateral kidney intact (U-IRI, in which fibrosis and atrophy predominates) to investigate the mechanism(s) underlying transition to CKD following AKI. The initial injury and early recruitment and activation of macrophages, dendritic cells (DCs), neutrophils, and T cells were similar through day 7 but markedly diverged afterwards between the two models. By day 14, kidneys subjected to U-IRI had greater numbers of macrophages with higher expression of *Ccl2*, *Ccl7*, *Ccl8*, *Ccl12*, and *Cxcl16*. These chemokines correlated with a second wave of *Ccr1*-positive neutrophils and *Cxcr6*-positive T cells, resulting in a proinflammatory milieu, accompanied by increased expression of tubular cell injury, oxidative stress and major histocompatibility complex genes. This second wave of immune dysfunction led to a distinct profile of tubule injury with morphologic kidney atrophy and a decreased proportion of differentiated tubule cells. Combined depletion of neutrophils and T cells beginning on day 5 after U-IRI was found to reduce tubular cell loss and the associated kidney atrophy. In kidney biopsy samples from patients with AKI, the number of interstitial T cells and neutrophils negatively correlated with 6-month recovery of GFR. Together, our findings demonstrate that macrophage persistence after AKI promotes a T cell- and neutrophil-mediated proinflammatory milieu that leads to progressive tubule damage.

## INTRODUCTION

Acute kidney injury (AKI) is the syndromic term used to describe the abrupt reduction in glomerular filtration rate (GFR) caused by an insult such as ischemia, sepsis, nephrotoxin exposure or urinary obstruction. Based on animal models, these insults often precipitate a rapid innate immune response dominated by neutrophil and proinflammatory macrophage recruitment with significant tubular and endothelial injury, leading to GFR reduction even after correction of the inciting event^1–6^. Under ideal conditions (e.g., transient insults in young healthy subjects), repair pathways that include neutrophil egress and a switch to reparative macrophage activation then promote clearance of cell debris and proliferation of surviving tubular cells, leading to restoration of nephron structure and GFR^7–10^. However, in many cases this repair phase is incomplete and includes maladaptive processes such as interstitial fibrosis and tubule atrophy^11^. As a result, people who survive AKI are at an 8.8-fold increase in risk for chronic kidney disease (CKD) and a 3.3-fold increase in risk for end stage renal disease (ESRD)^12–14^. To date, no therapy is available to improve the post-injury repair process or prevent progression from AKI to CKD. Defining the signals that promote maladaptive kidney repair following AKI may therefore help us identify therapeutic targets to prevent or even reverse tubule atrophy and the progression to CKD.

Investigations of animal models in which the initial injury stimulus is sustained, such as unilateral ureteral obstruction (UUO), have shown that macrophages progressively accumulate in the kidney interstitium adjacent to unrepaired tubules and promote kidney fibrosis^15–17^. However, the underpinnings of kidney fibrosis and atrophy following transient insults, as is frequently seen in hospitalized patients, are currently unknown. In models of transient injury such as bilateral ischemia-reperfusion injury (IRI), or unilateral IRI with the contralateral nephrectomy (IRI/CL-NX), kidneys undergo a biphasic response of initial tubule cell death driven in part by an innate proinflammatory activation, followed by tubule repair and restoration of normal or near-normal function^7–10^. However, using the model of unilateral IRI with the contralateral kidney left intact (U-IRI), we and others have shown that the injured kidney undergoes progressive atrophy and fibrosis rather than successful repair^18–21^. These transiently injured kidneys continue to exhibit large numbers of macrophages long after the tubule repair phase is complete (days 7-10). Knock-out of either the macrophage survival factor chitinase 3-like 1 (*Chi3l1* or *Brp-39*) or *Ccr2*, the chemokine receptor for monocyte chemoattractant protein-1 (*Mcp1* or *Ccl2*) reduces macrophage numbers and the degree of fibrosis^20, 21^, but does not significantly impact the degree of tubule or kidney atrophy^21^. This disconnection between fibrosis and atrophy suggests that the progressive loss of kidney tubules during AKI-to-CKD transition involves mechanisms in addition to accumulation of extracellular matrix^22^.

To better understand the pathogenesis of AKI-to-CKD transition and specifically the mechanism of kidney tubule atrophy, we compared the kidney response to identical times of ischemic injury between mice subjected to U-IRI (to induce atrophy) or IRI/CL-NX (to induce adaptive repair). We performed single cell RNA-sequencing (scRNA-seq) analysis on day 7, 14, and 30 after injury to identify major cell types in the kidney and the differential transcriptional response between the models in each cell type. We confirmed our previous finding that U-IRI leads to macrophage persistence beyond the period of normal repair, and now show that those macrophages express high levels of T cell and neutrophil activating chemokines including *Cxcl16* and *Mcp2* (*Ccl8*), corresponding with a second wave of infiltrating *Cxcr6*+ T cells and *Ccr1*+ neutrophils in the interstitium. This late recruitment of T cells and neutrophils closely associated with a proinflammatory milieu including *Tnf* and *Il1b*. Concomitantly, the tubular cells from U-IRI kidneys expressed higher levels of injury markers such as vascular cell adhesion molecule 1 (*Vcam1*) and class I & II major histocompatibility (MHC) genes and exhibited a dedifferentiated expression profile, correlating with late kidney atrophy. Depletion of T cells and neutrophils together, but not individually, was found to attenuate the second wave of injury and partially restore tubule mass in the U-IRI model. Consistent with the mouse models, we found that increasing numbers of T cells and neutrophils in the renal interstitium at the time of renal biopsy for AKI negatively correlated with 6-month recovery of GFR. Together, these findings suggest that failed tubule repair leads to macrophage persistence with a second wave of T cell and neutrophil-dependent proinflammatory immune activation that induces secondary tubule injury and promotes kidney atrophy during AKI-to-CKD transition.

## RESULTS

### U-IRI leads to tubule atrophy and tubule cell dedifferentiation

Phenotypically, the injured kidneys from the IRI/CL-NX mice had hypertrophied by day 7 after injury; whereas the injured kidneys from the U-IRI mice were equal in size to controls on day 7, but progressively atrophied afterwards (Fig. 1a and Supplementary Fig. 1a-c). By day 30, IRI/CL-NX kidneys weighed 38% more than age-matched control kidneys and the same as kidneys 30 days after contralateral nephrectomy alone (Supplementary Fig. 1d), while U-IRI kidneys weighed 58% less than control kidneys and 69% less than IRI/CL-NX kidneys (Fig. 1b). This resulted in a 38% and 46% decrease in cross-sectional area of the U-IRI kidneys relative to control and IRI/CL-NX kidneys, respectively (Fig. 1c). Staining for the general tubular epithelial cell marker KSP-cadherin^23, 24^ revealed a 56% reduction in absolute tubular area 30 days after U-IRI as compared to IRI/CL-NX (Fig. 1d,e), however, the proportion of renal parenchymal area comprised of tubules was only 12% lower in the atrophied U-IRI kidneys than in the hypertrophied IRI/CL-NX kidneys (Fig. 1e). This suggests that kidney atrophy after AKI is predominantly due to reduced tubular mass rather than replacement of tubule epithelia by fibrosis^21, 25^. To assess the differentiation state of the remaining tubules, LTL and megalin staining was performed to quantify the area of preserved proximal tubule brush border^26, 27^. This showed that PT brush border area comprised approximately 29% less of the remaining renal parenchymal area 30 days after U-IRI compared to IRI/CL-NX (Fig. 1f-i). Quantitative PCR and megalin IF staining analyses confirmed that both mRNA expression and protein levels of the differentiated proximal tubular brush border proteins megalin (*Lrp2*), NaPi-IIa (*Scl34a1*, mRNA level alone), and NaDC3 (*Slc13a3*, mRNA level alone) were significantly lower at 14 and 30 days in the U-IRI kidneys as compared to IRI/CL-NX kidneys (Fig. 2a-c).

**Figure 1.**
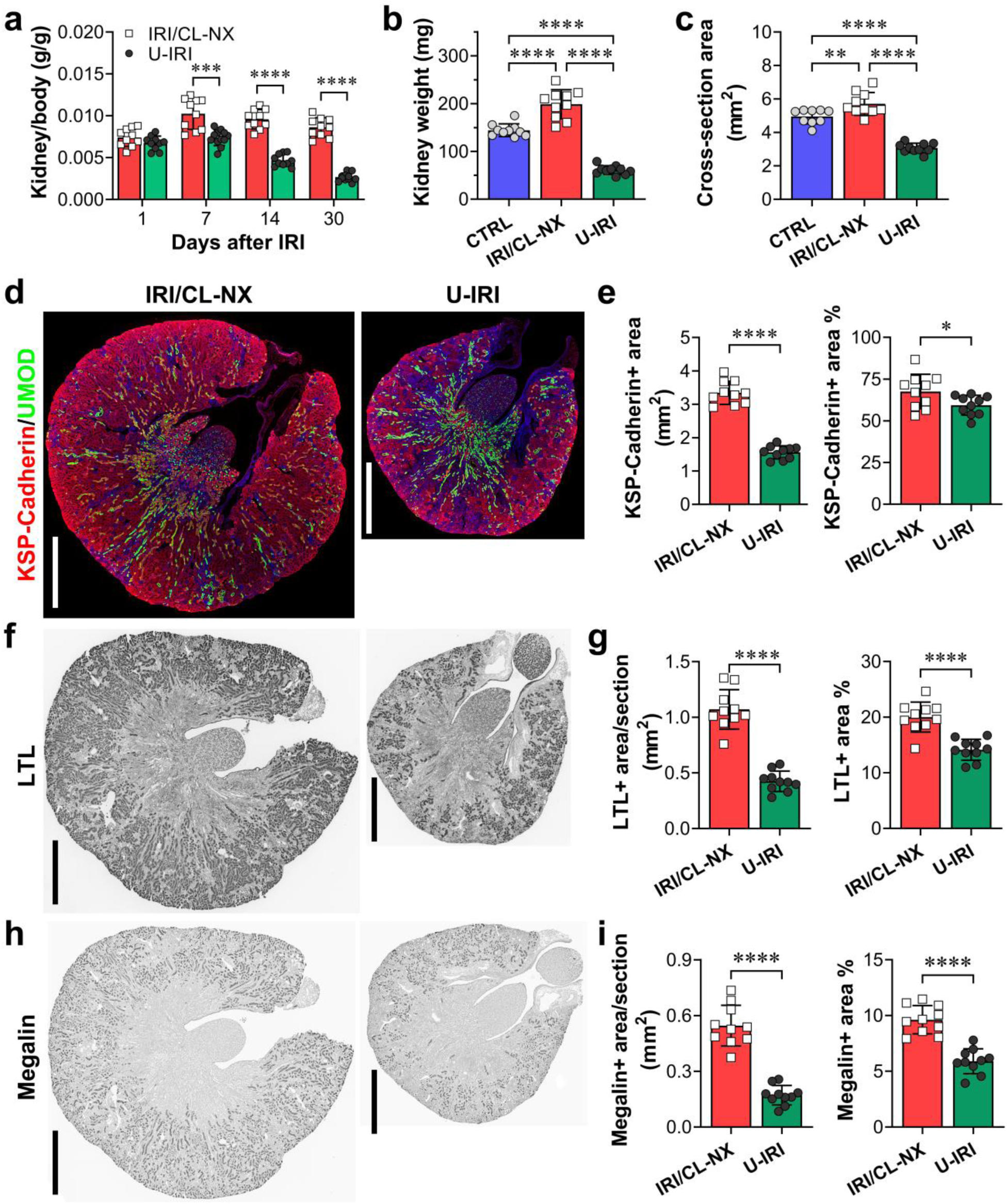
U-IRI leads to tubule atrophy. Wild-type mice were subjected to 27 minutes of IRI with contralateral nephrectomy (IRI/CL-NX) or unilateral IRI (U-IRI) and sacrificed on day 1, 7, 14 and 30 after injury. **a.** Kidney-to-body weight ratios were determined on day 1, 7, 14, and 30 after injury. n=10 kidneys/time point. p<0.0001 between models and in time series; ***p<0.001 in the indicated subgroup analyses. **b.** Kidney weights were determined on day 30 after injury. CTRL, age matched control. n=9-10 kidneys/time point. ****p<0.0001. **c.** Midline kidney cross-section area was determined on day 30 after injury. CTRL, age matched control. n=9-10 kidneys/time point. **p<0.01, ****p<0.0001. **d.** Midline kidney sections on day 30 after IRI were co-stained with KSP-cadherin (red), UMOD (green), and DAPI (blue). Scale bars, 1 mm. **e.** KSP-Cadherin-positive area as in (d) was quantified for the entire kidney section (left panel) and as a percentage of the section area (right panel). n=10 kidneys/group. *p<0.05, ****p<0.0001. **f.** Midline kidney sections underwent IHC staining for LTL (dark grey) on day 30 after IRI. Scale bars, 1 mm. **g.** LTL-positive area as in (f) was quantified for the entire kidney section (left panel) and as a percentage of the section area (right panel). n=10 kidneys/group. ****p<0.0001. **h.** Midline kidney sections underwent IHC staining for megalin (dark grey) on day 30 after IRI. Scale bars, 1 mm. **i.** Megalin-positive area as in (h) was quantified for the entire kidney section (left panel) and as a percentage of the section area (right panel). n=10 kidneys/group. ****p<0.0001.

**Figure 2.**
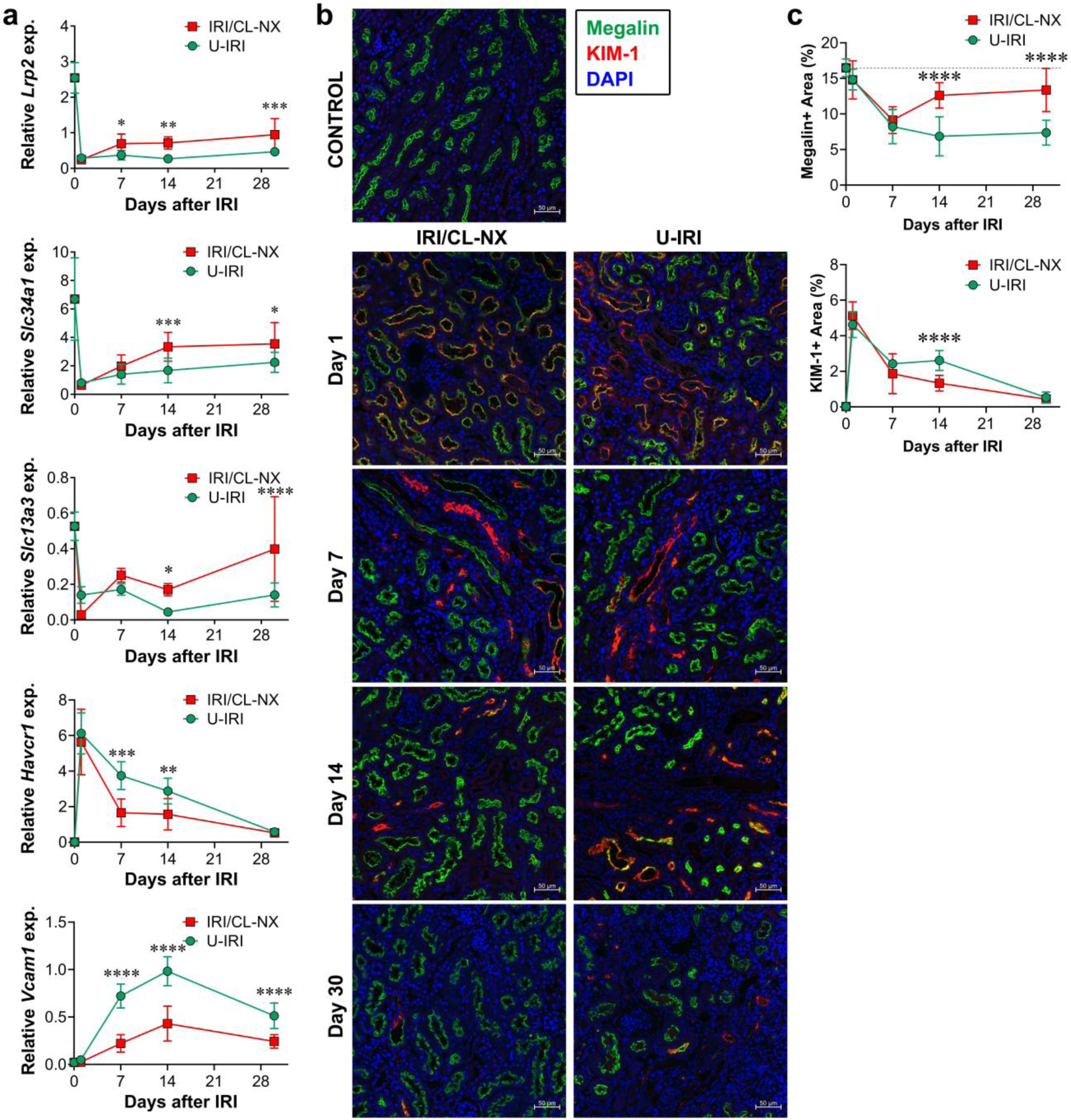
U-IRI leads to failure of tubular repair and tubular dedifferentiation. **a.** Quantitative RT-PCR analysis for *Lrp2* (megalin), *Slc34a1* (Napi2a), *Slc13a3* (NaDC3), *Havcr1* (Kim1), and *Vcam1* was performed on whole kidney RNA harvested on day 0, 1, 7, 14, and 30 after injury. n=10 kidneys/time point/group. Two-way ANOVA summarized in Supplementary Table 1. *p<0.05, ***p<0.01, ***p<0.001 at each time point. **b.** Midline kidney sections underwent IF staining for megalin (green) and KIM-1 (red) on day 0, 1, 7, 14, and 30 after IRI and the representative images were shown at 20×. Scale bars, 50 μm. **c.** Megalin-(top) and KIM-1-(bottom) positive areas as in (b) were quantified as a percentage of the 6-10 randomly selected area. n=8 kidneys/group. Two-way ANOVA [p < 0.0001 (interaction, time factor, and model factor) for megalin; p = 0.0003 (interaction), p <0.0001 (time factor), p = 0.0228 (model factor) for KIM-1]. ****p<0.0001.

Both mRNA and protein expression levels of the proximal tubule injury marker *Kim1* (*Havcr1*) were equally upregulated on day 1 after IRI in both models, confirming that the initial ischemic injury was equivalent following U-IRI and IRI/CL-NX (Fig. 2a-c and Supplementary Fig. 2)^28^. Of note, serum levels of both KIM-1 and NGAL were significantly higher on day 1 after IRI/CL-NX than after U-IRI, likely due to clearance by the intact contralateral kidney in mice subjected to U-IRI (Supplementary Fig. 3). However, *Kim1* remained substantially higher in the kidneys subjected to U-IRI on days 7 and 14 at the mRNA level and day 14 at the protein level (Fig. 2a-c). As compared to Kim1, *Vcam1* was not upregulated on day 1 after IRI in either model but markedly increased afterwards, and was significantly higher in the U-IRI kidneys that IRI/CL-NX kidneys from day 7 to day 30 (Fig. 2a), correlating with the period of progressive atrophy seen in this model (Fig. 1b). Together, these data suggest that tubular cells are subjected to equivalent initial ischemic injury in both models correlating with Kim1 expression, followed by delayed injury and failure to redifferentiate that correlates with *Vcam1* expression and is exaggerated in the U-IRI model.

### Kidney atrophy correlates with a second wave of immune activation and proximal tubule cell loss

Because the difference in kidney injury marker expression between the two models became apparent on day 7 (Fig. 2a), we performed single cell RNA sequencing (scRNA-seq) of the injured kidneys on days 7, 14, and 30 after IRI to identify the transcriptional and cellular differences underlying tubule repair and tubule atrophy. Unsupervised clustering generated twenty seven distinct cell types with gene expression profiles identifying them as proximal tubule (PT-S1, -S2, and -S3), injured PT, thick ascending limb (TAL), DCT/CNT (distal convoluted tubule/connecting tubule), collecting duct-principal cells (CD-PC), collecting duct-intercalated cells (CD-IC), endothelium, myofibroblasts, six clusters of macrophages (monocyte, infiltrating, M1, M2, proliferating, and resident macrophages), four clusters of dendritic cells [plasmacytoid DC (pDC), conventional DCs (cDC1 and cDC2), and proliferating cDC1], two clusters of neutrophils (Ngal high and low), four clusters of T cells [naïve, Cd4+ T helper/ regulatory T (Th/Treg), Cd8a+ cytotoxic T/natural killer T (Tc/NKT), and proliferating T cells], and B cells (Fig. 3a,b).

**Figure 3.**
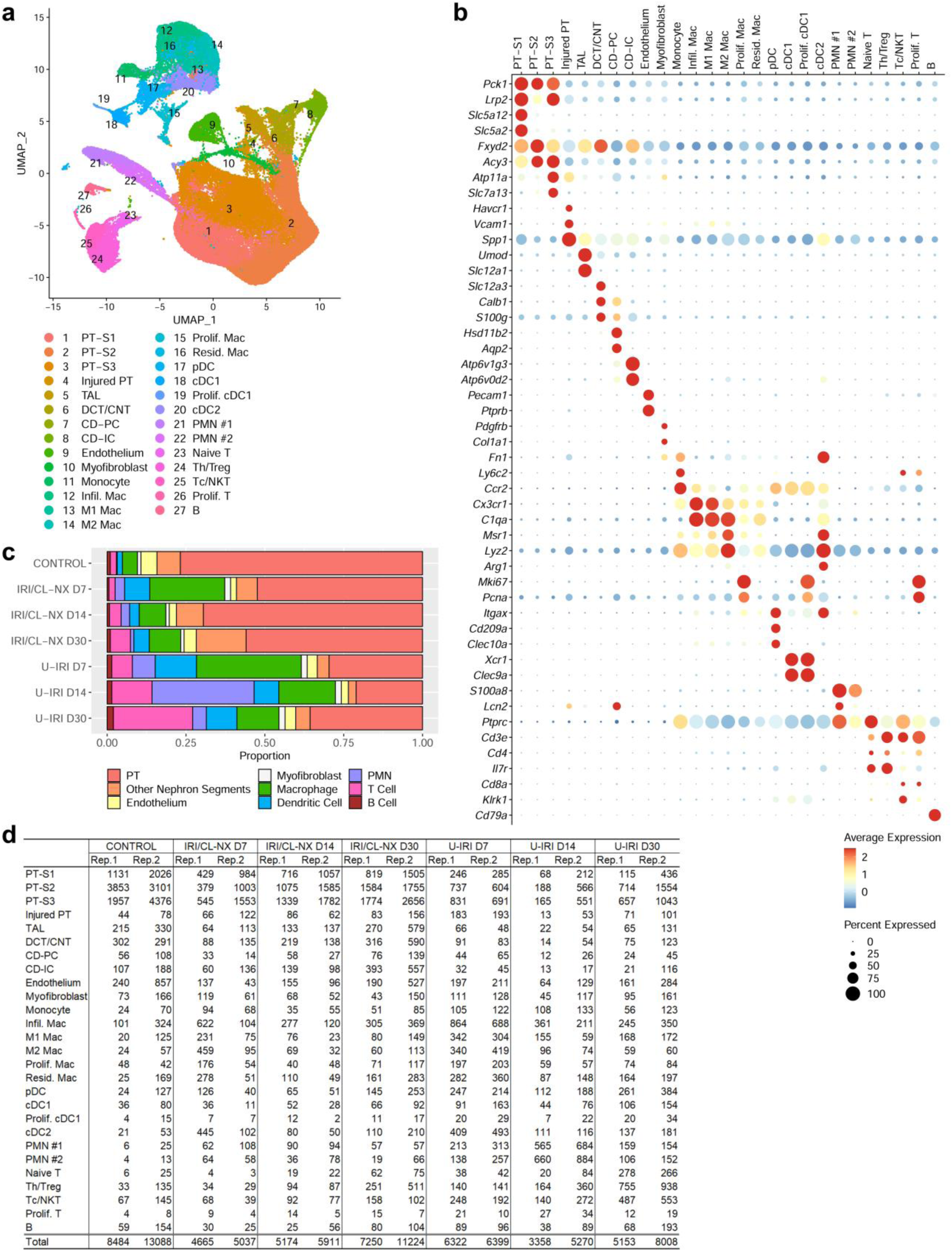
Integrated scRNA-seq analysis of differential cell type population between IRI/CL-NX and U-IRI kidneys and control kidneys. **a.** UMAP projection of 95,343 cells from integrated kidneys. Cell clusters were identified using the composite data from all cells by kidney cell and immune cell lineage-specific marker expression as shown in (**b**). PT, proximal tubule; TAL, thick ascending limb; DCT, distal convoluted tubule; CNT, connecting tubule; CD-PC, collecting duct-principal cell; CD-IC, collecting duct-intercalated cell; Infil. Mac, infiltrating macrophage; pDC, plasmacytoid dendritic cell; cDC, conventional dendritic cell; PMN, polymorphonuclear neutrophils; Th/Treg, T helper/regulatory T cells; Tc/NKT, cytotoxic T/natural killer T cells; B, B cells. **c.** The percentage of the cell populations is provided for each group. **d.** Table of the number of cells in each cluster in each kidney sample. Rep., replicate.

The kidney atrophy seen on day 14 and 30 after U-IRI was found to correspond to a decrease in tubular cell populations relative to the total cells present, with the greatest loss occurring in the PT cell compartment beginning by day 7 (Fig. 3c,d and Supplementary Fig. 4). Along with tubular cell loss, U-IRI kidneys exhibited an increased percentage of immune cells including macrophages, cDCs, neutrophils, and T cells as compared to the IRI/CL-NX kidneys. Of the immune cells identified, macrophages and DCs were the predominant cell types in the kidney on day 7 after injury in both models. However, both neutrophils and T cell populations markedly increased by day 14 after U-IRI as compared to those cells in IRI/CL-NX kidneys.

Consistent with the scRNA-seq analysis, quantitative PCR of whole kidney RNA revealed that markers for T cells (*Cd3e, Cd4* and *Cd8a*), neutrophils (*Ly6g*), macrophages (*F4/80* and *Cd68*) and dendritic cells (*Itgax*) significantly increased in the U-IRI model after day 7, with T cell, macrophage and dendritic cell markers remining high through day 30 (Fig. 4a, Supplementary Fig. 5). IHC staining for F4/80, CD11c, Ly6G, CD3ε, CD4, and CD8α on day 14 after injury confirmed that the numbers of interstitial macrophages, DCs, neutrophils, CD4+ T helper cells, and CD8+ cytotoxic T cells were significantly higher in the cortex of U-IRI kidneys as compared to contralateral and IRI/CL-NX kidneys, and all but Ly6G were higher in the outer medulla in the U-IRI model (Fig. 4b, 4c, Supplementary Fig. 6-11). Together, these data demonstrate that macrophages and dendritic cells are the major cell responders in the first 7 days after IRI in both injury models, with the number of macrophages, T cells and neutrophils plateauing between 7 and 14 days in the IRI/CL-NX model whereas macrophage numbers continue increasing between days 7-14 in the U-IRI model accompanied by a second wave of immune activation involving recruitment of neutrophils and T cells.

**Figure 4.**
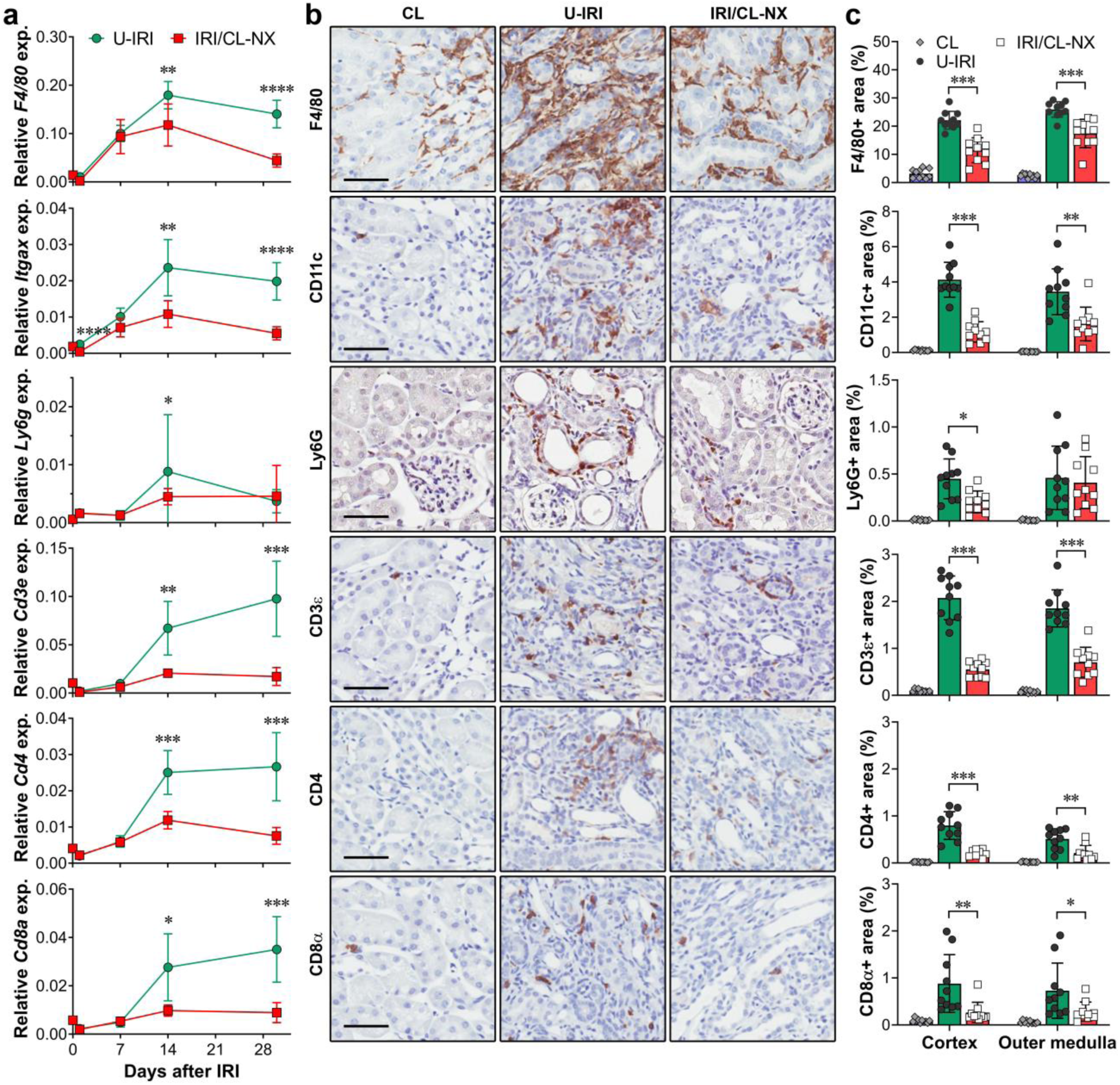
U-IRI promotes late immune cell accumulation. **a.** Quantitative RT-PCR analysis for *F4/80*, *Itgax*, *Ly6g*, *Cd3e*, *Cd4*, and *Cd8a* was performed on whole kidney RNA harvested on day 0, 1, 7, 14, and 30 after injury. n=10 kidneys/time point/group. Two-way ANOVA summarized in Supplementary Table 1. *p<0.05, **p<0.01, ***p<0.001 at each time point. **b.** Control (CL), U-IRI and IRI/CL-NX kidney sections on day 14 after IRI were immunostained with F4/80, CD11c, Ly6G, CD3ε, CD4, and CD8α. Representative images of kidney sections are shown at 40x magnification (20x magnification shown in Supplementary Fig. 4-9). Scale bars, 50 µm. **c.** F4/80-, CD11c-, Ly6G-, CD3ε-, CD4-, and CD8α-positive areas as in (b) were quantified. n=10 kidneys/group. Two-way ANOVA was summarized in Supplementary Table 2. *p<0.05, **p<0.01, ***p<0.001 in the indicated subgroup analyses.

### Chemokine-receptor interactions define the immune signature of the second wave of inflammation

To identify the pathways mediating the second wave of immune cell recruitment following IRI, we analyzed the gene expression of homing receptors and their known chemokine ligands in the integrated single cell dataset from both injury models. Macrophages predominantly expressed *Ccr2*, *Ccr5* and *Cx3cr1*; DCs expressed *Ccr2* and *Xcr1*; neutrophils expressed *Ccr1*, *Cxcr2*, and *Cxcr4*; and T cells expressed *Cxcr3* and *Cxcr6* (Supplementary Fig. 12). Correspondingly, homing chemokines for macrophages included the *Ccr2* ligands *Ccl2*, *Ccl7*, and *Ccl12* (expressed by macrophages themselves); the *Ccr5* ligands *Ccl3*, *Ccl4* and *Ccl8* (made by macrophages and/or neutrophils); and the *Cx3cr1* ligand *Cx3cl1* (expressed at low levels by Injured PT and myofibroblasts) (Supplementary Figs. 13, 14). Chemokines for neutrophil recruitment and activation included the *Ccr1* ligands *Ccl3*, *Ccl5*, *Ccl8* and *Ccl9* (expressed by neutrophils, Cd8a+ T cells, macrophages and dendritic cells, respectively); the *Cxcr2* ligand *Cxcl2* (expressed by neutrophils); and the *Cxcr4* ligand *Cxcl12* (expressed by myofibroblasts at a low level). Finally, T cell recruitment and activation appears to predominantly depend on the *Cxcr6* ligand *Cxcl16* (expressed by macrophages) since the other potential T cell recruitment chemokines were expressed either at low levels (*Cxcl9* and *Cxcl10*) or not detected (*Cxcl11*) (Supplementary Figs. 13, 14). Together, the data suggests that infiltrating, M1, and M2 macrophages are the predominant cells to express immune-recruiting chemokines in the injured kidneys.

To better understand how macrophages respond to kidney atrophy, we identified the differentially expressed genes (DEG) of infiltrating, M1, and M2 macrophages between the U-IRI and IRI/CL-NX kidneys. On day 14, we found a set of chemokines including *Ccl2*, *Ccl7*, *Ccl8*, and *Ccl12*, that were significantly upregulated in those macrophages from the U-IRI kidneys as compared to the IRI/CL-NX kidneys (Fig. 5a-c). The upregulation of these chemokines by day 14 in U-IRI kidneys correlates with the second wave of immune cells observed on days 14 and 30 in this model (Fig. 4a). Quantitative PCR for the macrophage receptor *Ccr2* and its activating chemokines *Ccl12*, *Ccl7* and *Ccl2*^29^; the neutrophil receptor *Ccr1* and its activating chemokine *Ccl5*; and the T cell receptor *Cxcr6* and its activating chemokine *Cxcl16* from whole kidney mRNA confirmed the significant increase of these ligand-receptor pairs between days 14 and 30 in the U-IRI injury model (Fig. 5e and Supplementary Fig. 15).

**Figure 5.**
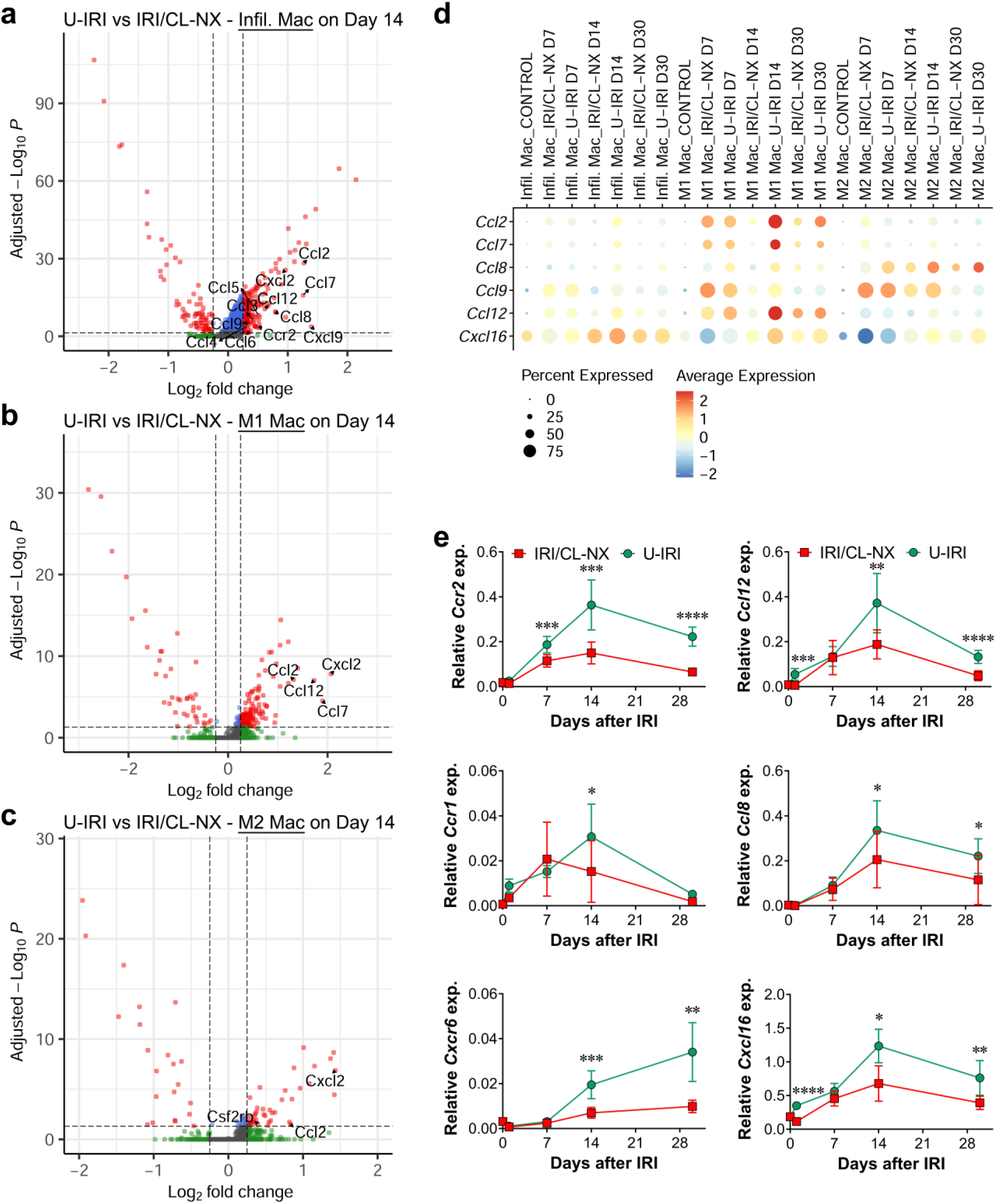
U-IRI promotes chemokine expression at the late stage of IRI. **a-c.** Volcano plots demonstrating differential gene expression in U-IRI compared to IRI/CL-NX derived infiltrating macrophages (**a**), M1 macrophages (**b**), and M2 macrophages (**c**) on day 14 after injury. **d.** The distribution and relative expression of chemokines were visualized in the dot plot. **e.** Quantitative RT-PCR analysis for the indicated chemokines and chemokine receptors was performed on whole kidney RNA harvested on day 0, 1, 7, 14, and 30 after injury. n=10 kidneys/time point/group. Two-way ANOVA summarized in Supplementary Table 1. *p<0.05, **p<0.01, ***p<0.001, ****p<0.0001 at each time point.

To identify which of these ligand-receptor interactions resulted in functional immune cell responses, we utilized the NicheNet ligand-receptor-target algorithm developed by Saeys and colleagues^30, 31^. Since the chemokines were predominantly expressed by macrophages at 14 days (Supplemental Fig. 13), we focused on immune cell-secreted ligands. This analysis revealed extensive chemokine-receptor interactions between infiltrating, M1, and M2 macrophage secreted ligands and their receptors on neutrophils (*Ccr1*, *Cxcr2*, and *Cxcr4*), Cd4+ Th/Treg, and Cd8a+ Tc/NKT (*Ccr2*, *Ccr5*, *Cxcr3*, and *Cxcr6*) (Fig. 6a,b and Supplementary Figs. 16, 17). For example, *Ccl3*, *Ccl4*, and *Ccl8* that were significantly upregulated in M1 and M2 macrophages on day 14 after U-IRI were predicted to interact with Ccr1-expressing cells such as PMN #1. As a result, the pathways involved in leukocyte (neutrophils, monocytes, and lymphocytes) chemotaxis were significantly enriched in the infiltrating and M1 macrophages (Fig. 6c). The functional role of these macrophage-secreted chemokines in recruiting the second wave of T cells and PMNs was supported by numerous examples of both CD3ε+ T cells and Ly6G+ PMNs found adjacent to F4/80+ macrophages in kidneys 14 days after U-IRI. Together, macrophages that persist beyond day 7 in U-IRI kidneys are selectively activated to express high levels of chemokines, which are predicted to promote a second wave of inflammatory neutrophil and T cell recruitment, resulting in a local nidus for extensive inter-and intra-immune cell cross-talk^32, 33^.

**Figure 6.**
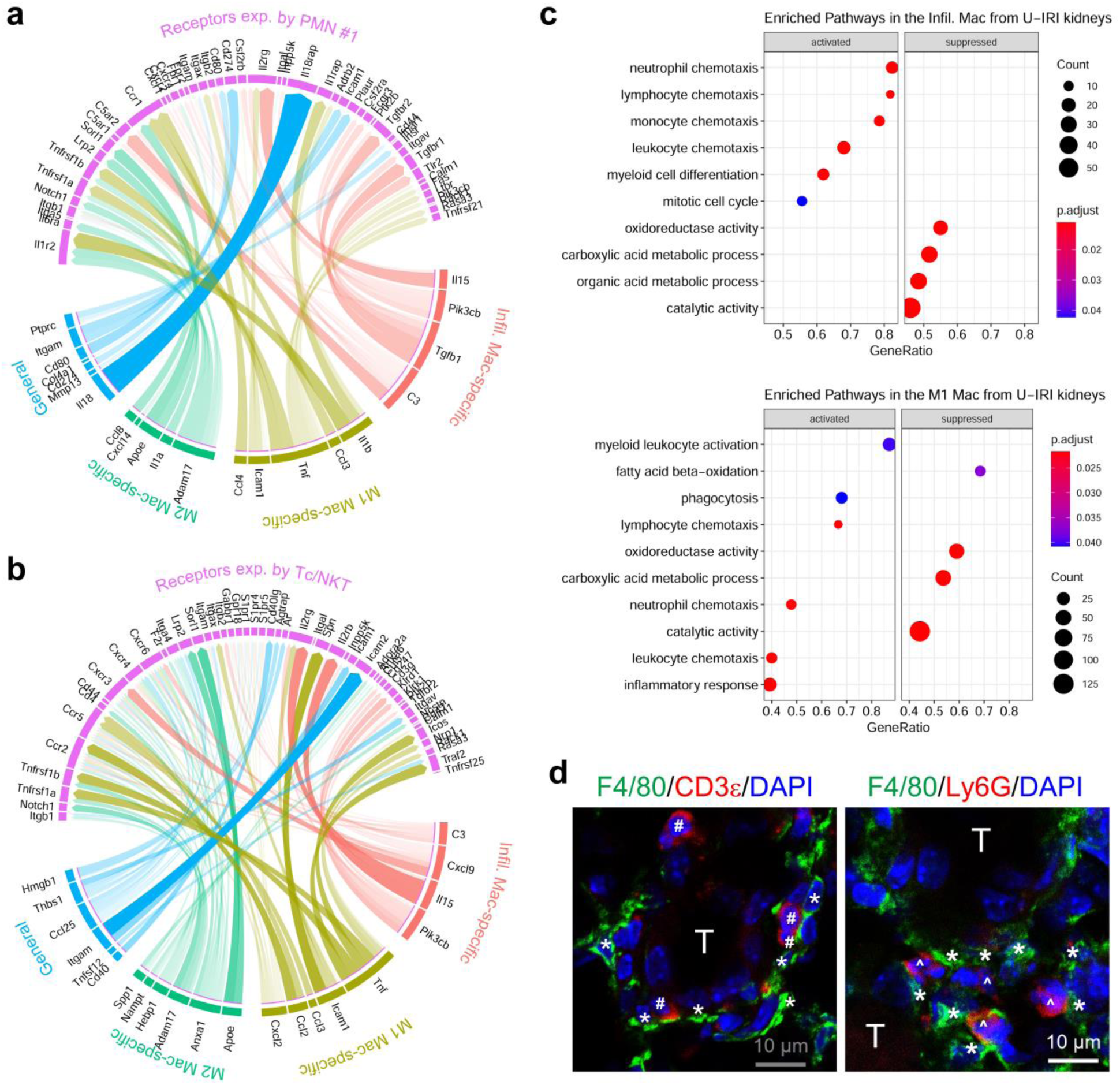
U-IRI promotes late immune cell recruitment. **a** and **b.** Based on DEG between U-IRI and IRI/CL-NX kidneys on day 14 after injury, the potential ligands expressed by the infiltrating, M1, and M2 macrophages were linked to their corresponding receptors based on the potential target genes (as shown in Supplementary Fig. 16) for PMN #1 (**a**) and Tc/NKT cells (**b**) and visualized by a chord diagram. **c.** The top relevant enriched GO terms for infiltrating (top panel) and M1 (lower panel) macrophages were visualized in the dot plots. **d.** Representative images of IF staining for F4/80 (green), CD3ε (red on the left panel), and Ly6G (red on the right panel) on the kidney sections 14 days after U-IRI. T, tubules; *, macrophahges; #, T cells; ^, PMNs. Scale bars: 10 μm.

### Late immune activation promotes tubule oxidative stress and secondary injury

To understand the impact of this second wave of immune activation on tubule injury and atrophy, we determined the DEG of PMNs, Tc/NKT, and Th/Tregs between the U-IRI and IRI/CL-NX kidneys. On day 14, we found that PMNs in the U-IRI kidney significantly upregulated inflammatory gene expression including *Il1b*, *Il1f9*, *Tnf*, *Tnfaip3*, *Ifitm1*, etc. and *Lcn2* (Fig. 7a,b). Cd8+ Tc/NKT cells in the U-IRI kidney expressed significantly higher levels of T cell activation genes such as *Tyrobp*, *Klre1*, *Serpinb9*, *Ripor2*, *Fasl*, *Xbp1*, *Zfp36l2*, etc. (Fig. 7c). Cd4+ Th/Treg cells in the U-IRI kidneys also increased the expression of T cell activation genes such as *Fyn*, *Gpr183*, *Laptm5*, *Lax1*, *Lfng*, *Itk*, *Trp53*, *Zfp36l1*, etc. (Supplemental Fig. 18a). The upregulation of these genes is predicted to activate pathways involved in production of interleukin 1, 6, and 12, interferon γ, nitric oxide, and superoxide anion in the PMNs (Fig. 7d,e), cell killing and regulation of apoptosis in the Tc/NKT cells (Fig. 7f), and T cell activation and antigen recognition in the Th/Treg cells (Supplemental Fig. 18a,c) on day 14 after U-IRI. Quantitative PCR analysis confirmed that the mRNA expression levels of *Il1b*, *Tnf*, *Fasl*, *Ltb*, and *Cd40lg* were increased to a greater degree at 14 and/or 30 days in the U-IRI kidneys as compared to those in the IRI/CL-NX kidneys (Fig. 7g and Supplemental Fig. 18b), consistent with the increase in neutrophil and T cell number seen during the second wave of immune activation (Supplementary Tables 3 and 4). Ligand-receptor-target analyses reveals that PMNs, Tc/NKT, and Th/Treg are projected to induce PT cells to increase expression of several genes including *Vcam1*, *Spp1*, *Csf1*, and *Cp* (ferroxidase) that have been previously implicated as injury markers (Fig. 7h and Supplementary Figs. 19-21).

**Figure 7.**
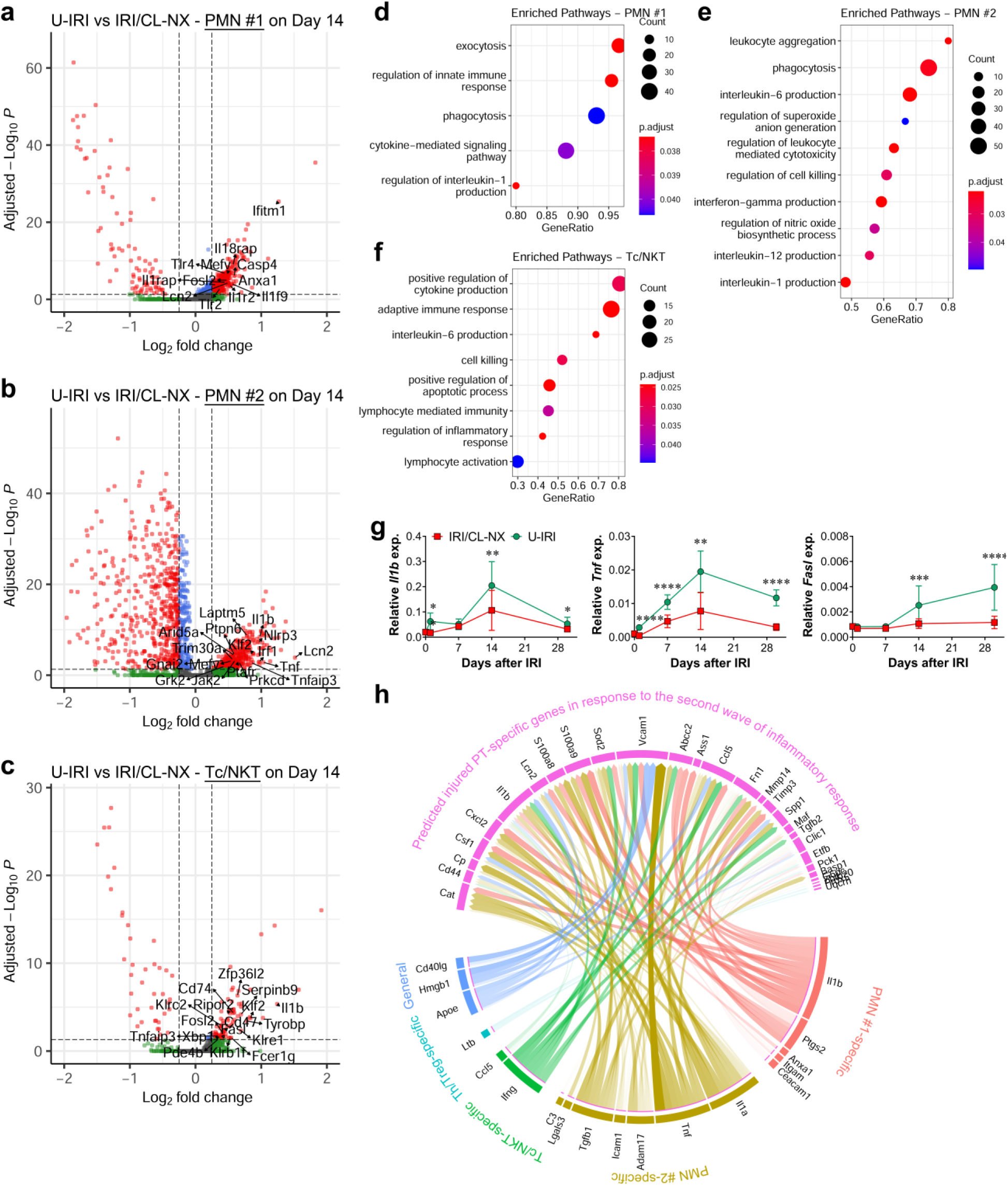
U-IRI promotes late inflammation. **a-c.** Volcano plots demonstrating differential gene expression in U-IRI compared to IRI/CL-NX derived PMN #1 (**a**), PMN #2 (**b**), and Cd8a+ Tc/NKT cells (**c**) on day 14 after injury. **d**-**f.** Based on DEG between U-IRI and IRI/CL-NX kidneys on day 14 after injury, the top relevant enriched GO terms for PMN #1 (**d**), PMN #2 (**e**), and Cd8a+ Tc/NKT cells (**f**) were visualized in the dot plots. **g.** Quantitative RT-PCR analysis for indicated genes was performed on whole kidney RNA harvested on day 0, 1, 7, 14, and 30 after injury. n=10 kidneys/time point/group. Two-way ANOVA summarized in Supplementary Table 1. *p<0.05, **p<0.01, ***p<0.001, ****p<0.0001 at each time point. **h.** Based on DEG between U-IRI and IRI/CL-NX kidneys on day 14 after injury, the potential ligands expressed by the PMNs, Cd8a+ Tc/NKT, and Cd4+ Th/Treg cells were linked to their corresponding potential target genes for the injured PT cells and visualized by a chord diagram

To determine if the second wave of immune activation induced the predicted injury response, we analyzed the DEG of PT cells between the U-IRI and IRI/CL-NX kidneys. On day 14, we found that proximal tubule cells from U-IRI kidneys expressed significantly higher levels of several injury markers including *Vcam1*, *Csf1*, and *Spp1*, as well as multiple major histocompatibility complex (MHC) class II genes (*H2-Aa*, *H2-Ab1*, *H2-Eb1*, and *Cd74*) and class I genes (*H2-D1* and *H2-K1*), while downregulating anti-oxidative stress genes (*Gpx1*, *Gpx3*, *Gpx4*, *Gsta2*, *Gstm1*, *Gstp1*, *Gatm*, and *Iscu*) as well as the PT differentiation marker (*Lrp2*)(Fig. 8a,b and Supplemental Fig. 22a,b). This shift of gene expression by tubule cells from anti-oxidative stress genes induced following IRI/CL-NX to expression of MHC genes and injury markers after U-IRI was observed beginning on day 7 and extending to day 30 and seen in most nephron segments (PT, TAL, DCT/CNT, and CD; Fig. 8c). The increase of MHC class I expression is associated with the increase of *Vcam1* expression in the injured PT and *Lcn2* expression in the CD-PC cells in the U-IRI kidneys (Fig. 8c). Consistent with the single cell data, quantitative PCR analysis confirmed that mRNA expression levels of *H2-Aa*, *H2-Ab1*, and *Cd74* were markedly upregulated in U-IRI kidneys on days 14 and 30 after injury (Fig. 8d and Supplemental Fig. 22c). The expression kinetics of *H2-Aa*, *H2-Ab1*, and *Cd74* highly correlated with the expression of T cell markers *Cd3e*, *Cd4*, and *Cd8a* as well as *Tnf* and *Ltb* (Supplementary Tables 4,5). Taken together, the second wave immune response correlated with increased antigen presentation, T cell activation, and T cell-mediated cytotoxicity in the injured PT after U-IRI (Fig. 8e).

**Figure 8.**
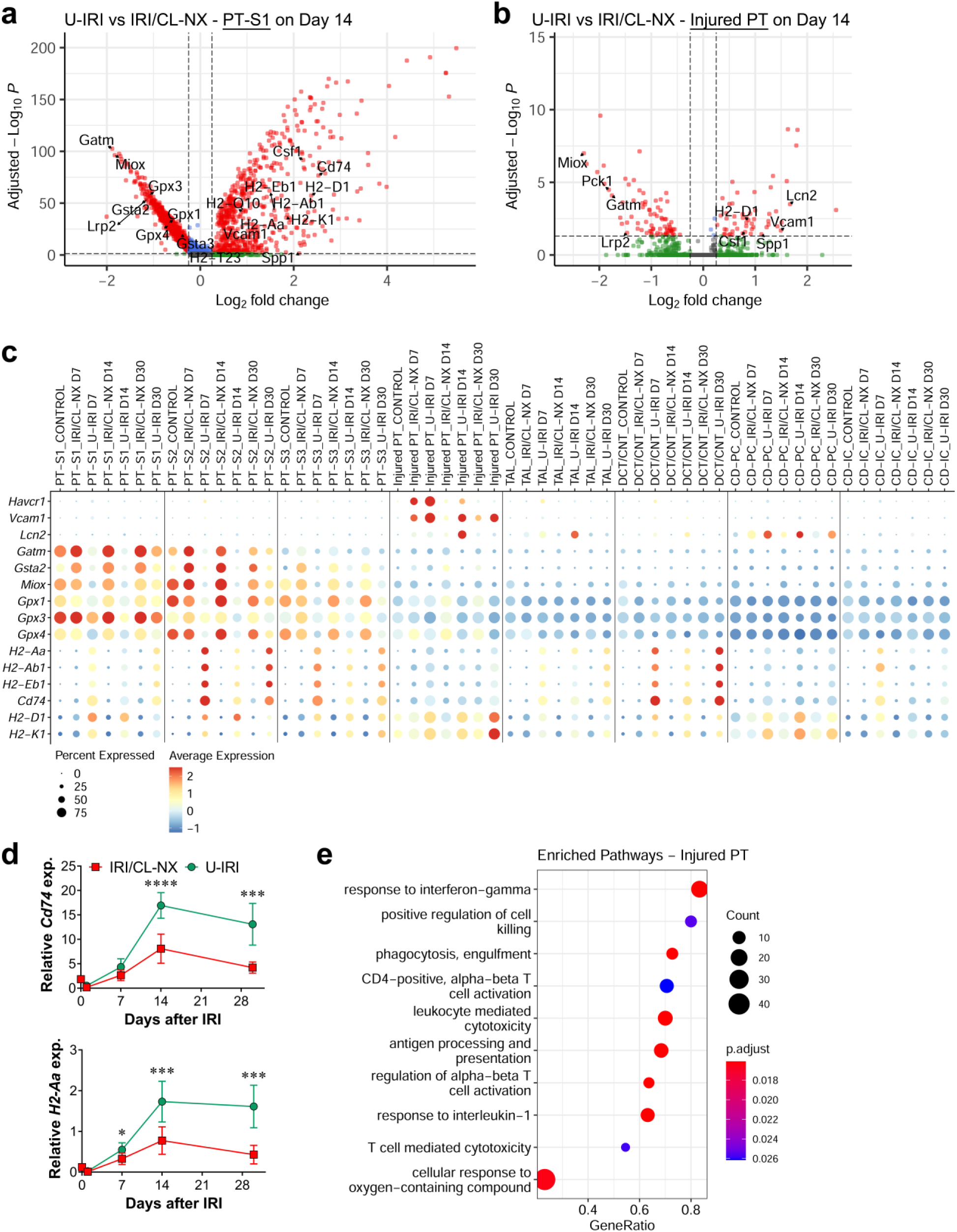
Upregulation of MHC class I and II genes in response to inflammation-induced tubular stress. **a** and **b.** Volcano plots demonstrating the differential average gene expression in cells from the PT-S1 (**a**) and injured PT (**b**) 14 days after U-IRI compared to IRI/CL-NX. **c.** The distribution and relative expression of the injury marker (*Havcr1*, *Vcam1*, and *Lcn2*), anti-oxidative stress and detoxification genes (*Gatm*, *Gsta2*, *Miox*, *Gpx1*, *Gpx3*, and *Gpx4*), and the major histocompatibility complex class II (*H2-Aa*, *H2-Ab1*, *H2-Eb1*, and *Cd74*) and class I (*H2-D1* and *H2-K1*) are visualized in a dot plot. **d.** Quantitative RT-PCR analysis for indicated genes was performed on whole kidney RNA harvested on day 0, 1, 7, 14, and 30 after injury. n=10 kidneys/time point/group. Two-way ANOVA was summarized in Supplementary Table 1. *p<0.05, ***p<0.001, ****p<0.0001 at each time point. **e.** Based on DEG between U-IRI and IRI/CL-NX kidneys on day 14 after injury, the top relevant enriched GO terms for injured PT were visualized in the dot plots.

### Depletion of T cells and neutrophils together attenuates kidney atrophy and loss of nephrons

To understand whether the second wave of inflammatory T cells and neutrophils promotes kidney tubule atrophy, we depleted either neutrophils, T cells, or both beginning on day 5 following U-IRI and analyzed the phenotype of the injured kidney on day 30. We found that depletion of neutrophils and T cells together in the U-IRI model of injury (Fig. 9a-c), but not either cell type individually (Supplemental Figs. 23, 24), improved kidney-to-body weight ratio by 20% following U-IRI (Fig. 9d). Quantitative analysis of LTL staining revealed an 80% increase in the absolute area of proximal tubule brush border following dual T cell and neutrophil depletion, reflecting by a 50% increase in the percent of the total cross-sectional area comprised by PT brush border (Fig. 9e,f). Quantitative PCR analysis confirmed that the mRNA expression levels of differentiated proximal tubular transporters *Lrp2* and *Scl34a1* increased 1.5- and 1.7-fold, respectively, following depletion of both T cells and neutrophils in the U-IRI kidneys (Fig. 9g). Together, these results suggest that the second wave of activated T cells and neutrophils promote loss of tubule cell differentiation and reduced nephron mass following U-IRI.

**Figure 9.**
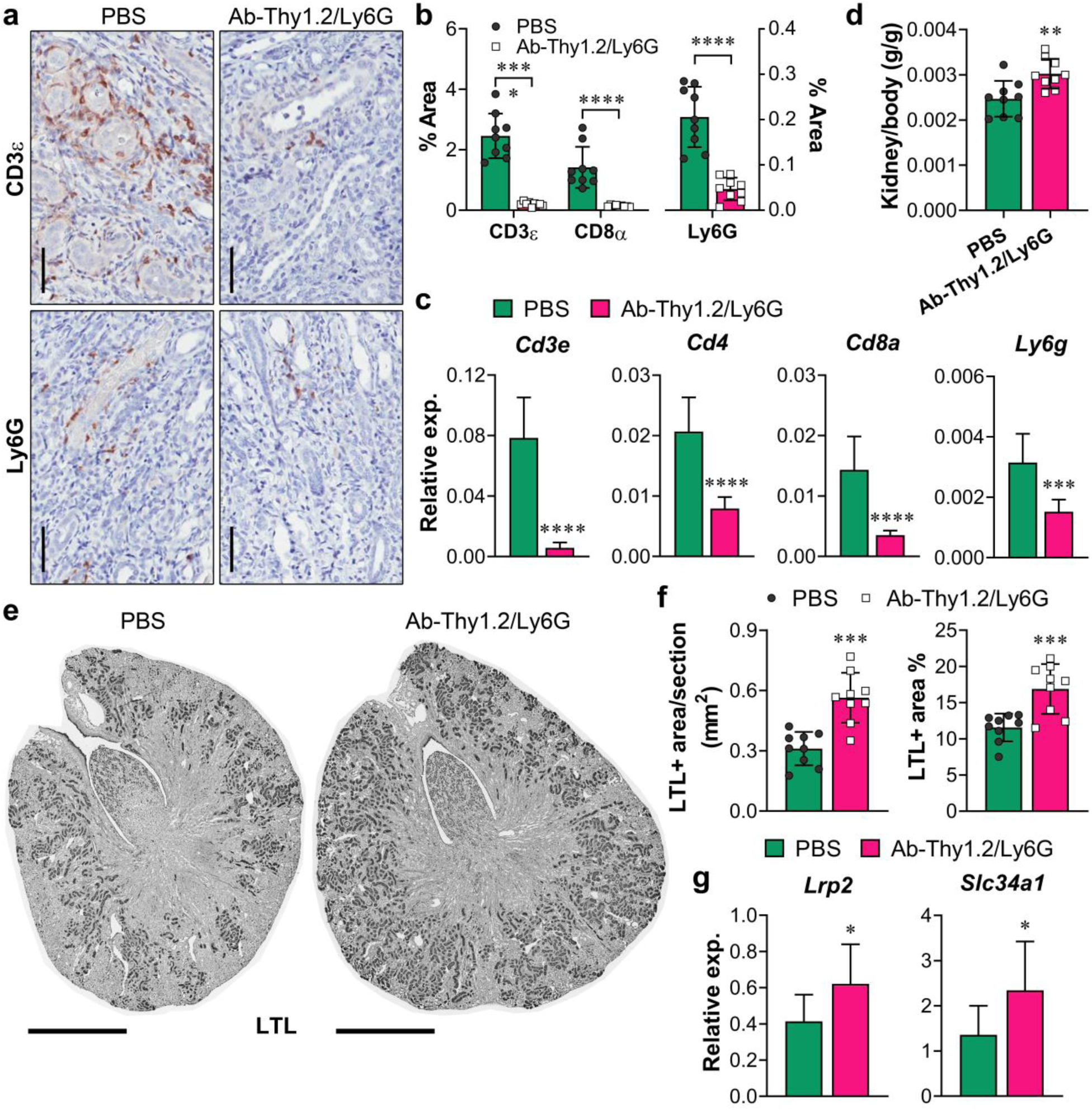
Dual depletion of T cells and neutrophils attenuates kidney tubule atrophy. WT mice were treated as described in Methods with either PBS or a combination of antibodies (Ab)-against Thy1.2 and Ly6G beginning 5 days after U-IRI and sacrificed on day 30. **a.** The IRI kidney sections were immunostained with CD3ε and Ly6G. Representative images of kidney sections are shown. Scale bars, 25 µm. **b.** CD3ε-, CD8α-, and Ly6G-positive areas were quantified using Image J. n=9 kidneys/group. ****p<0.0001. **c.** Quantitative RT-PCR analysis for *Cd3e*, *Cd4*, *Cd8a*, and *Ly6g* was performed on whole kidney RNA on day 30 after U-IRI ± dual T cell/neutrophil depletion. n=9 kidneys/group. ***p<0.001, ****p<0.0001. **d.** Kidney-to-body weight ratios on day 30 following U-IRI ± dual T cell/neutrophil depletion. n=9 kidneys/group. **p<0.01. **e.** Kidney sections from day 30 after U-IRI ± dual T cell/neutrophil depletion were immunostained with LTL (dark grey). Representative images are shown. Scale bars, 1 mm. **f.** LTL-positive area was quantified from the entire section (left panel) and as a percentage of the section (right panel). n=9 kidneys/group. ***p<0.001. **g.** Quantitative RT-PCR analysis for *Lrp2* and *Slc34a1* was performed on whole kidney RNA from U-IRI mice ± dual T cell/neutrophil depletion. n=9 kidneys/group. *p<0.05.

### Accumulation of T cells and neutrophils is negatively associated with GFR recovery in patients with AKI

Our animal model findings prompted us to investigate the relevance of T cell and neutrophil accumulation for recovery of kidney function in patients with AKI. To assess this, ten kidney biopsies from patients with AKI, an estimated GFR (eGFR) >60 mL/min/1.73m^2^ prior to AKI episode, and no history or pathologic indication of glomerulonephritis (GN), diabetic kidney disease (DKD), or acute interstitial nephritis (AIN) were selected from Yale kidney biopsy biorepository (Supplementary Tables 6 and 7)^34, 35^. For each of these patients, eGFR declined by 61 (36, 71) mL/min at the time of AKI from pre-AKI level with partial or full amounts of recovery 6 months after AKI (Fig. 10a and Supplementary Tables 6 and 7). We quantified the number of T cells and neutrophils in sections from each of these kidney biopsies, and compared to those seen in kidney biopsies from healthy living kidney donors^36^. As expected, very few T cells or neutrophils were found in kidneys from living donors, whereas T cells and neutrophils were easily detected in the kidney interstitium of some patients at the time of biopsy for AKI (Fig. 10b). Nonparametric Spearman correlation analysis of the number of T cells or neutrophils (quantified as a percentage of total cells identified in the cortex) with either the absolute or relative increase in eGFR at 6 months after AKI revealed a strong negative correlation between inflammatory cell number and eGFR recovery following AKI (Fig. 10c,d).

**Figure 10.**
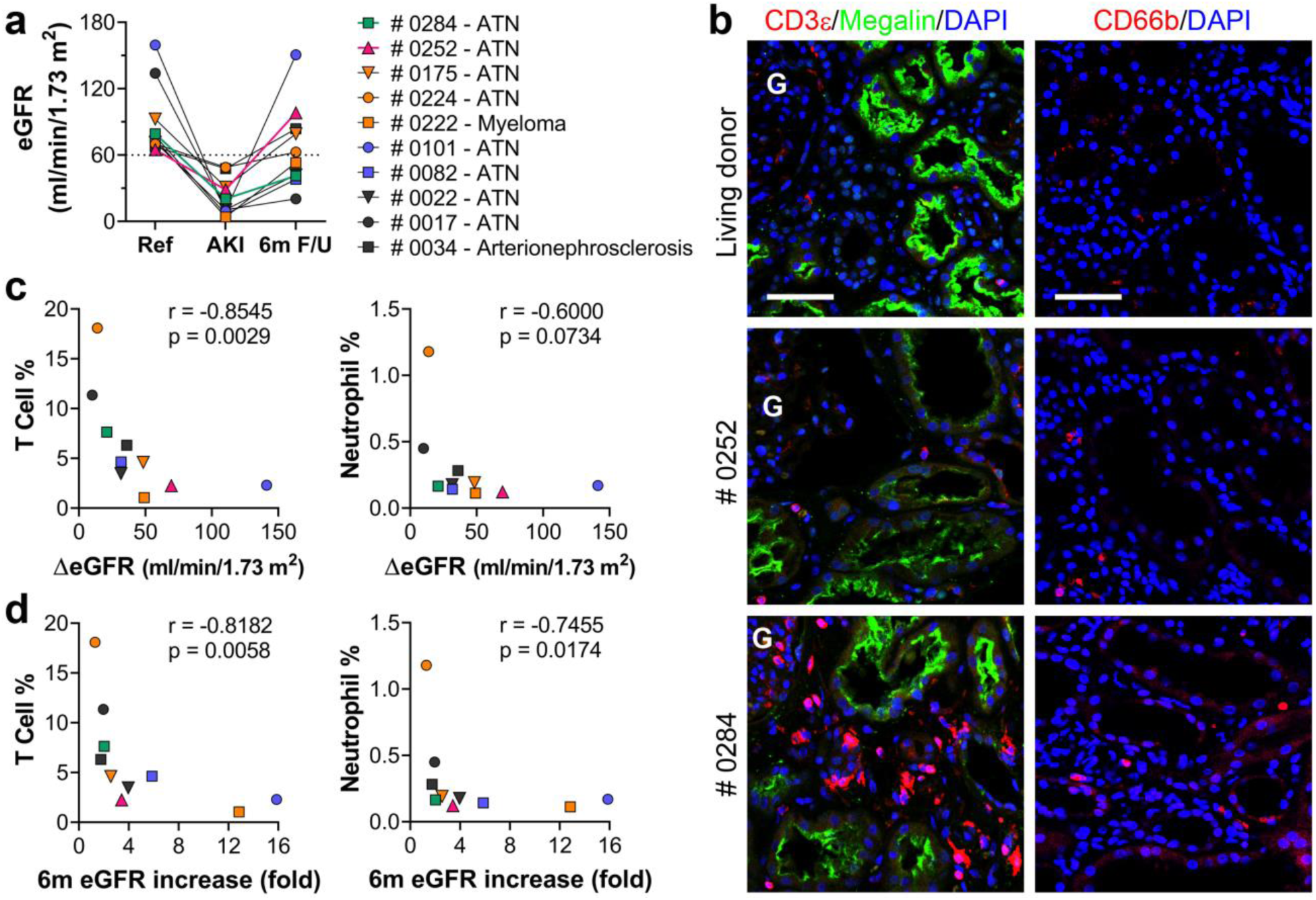
Accumulation of T cells and neutrophils negatively associates with GFR recovery in patients with AKI. **a.** The estimated GFR (eGFR) was determined at reference, biopsy (AKI), and 6-month follow-up (6m F/U). **b.** Biopsy sections were immunofluorescence-stained with CD3ε and megalin (left panel) or CD66b (right panel). Representative images of biopsies from a patient with full recovery (Case #0252, 150% recovery of GFR, magenta triangle in (**a**)), low-recovery (Case #0284, 52% recovery of GFR, green square in (**a**)), and a healthy living donor are shown. Scale bars, 50 µm. **c.** Correlation between absolute eGFR increase within 6 months after biopsy and T cell infiltrate (%) or neutrophil infiltrate (%) at the time of biopsy was determined by nonparametric Spearman correlation coefficient r. **d.** Correlation between relative eGFR increase (fold change) within 6 months after biopsy and T cell infiltrate (%) or neutrophil infiltrate (%) at the time of biopsy was determined by nonparametric Spearman correlation coefficient r.

## DISCUSSION

Kidney injury has been extensively investigated using mouse models of either bilateral IRI or IRI/CL-NX to develop our current mechanistic understanding of tubule injury and repair including mechanisms of cell death, the innate immune response to injury, proliferation of surviving tubular cells to replace those that are lost, clearance of casts and restoration of GFR^37–40^. Recently, our group and others have shown that performing U-IRI with the contralateral kidney intact leads to a different outcome in which fibrosis and kidney atrophy predominate rather than tubule repair and restoration of function^7, 20, 21, 37^.

In this study, we show that the initial macrophage, DC, neutrophil and T cell response to ischemia/reperfusion injury is similar during the first 7 days after injury in the two models. However, after 7 days the responses markedly diverge with a surge in T cells and neutrophils observed in the U-IRI kidneys as compared to IRI/CL-NX kidneys. The divergence becomes detectable on day 7 with more sustained *Kim1* expression by the proximal tubule and de novo upregulation of *Vcam1*, and ends with extensive kidney atrophy by day 30. As previously reported by our group and others, the decreased kidney size following U-IRI is accompanied by excessive macrophage persistence and a transition to a *Pdgfb+* and *Tgfb1+* pro-fibrotic expression profile^21, 25^. However, quantitative analysis of the remaining tubule area relative to total kidney area suggests that the major phenotypic change underlying kidney atrophy after U-IRI is tubule loss rather than increased fibrosis (Fig. 1). In fact, the transition of macrophages to a profibrotic phenotype was seen in both models of IRI (Supplementary Fig. 5b-f), consistent with evidence that even “reparative” models of kidney injury such as IRI/CL-NX or bilateral IRI lead to increased kidney fibrosis^29, 41^.

Similar to the results of Liu and colleagues who studied the slowly progressive kidney atrophy and fibrosis seen following bilateral IRI^41^, we find that the accelerated renal atrophy following U-IRI correlates with sustained tubular injury and inflammatory cell activation. By performing single cell sequencing and cell-cell ligand-receptor-target interaction analyses, we now demonstrate that this delayed increase in inflammation in the U-IRI kidneys correlates with the unique chemokine (*Ccl2*, *Ccl7*, *Ccl8*, *Ccl12*, and *Cxcl16*) expression by macrophages, that in turn can recruit more macrophages^21^ as well as a second wave of infiltrating proinflammatory neutrophils and T cells between 7-30 days after injury.

Initial neutrophil recruitment and transmigration peaks at approximately 24 hours after IRI and promotes reactive oxygen species (ROS) production, activation of resident mononuclear phagocytes, and differentiation of recruited monocytes into proinflammatory macrophages, all of which contribute to the early components of reperfusion injury^6^. In models of injury where repair pathways predominate, CD169+ macrophages limit subsequent neutrophil infiltration in the kidney by downregulating intercellular adhesion molecule-1 (ICAM-1) expression on vascular endothelial cells^42^, and neutrophil numbers in the interstitium dramatically decrease after day 3, corresponding to the period when pro-reparative macrophages become prominent. In contrast, intrarenal T cells are not typically seen in large numbers in the first week after IRI, but have been identified in the late stages of IRI where they can promote upregulation of proinflammatory cytokine expression including *Il1b*, *Il6*, *Tnf,* and *Ifng*^43–45^. We now show that in a model where atrophy predominates, this late neutrophil and T cell infiltrate correlates with sustained tubular cell dedifferentiation and the increased expression of multiple homing chemokines by macrophages as well as the infiltrating T cells and neutrophils themselves.

Infiltrating T cells can be activated through antigen-independent mechanisms by inflammatory cytokines and reactive oxygen intermediates as well as through interaction with antigen-presenting cells (APCs). Our results illustrate that in response to the late cell stress induced by inflammatory cytokines, proximal tubular epithelial cells in U-IRI kidneys upregulated MHC class II (*H2-Aa*, *H2-Ab1*, *H2-Eb1*, and *Cd74*) and class I (*H2-D1* and *H2-K1*) genes, and thus become antigen-presenting cells capable of T cell receptor (TCR) engagement with both Cd4+ and Cd8a+ T cell activation^46^. Consistent with this, *Lat*, which encodes the LAT adaptor protein component of the TCR activation complex, was significantly upregulated in intrarenal Cd4+ T cells (Supplemental Fig. 18a), and Cd8+ T cells were activated to express cytotoxins such as perforin, granzymes, and FAS ligand (Fig. 7c) following U-IRI. Interestingly, neutrophils that increase inflammatory cytokine and nitric oxide production have also been shown to efficiently interact with and prime naïve T cells directly or indirectly in vivo^47^, suggesting that the simultaneous recruitment of both cell types beginning by day 7 after U-IRI may be important in promoting the secondary tubule injury as indicated by proximal tubule-*Vcam1* expression and collecting duct-*Lcn2* expression (Figs. 2, 7h, and Supplemental Fig. 20). This immune-mediated tubule injury may differ from the initial sterile ischemic injury as *Havcr1* expression gradually decreased despite the tubular atrophy, whereas other injury markers such as *Vcam1* are upregulated only at this later stage of AKI to CKD transition. Consistent with this, a recent study by Kirita et al identified a distinct proinflammatory proximal tubule cell (FR-PTCs) state that fails to repair at the late stage of bilateral IRI^4^. These FR-PTCs were shown to increase *Vcam1* expression and were involved in positive regulation of lymphocyte activation^4^.

The question that our single cell sequencing data raised is whether or not the late influx of T cells and neutrophils, occurring after proreparative macrophage activation has waned, is responsible for the late tubule dedifferentiation, sustained injury and atrophy seen following U-IRI. Experiments in which we selectively depleted either T cells alone or neutrophils alone failed to reduce the progressive kidney and tubule atrophy (Supplemental Figs. 23 and 24), but when we depleted both T cells and neutrophils together there was a significant reduction in the degree of tubule atrophy following U-IRI with preservation of proximal tubule brush border and differentiation (Fig. 9). However, depletion of T cells and neutrophils did not fully prevent kidney atrophy when compared to the contralateral kidneys and control kidneys. This partial effect may be due to our inability to fully eliminate T cell and neutrophil infiltration (approximately 80% reduction based on immunostaining), because accumulation of macrophages and DCs was not prevented by the administration of the anti-Thy1.2 and anti-Ly6G antibodies (data not shown).

Alternatively, the partial response to cell depletion reflect that dichotomy of T cell responses to kidney injury^48^. It has been shown that CCR5 was upregulated in the CD3+ infiltrating T cells, and blockade of CCR5 using a neutralizing antibody in mice protected renal function after U-IRI^49^. In addition, the store-operated calcium entry (SOCE) channel, Orai1, participates in the activation of Th17 cells and influences renal injury^50^. Blockage of SOCE in rats attenuated Th17 cell activation, inflammation, and severity of AKI following IRI or intramuscular glycerol injection^50^. In contrast, regulatory T (Treg) cells (TCRβ+,CD4+,CD25+,Foxp3+) that infiltrated IRI kidneys during the healing process have been shown to promote kidney repair, likely by modulating proinflammatory cytokine production of other T cell subsets^51^.

Finally, we show that patients who underwent a renal biopsy at the time of AKI exhibit a strong negative correlation between the number of infiltrating neutrophils and T cells and the subsequent recovery of estimated GFR (Fig. 10). While acknowledging the limitations of eGFR in patients with AKI, our findings are consistent with studies showing large numbers of terminally differentiated CD4+ T helper and CD8+ cytotoxic T cells in patients with ESRD,^52^ and raise the ultimate question of whether or not these data may provide new targets for slowing or even preventing the progression from AKI to CKD in humans. By analyzing the chemokine-receptor interactions among cell types in the injured kidneys, we identified several potentially important signaling interactions. For instance, interactions were identified between infiltrating macrophages and dendritic cells through CCL2-CCR2 signaling, and between macrophages, DC and T cells via CXCL16-CXCR6 signaling. We recently reported that blocking CCL2-CCR2 signaling by knock-out of *Ccr2* or use of the CCR2 antagonist RS102895 significantly reduced the number of macrophages, DCs and T cells following U-IRI, resulting in a reduction in ECM deposition and kidney injury marker expression^21^. However, the decrease in inflammatory cell numbers was modest in that study and there was no protection against kidney atrophy. It is perhaps not surprising that blocking a single pathway was ineffective given the complexity and redundance built into inflammatory activation pathways, with alternative recruitment and activation signals such as CCL8/CCL6/CCL9/CCL5-CCR1 signaling identified between these same cell populations. Ultimately effective strategies are likely to require targeting multiple inflammatory pathways and/or the initiation process that is likely mediated by the injured tubular and endothelial cells.

In summary, our results show that cross-talk between unrepaired tubular cells and macrophages, occurring after the initial reperfusion injury and associated first wave of neutrophil and macrophage infiltration has waned, promotes a second wave of inflammatory cell infiltration and activation that underlies the sustained tubule injury, dedifferentiation and atrophy seen after unilateral IRI. Macrophages appear to play a central role in recruiting the other immune cells, while the injured tubular cells themselves are primed to promote T cell activation. The combined presence of neutrophils and activated T cells appears to be required for the subsequent tubule loss and kidney atrophy, suggesting that blocking their recruitment or activation may be a logical approach for suppressing AKI-to-CKD transition.

## LIMITATIONS

There are some limitations in our studies. Despite our effort to optimize the dissociation method, the intrarenal cells including tubular epithelial cells, podocytes, endothelial cells, myofibroblast, immune cells might not be evenly dissociated from the injured kidneys as compared to controls. Hence, although the gene expression profiles and cell clustering should be highly accurate, the cell numbers of those populations in the scRNA-seq analyses may be impacted by these differences in cell disassociation.

## METHODS

### Animal Surgery and Experimental Protocol

All animal protocols were approved by the Yale University Animal Care and Use Committee. C57BL/6 (Envigo) wild-type mice (age 9-11 weeks) were used in this work. All mice were maintained on a 12-hour light and 12-hour dark cycle and with free access to standard food and water before and after surgery. Due to the substantial difference in susceptibility to IRI injury between male and female mice^53^, male mice were exclusively used to reduce total numbers of mice required for statistical analysis. Before surgery, all mice were subjected to anesthesia by intraperitoneal injection with ketamine (100 mg/kg) and xylazine (10 mg/kg) on a 37 °C warming pad. To establish the unilateral ischemia/reperfusion injury (U-IRI) model, the abdomen was opened, and warm renal ischemia was induced using a nontraumatic microaneurysm clip (FST Micro Clamps) on the left renal pedicle for 27 min, leaving the right kidney intact. To establish the unilateral IRI with contralateral nephrectomy (IRI/CL-NX) model, the right kidney was surgically removed at the time of left kidney ischemia. During surgery, all mice received intraperitoneal phosphate-buffered saline (PBS) and buprenorphine (0.1 mg/kg) to avoid dehydration and postoperative pain, respectively. The mice were sacrificed on day 1, 7, 14 and 30 after U-IRI or IRI/CL-NX (n=10 mice/end point). Baseline control mice were sacrificed and denoted as day 0 for the injury (n=10 mice) or at day 30 and designated as age-matched controls (n=10). To establish the nephrectomy (NX) alone model, the right kidney was surgically removed, and the mice were sacrificed on day 30 after NX (n = 8). Additional mice were sacrificed on day 7, 14, and 30 after U-IRI or IRI/CL-NX (n = 2 mice at each time point/model and 2 healthy control mice) for kidney cell isolation followed by single cell RNA sequencing. Blood and tissue samples were obtained at the indicated times after the surgery. Serum creatinine and blood urea nitrogen (BUN) were measured at the Yale George M. O’Brien Kidney Center.

### In vivo depletion of T cells and neutrophils

To deplete T cells and neutrophils, wild-type mice were i.p. injected with 200 µg anti-mouse Thy1.2 (CD90.2) antibody (clone 30H12, BioXCell) once in three days and 200 µg anti-mouse Ly6G (clone 1A8, BioXCell) (n=9 mice) every other day or PBS control (n=9 mice) starting on day 5 after U-IRI. The depletion efficacy was confirmed at the kidney level using immunohistochemistry against CD3ε, CD4, CD8α, and Ly6G, respectively, and qPCR analysis for *Cd3e*, *Cd4*, *Cd8a*, and *Ly6g*, respectively, on the whole kidney RNA at the end point (day 30 after U-IRI).

### ELISA of Serum KIM-1 and NGAL Levels

Serum KIM1 and NGAL concentrations were measured using mouse TIM-1/KIM-1/HAVCR and Lipocalin-2/NGAL quantikine ELISA kits (R&D Systems) according to the manufacturer’s instructions.

### Preparation of Single Cell Suspension

Euthanized mice were perfused with chilled 3× PBS (10 mL) via the left ventricle. Kidneys were harvested, minced into approximately 1 mm^3^ cubes, and digested using Liberase™ (100 µg/mL) and DNase I (10 µg/mL) (Roche Diagnostics) for 25 min at 37 °C. Reaction was deactivated by adding chilled DMEM with 10% FBS. The solution was then passed through a 40-µm cell strainer. After centrifugation at 300 g for 10 min at 4 °C, the cell pellet was resuspended in chilled DMEM with 10% FBS and passed through another 40-µm cell strainer. The dead cells were removed using Dead Cell Removal Kit (Miltenyi Biotec). Cell number and viability were analyzed using trypan blue staining (Invitrogen). This method generated single cell suspensions with greater than 80% viability.

### Single-Cell RNA Sequencing (scRNA-seq) Library Generation and Sequencing

scRNA-seq library and sequencing were performed at the Yale Center for Genome Analysis (YCGA). Briefly, single cells, reagents and a single Gel Bead containing barcoded oligonucleotides were encapsulated into nanoliter-sized Gel Bead in Emulsion (GEM) using the GemCode^TM^ Technology 10X Genomics. Lysis and barcoded reverse transcription of polyadenylated mRNA from single cells were performed inside each GEM. The scRNA-seq libraries were finished in single bulk reaction. The cDNA libraries were constructed using the 10x Chromium^TM^ Single cell 3’ Library Kit. Qualitative analysis was performed using the Agilent Bioanalyzer High Sensitivity DNA assay as shown in Supplementary Fig. 24. The final libraries from IRI/CL-NX and U-IRI kidneys were sequenced on an Illumina HiSeq 4000 sequencer. Cell Ranger version 5.0.1 was used to process Chromium single cell 3’ RNA-seq output and align the Read to the mouse reference transcriptome (mm10-2020-A), all of which were provided by the YCGA.

### scRNA-seq Data Analysis

Downstream data analysis was performed using the Seurat v4.0 R package. The Seurat integration strategy was performed to identify common cell types and enable comparative analyses between IRI/CL-NX and U-IRI kidneys at each time point^54, 55^. Briefly, all the datasets were first merged for the quality control (QC) analysis. Poor quality cells with <200 unique genes and <500 unique molecular identifier (UMI) counts (likely cell fragment) and >100,000 UMI (potentially cell duplet) were excluded. Cells were excluded if their mitochondrial gene percentages were over 50%. Low-complexity cells like red blood cells with <0.8 log10 genes per UMI counts were also excluded^56^. Only genes expressed in 5 or more cells were used for further analysis. The QC filters resulted in a total of 95,343 cells with a median of 2743 UMI counts per cell at a sequencing depth of 44,719 genes across 95,343 cells. The merged dataset was split, normalized, cell cycle scored, SCTransformed, and integrated using integrated anchors^57^. Confounding sources of variation including mitochondrial gene content were removed for downstream clustering analysis.

Principle component analysis (PCA) was performed on the scaled data. The top 20 principal components were chosen for cell clustering and neighbors finding with k.param = 20, perplexity of 30, and resolution = 0.8. The Uniform Manifold Approximation and Projection (UMAP) was used to visualize the single cells in two-dimensional space. Each cluster was screened for marker genes by differential expression analysis based on the non-parameteric Wilcoxon rank sum test for all clusters with genes expressed in at least 25% of cells either inside or outside of a cluster. Based on the kidney cell and immune cell lineage-specific marker expression, eighteen cell clusters were identified. Both IRI/CL-NX and U-IRI datasets at each time point could be split from the integrated dataset for differential analyses. The average expression of both IRI/CL-NX and U-IRI cells was plotted on using volcano plots. The outliers were used to identify the genes that were differentially expressed between the models in each cluster. All the changes in gene expression were visualized in dot plots or volcano plots. Gene set enrichment analyses were performed using the ClusterProfiler and Gene Ontology (GO) Resource^58^. The potential ligand-receptor interaction analyses were performed using the NicheNet R package^30, 31^ by linking potential ligands expressed by sender cells to their target genes that were differentially expressed by receiver cells and their corresponding receptors. The ligand-receptor pairings for each cell type were visualized by a chord diagram using the R package circlize^59^.

### Quantitative PCR Analysis

Whole kidney RNA was extracted with an RNeasy Mini kit (Qiagen) and reverse transcribed using the iScript cDNA Synthesis Kit (Bio-Rad). Gene expression analysis was determined by quantitative real-time PCR using an iCycler iQ (Bio-Rad) and normalized to hypoxanthine-guanine phosphoribosyltransferase (*Hprt*). The primers included previously published sequences,^21^ and those provided in Supplementary Table 8. The data were expressed using the comparative threshold cycle (ΔCT) method, and the mRNA ratios were given by 2^-ΔCT^.

### Immunohistochemistry (IHC) and Immunofluorescence (IF)

Kidneys were fixed in 10% neutral buffered formalin and embedded in paraffin. Citrate buffer antigen retrieval was used, and the primary antibodies were omitted as negative controls. F4/80-, CD11c-, Ly6G-, CD3ε-, CD4-, and CD8α-positive cells were detected by IHC using primary monoclonal antibodies against F4/80-, CD11c-, Ly6G-, CD3ε-, CD4-, CD8α (#70076, #97585, #87048S, #99940, #25229, and #98941; Cell Signaling Technology, respectively), and megalin (anti-MC220)^60^ as described previously^25^. Lotus tetragonolobus lectin (LTL) was detected by IHC using primary antibodies against biotinylated LTL (#B-1325, Vector Laboratories). All the IHC staining slides were scanned using Aperio LV1 Real-time slide scanner and processed using ImageScope software. Six independent fields in cortex and four independent fields in outer medulla were analyzed per kidney section. The percent area of F4/80-, CD11c-, Ly6G-, CD3ε-, CD4-, and CD8α-positive staining was quantified using IHC Profiler in ImageJ (NIH).

Kidney-specific cadherin (KSP-Cadherin) and UMOD were detected by IF using primary antibodies against UMOD (#sc-20631, Santa Cruz Biotechnology) and KSP-Cadherin (provided by Robert Brent Thomson)^24^. All the IF staining slides were scanned by Pannoramic 250 FLASH III (iHisto Inc, MA), and the images were processed using CaseViewer software. Megalin and KIM-1 were detected by IF using primary antibodies against megalin (anti-MC220)^60^ and TIM-1/KIM-1/HAVCR (#AF1817, Novus Biologicals)^61^. Six to ten random fileds in cortex were imaged per kidney section. The percent area of megalin and KIM-1-positive staining was quantified using ImageJ.

### Confocal microscopy

Three mice were sacrificed 14 days after U-IRI. The kidneys were perfused with 4% paraformaldehyde (PFA) and embedded in optimum cutting temperature (OCT) compound (Tissue Tek). Kidneys were cryosectioned at 5 μm thickness and mounted on Superfrost slides. Sections were washed with PBS, blocked with 10% normal donkey serum, and then stained with primary antibodies against F4/80 (Cl:A3-1, Bio-Rad) and Ly6G or CD3ε (#87048S and #99940, Cell Signaling Technology). The sections were mounted with VECTASHIELD® HardSet™ Antifade Mounting Medium with DAPI (4,6-diamidino-2-phenylindole). The fluorescence images were obtained by confocal microscopy (Zeiss LSM 880).

### Western Blot Analysis

Kidney lysates were fractioned using a RIPA lysis and extraction buffer (Thermo Fisher Scientific) fixed with cOmplete, EDTA-free, protease inhibitor (Roche). Western blot analysis was completed with antibodies selectively against TIM-1/KIM-1/HAVCR (#AF1817, Novus Biologicals) and GAPDH (#51332; Cell Signaling Technology) as described previously^21^.

### Human kidney biopsy

Human kidney biopsy samples were obtained in accordance with the policies of Yale University’s Human Investigations Committee (HIC approval number 11110009286). Written informed consent for research use of patient samples was obtained from each patient by study team. Living donor samples were collected as previously described^36^. The banked AKI samples obtained between 2015 and 2018 were acquired retrospectively after chart review. All data were deidentified and maintained on a secure database. We estimated GFR (eGFR) using the CKD-epidemiology equation^62^. Ten cases were identified using the following criteria: baseline eGFR ≥60 mL/min/1.73m^2^, presence of AKI, exclusion of those with GN, DKD, or AIN, availability of biopsy sample, and availability of 6-month follow-up eGFR (8/10 with pathologic diagnosis of ATI, acute tubular injury). CD3ε- and CD66b-cells were detected by IF using primary monoclonal antibodies against CD3ε (#NBP2-53387, Novus Biologicals) and CD66b (#305102, BioLegend), respectively, and proximal tubules were visualized by IF using polyclonal antibody against LRP2 (#19700-1-AP, Thermo Fisher Scientific) to identify renal cortical regions on the biopsy specimen. All the IF staining sections were tile-scanned by Zeiss LSM 880 with Airyscan Microscope at the Yale Center for Advanced Light Microscopy Facility, and the images were processed using Zeiss Zen lite software. The number of T cells and neutrophils in the entire sample cortex were counted in a blinded manner and normalized to the total number of cells (identified by DAPI+ nuclei) in the same cortical region.

### Statistical Analysis

The data were expressed as means ± standard deviation (SD). Two-group comparison was performed by Student *t*-test. Multigroup comparison was performed by one-way analysis of variance (ANOVA) for group mean comparison followed by Tukey’s multiple comparison test for subgroup comparison. Two-group time-course comparison was performed by two-way ANOVA for model comparison to test whether there is a difference between the models and in the time course, followed by Bonferroni post-tests for subgroup comparison at each time point (Supplementary Table 1, 2). Correlation of gene expression was performed by Pearson correlation coefficient R with two-tailed *P* value (Supplementary Tables 3-5). Correlation of T cells/neutrophils and eGFR was performed by nonparametric Spearman correlation coefficient r with two-tailed *P* value. All the statistical analysis was performed using Prism 8 (GraphPad Software). A value of *P*<0.05 was considered statistically significant.

### Data Availability

All relevant data are available from the corresponding authors on reasonable request. Sequencing data are deposited in GEO under accession number GSE197626.

## ACKNOWLEDGEMENTS

We are grateful to Dr. Joseph Craft from the Section of Rheumatology at the Yale School of Medicine for providing advice in T cell depletion and Robert Brent Thomson for providing the primary antibody against mouse KSP-Cadherin. This work was supported by National Institutes of Health Grant T32 DK007276 and K01 DK120783 (to LX), R01 DK093771 (to LGC), K23 DK117065 (to DGM), P30 DK079310 (George M. O’Brien Kidney Center at Yale), and S10 OD023598 (Yale Center for Advanced Light Microscopy Facility).

## AUTHOR CONTRIBUTIONS

LX designed and performed the primary experiments, analyzed the data, and wrote the manuscript. JG in part performed single cell preparation, data interpretation, and edited manuscript. DGM obtained human samples, performed patient chart review, and edited the manuscript. LGC oversaw the project and edited the manuscript.

## DISCLOSURES

The authors declare no conflict interests.

## Notes

### Competing Interest Statement

The authors have declared no competing interest.

