## Supplementary material for "Immune-mediated Tubule Atrophy Promotes Acute Kidney Injury to Chronic Kidney Disease Transition": IRI repair models-SI_vF.pdf

**Supplemental Table 1.** Two-way ANOVA for gene expression analysis.

| | Time Factor | Model Factor | Interaction |
|---|---|---|---|
| *Havcr1* | **** | *** | *** |
| *Vcam1* | **** | **** | **** |
| *Lrp2* | **** | **** | ** |
| *Slc34a1* | **** | * | ns |
| *Slc13a3* | **** | **** | ** |
| *Cd68* | **** | **** | *** |
| *F4/80* | **** | **** | **** |
| *Itgax* | **** | **** | **** |
| *Ly6g* | **** | ns | ns |
| *Cd3e* | **** | **** | **** |
| *Cd4* | **** | **** | **** |
| *Cd8a* | **** | **** | **** |
| *Ccl4* | **** | **** | **** |
| *Ccl5* | **** | **** | **** |
| *Ccl7* | *** | **** | **** |
| *Ccl8* | **** | * | ** |
| *Ccl12* | **** | **** | **** |
| *Cxcl16* | **** | *** | **** |
| *Ccr1* | **** | ns | * |
| *Ccr2* | **** | **** | **** |
| *Cxcr6* | **** | **** | **** |
| *Il1b* | **** | **** | **** |
| *Tnf* | **** | **** | **** |
| *Fasl* | **** | **** | **** |
| *Ltb* | ** | **** | **** |
| *Cd40lg* | **** | **** | **** |
| *H2-Aa* | **** | **** | **** |
| *H2-Ab1* | **** | **** | **** |
| *Cd74* | **** | **** | **** |
| *Arg1* | **** | ns | ns |
| *Mrc1* | **** | **** | **** |
| *Msr1* | **** | *** | ns |
| *Pdgfb* | **** | **** | **** |
| *Tgfb1* | **** | **** | **** |

*p<0.05; **p<0.01; ***p<0.001; ****p<0.0001; ns, not significant.

**Supplemental Table 2.** Two-way ANOVA for histological assessments in Figure 3b and c.

| | **Cortex/Outer medulla Factor** | **Model Factor** | **Interaction** |
|---|---|---|---|
| F4/80+ area | **** | **** | **** |
| CD11c+ area | ns | **** | **** |
| Ly6G+ area | ns | **** | ns |
| CD3ε+ area | ns | **** | ** |
| CD4+ area | *** | **** | **** |
| CD8α+ area | ns | *** | * |

*p<0.05; **p<0.01; ***p<0.001; ****p<0.0001; ns, not significant.

**Supplemental Table 3.** Pearson correlation coefficient with two-tailed p value analysis for gene expression in the IRI/CL-NX kidneys.

| IRI/CL-NX | | MΦ/DC | | | | | | | | Neutrophil | | | | T Cell | | | | | | | |
|---|---|---|---|---|---|---|---|---|---|---|---|---|---|---|---|---|---|---|---|---|---|
| | *Gene* | *Cd68* | | *F4/80* | | *Itgax* | | *Ccr2* | | *Ly6g* | | *Ccr1* | | *Cd3e* | | *Cd4* | | *Cd8* | | *Cxcr6* | |
| **Identity** | *F4/80* | 0.9510 | *0.0129* | | | | | | | | | | | | | | | | | | |
| | *Itgax* | 0.9659 | *0.0075* | 0.9707 | *0.0060* | | | | | | | | | | | | | | | | |
| | *Ccr2* | 0.9729 | *0.0053* | 0.9963 | *0.0003* | 0.9738 | *0.0051* | | | | | | | | | | | | | | |
| | *Ly6g* | 0.6937 | *0.1939* | 0.4735 | *0.4204* | 0.6455 | *0.2395* | 0.5266 | *0.3619* | | | | | | | | | | | | |
| | *Ccr1* | 0.7526 | *0.1421* | 0.8669 | *0.0571* | 0.7243 | *0.1664* | 0.8545 | *0.0652* | 0.0634 | *0.9194* | | | | | | | | | | |
| | *Cd3e* | 0.6311 | *0.2535* | 0.5623 | *0.3238* | 0.7366 | *0.1557* | 0.5688 | *0.3170* | 0.7969 | *0.1065* | 0.0773 | *0.9017* | | | | | | | | |
| | *Cd4* | 0.8760 | *0.0514* | 0.8326 | *0.0801* | 0.9399 | *0.0175* | 0.8401 | *0.0749* | 0.7992 | *0.1047* | 0.4568 | *0.4393* | 0.9032 | *0.0356* | | | | | | |
| | *Cd8a* | 0.7042 | *0.1843* | 0.6417 | *0.2431* | 0.7930 | *0.1095* | 0.6504 | *0.2347* | 0.7826 | *0.1176* | 0.1810 | *0.7708* | 0.9850 | *0.0022* | 0.9124 | *0.0307* | | | | |
| | *Cxcr6* | 0.5445 | *0.3426* | 0.3666 | *0.5439* | 0.5572 | *0.3292* | 0.4056 | *0.4980* | 0.8781 | *0.0501* | -0.1105 | *0.8596* | 0.8921 | *0.0419* | 0.7325 | *0.1593* | 0.9007 | *0.0370* | | |
| **Mφ phenotype marker** | *Arg1* | 0.6630 | *0.2226* | 0.7867 | *0.1144* | 0.6177 | *0.2669* | 0.7748 | *0.1239* | | | | | | | | | | | | |
| | *Mrc1* | 0.8767 | *0.0510* | 0.9635 | *0.0083* | 0.8727 | *0.0535* | 0.9553 | *0.0113* | | | | | | | | | | | | |
| | *Msr1* | 0.8013 | *0.1031* | 0.8508 | *0.0676* | 0.7279 | *0.1633* | 0.8584 | *0.0626* | | | | | | | | | | | | |
| | *Pdgfb* | 0.8266 | *0.0844* | 0.9503 | *0.0132* | 0.8519 | *0.0669* | 0.9305 | *0.0218* | | | | | | | | | | | | |
| | *Tgfb1* | 0.8830 | *0.0472* | 0.9722 | *0.0055* | 0.8906 | *0.0427* | 0.9623 | *0.0087* | | | | | | | | | | | | |
| **Chemokine** | *Ccl2* | 0.9800 | *0.0034* | 0.9196 | *0.0270* | 0.9341 | *0.0201* | 0.9461 | *0.0149* | 0.6819 | *0.2048* | 0.7139 | *0.1756* | 0.6306 | *0.2540* | 0.8330 | *0.0799* | 0.7263 | *0.1646* | 0.6059 | *0.2787* |
| | *Ccl4* | 0.7675 | *0.1298* | 0.5689 | *0.3169* | 0.6960 | *0.1917* | 0.6236 | *0.2610* | 0.9131 | *0.0303* | 0.2034 | *0.7429* | 0.7563 | *0.1390* | 0.7579 | *0.1377* | 0.8098 | *0.0967* | 0.9075 | *0.0333* |
| | *Ccl5* | 0.9168 | *0.0284* | 0.9240 | *0.0249* | 0.9759 | *0.0045* | 0.9221 | *0.0258* | 0.6678 | *0.2180* | 0.6516 | *0.2336* | 0.7676 | *0.1297* | 0.9644 | *0.0080* | 0.7878 | *0.1136* | 0.5345 | *0.3534* |
| | *Ccl7* | 0.8707 | *0.0547* | 0.9447 | *0.0155* | 0.9161 | *0.0288* | 0.9305 | *0.0218* | 0.4173 | *0.4846* | 0.8411 | *0.0742* | 0.5006 | *0.3903* | 0.8091 | *0.0972* | 0.5343 | *0.3536* | 0.2117 | *0.7324* |
| | *Ccl8* | 0.9291 | *0.0224* | 0.8419 | *0.0736* | 0.9413 | *0.0169* | 0.8644 | *0.0587* | 0.8617 | *0.0604* | 0.4922 | *0.3996* | 0.8479 | *0.0695* | 0.9799 | *0.0034* | 0.8705 | *0.0548* | 0.7455 | *0.1481* |
| | *Ccl12* | 0.9365 | *0.0190* | 0.9854 | *0.0021* | 0.9634 | *0.0084* | 0.9797 | *0.0035* | 0.4886 | *0.4037* | 0.8534 | *0.0659* | 0.5556 | *0.3309* | 0.8473 | *0.0699* | 0.6109 | *0.2737* | 0.3210 | *0.5984* |
| | *Cxcl16* | 0.9457 | *0.0151* | 0.9441 | *0.0157* | 0.9907 | *0.0011* | 0.9462 | *0.0148* | 0.6803 | *0.2063* | 0.6760 | *0.2103* | 0.7640 | *0.1327* | 0.9635 | *0.0083* | 0.7979 | *0.1057* | 0.5591 | *0.3272* |
| **Inflammatory Milieu** | *Tnf* | 0.9508 | *0.0130* | 0.9730 | *0.0053* | 0.9899 | *0.0012* | 0.9712 | *0.0058* | 0.6006 | *0.2841* | 0.7604 | *0.1356* | 0.6836 | *0.2032* | 0.9233 | *0.0252* | 0.7280 | *0.1632* | 0.4628 | *0.4325* |
| | *Ltb* | 0.8570 | *0.0635* | 0.7606 | *0.1354* | 0.8903 | *0.0429* | 0.7828 | *0.1175* | 0.8769 | *0.0509* | 0.3457 | *0.5687* | 0.9328 | *0.0207* | 0.9679 | *0.0069* | 0.9574 | *0.0105* | 0.8677 | *0.0566* |
| | *Fasl* | 0.4687 | *0.4258* | 0.3012 | *0.6223* | 0.5150 | *0.3745* | 0.3327 | *0.5844* | 0.8750 | *0.0520* | -0.2048 | *0.7411* | 0.9245 | *0.0246* | 0.7414 | *0.1516* | 0.8977 | *0.0387* | 0.9769 | *0.0042* |
| | *Cd40lg* | 0.7448 | *0.1487* | 0.5267 | *0.3619* | 0.6822 | *0.2045* | 0.5821 | *0.3031* | 0.9916 | *0.0009* | 0.1344 | *0.8294* | 0.7833 | *0.1171* | 0.8019 | *0.1027* | 0.7914 | *0.1107* | 0.8869 | *0.0449* |
| | *Il1b* | 0.9253 | *0.0242* | 0.9460 | *0.0149* | 0.9734 | *0.0052* | 0.9426 | *0.0163* | 0.6126 | *0.2720* | 0.7238 | *0.1668* | 0.6913 | *0.1961* | 0.9299 | *0.0220* | 0.7178 | *0.1721* | 0.4481 | *0.4492* |
| **MHC** | *H2-Aa* | 0.9113 | *0.0313* | 0.8652 | *0.0582* | 0.9596 | *0.0097* | 0.8754 | *0.0518* | 0.7977 | *0.1059* | 0.5123 | *0.3775* | 0.8732 | *0.0532* | 0.9969 | *0.0002* | 0.8923 | *0.0418* | 0.7134 | *0.1760* |
| | *H2-Ab1* | 0.8901 | *0.0430* | 0.8516 | *0.0671* | 0.9461 | *0.0149* | 0.8586 | *0.0625* | 0.7831 | *0.1173* | 0.5081 | *0.3821* | 0.8546 | *0.0651* | 0.9931 | *0.0007* | 0.8587 | *0.0624* | 0.6637 | *0.2219* |
| | *Cd74* | 0.8378 | *0.0765* | 0.8012 | *0.1032* | 0.9167 | *0.0285* | 0.8049 | *0.1004* | 0.7879 | *0.1135* | 0.4160 | *0.4860* | 0.9115 | *0.0312* | 0.9963 | *0.0003* | 0.9069 | *0.0336* | 0.7180 | *0.1719* |
| **Two-way ANOVA** | | R | *p* | R | *p* | R | *p* | R | *p* | R | *p* | R | *p* | R | *p* | R | *p* | R | *p* | R | *p* |

Statistically significant correlations between immune cell markers and indicated gene expression were highlighted in yellow and green for macrophage/DC, neutrophil, and T cell, respectively. R, Pearson R; p, p value.

**Supplemental Table 4.** Pearson correlation coefficient with two-tailed p value analysis for gene expression in the U-IRI kidneys.

| **U-IRI** | | **MΦ/DC** | | | | | | | | **Neutrophil** | | | | **T Cell** | | | | | | | |
|---|---|---|---|---|---|---|---|---|---|---|---|---|---|---|---|---|---|---|---|---|---|
| | *Gene* | *Cd68* | | *F4/80* | | *Itgax* | | *Ccr2* | | *Ly6g* | | *Ccr1* | | *Cd3e* | | *Cd4* | | *Cd8* | | *Cxcr6* | |
| **Identity** | *F4/80* | 0.9891 | *0.0014* | | | | | | | | | | | | | | | | | | |
| | *Itgax* | 0.9868 | *0.0018* | 0.9817 | *0.0030* | | | | | | | | | | | | | | | | |
| | *Ccr2* | 0.9751 | *0.0047* | 0.9861 | *0.0020* | 0.9604 | *0.0094* | | | | | | | | | | | | | | |
| | *Ly6g* | 0.8361 | *0.0777* | 0.8153 | *0.0926* | 0.8593 | *0.0620* | 0.8702 | *0.0550* | | | | | | | | | | | | |
| | *Ccr1* | 0.7092 | *0.1798* | 0.7023 | *0.1861* | 0.6522 | *0.2329* | 0.8081 | *0.0980* | 0.8313 | *0.0810* | | | | | | | | | | |
| | *Cd3e* | 0.8118 | *0.0952* | 0.7942 | *0.1086* | 0.8777 | *0.0504* | 0.7123 | *0.1771* | 0.6586 | *0.2268* | 0.2362 | *0.7020* | | | | | | | | |
| | *Cd4* | 0.8986 | *0.0381* | 0.8873 | *0.0446* | 0.9529 | *0.0122* | 0.8362 | *0.0776* | 0.7954 | *0.1076* | 0.4347 | *0.4645* | 0.9762 | *0.0044* | | | | | | |
| | *Cd8a* | 0.8325 | *0.0802* | 0.8175 | *0.0910* | 0.8999 | *0.0374* | 0.7472 | *0.1467* | 0.7134 | *0.1760* | 0.2976 | *0.6268* | 0.9966 | *0.0002* | 0.9887 | *0.0014* | | | | |
| | *Cxcr6* | 0.7654 | *0.1315* | 0.7429 | *0.1503* | 0.8307 | *0.0815* | 0.6462 | *0.2388* | 0.5758 | *0.3097* | 0.1412 | *0.8208* | 0.9936 | *0.0006* | 0.9467 | *0.0146* | 0.9812 | *0.0031* | | |
| **Mφ phenotype marker** | *Arg1* | 0.3503 | *0.5632* | 0.3740 | *0.5351* | 0.2136 | *0.7301* | 0.4501 | *0.4469* | | | | | | | | | | | | |
| | *Mrc1* | 0.9684 | *0.0067* | 0.9882 | *0.0015* | 0.9416 | *0.0168* | 0.9848 | *0.0023* | | | | | | | | | | | | |
| | *Msr1* | 0.7774 | *0.1218* | 0.7720 | *0.1261* | 0.6747 | *0.2116* | 0.8227 | *0.0872* | | | | | | | | | | | | |
| | *Pdgfb* | 0.6737 | *0.2124* | 0.7121 | *0.1772* | 0.5719 | *0.3138* | 0.7463 | *0.1474* | | | | | | | | | | | | |
| | *Tgfb1* | 0.9803 | *0.0033* | 0.9763 | *0.0044* | 0.9398 | *0.0176* | 0.9795 | *0.0035* | | | | | | | | | | | | |
| **Chemokine** | *Ccl2* | 0.9203 | *0.0267* | 0.9285 | *0.0227* | 0.8559 | *0.0642* | 0.9459 | *0.0150* | 0.7146 | *0.1750* | 0.8162 | *0.0919* | 0.5339 | *0.3540* | 0.6630 | *0.2225* | 0.5592 | *0.3271* | 0.4781 | *0.4153* |
| | *Ccl4* | 0.9484 | *0.0140* | 0.9085 | *0.0328* | 0.9639 | *0.0082* | 0.8698 | *0.0553* | 0.8178 | *0.0908* | 0.5440 | *0.3432* | 0.9228 | *0.0254* | 0.9611 | *0.0092* | 0.9330 | *0.0206* | 0.8953 | *0.0400* |
| | *Ccl5* | 0.9823 | *0.0028* | 0.9926 | *0.0008* | 0.9748 | *0.0048* | 0.9981 | *<0.0001* | 0.8691 | *0.0557* | 0.7746 | *0.1241* | 0.7532 | *0.1416* | 0.8671 | *0.0570* | 0.7856 | *0.1152* | 0.6907 | *0.1966* |
| | *Ccl7* | 0.6896 | *0.1976* | 0.6893 | *0.1980* | 0.6647 | *0.2210* | 0.7986 | *0.1051* | 0.8945 | *0.0404* | 0.9776 | *0.0040* | 0.2907 | *0.6351* | 0.4884 | *0.4038* | 0.3598 | *0.5519* | 0.1878 | *0.7623* |
| | *Ccl8* | 0.9682 | *0.0068* | 0.9672 | *0.0071* | 0.9871 | *0.0018* | 0.9696 | *0.0063* | 0.9241 | *0.0248* | 0.7296 | *0.1618* | 0.8293 | *0.0825* | 0.9296 | *0.0222* | 0.8637 | *0.0591* | 0.7672 | *0.1300* |
| | *Ccl12* | 0.8796 | *0.0492* | 0.8769 | *0.0509* | 0.8632 | *0.0594* | 0.9417 | *0.0168* | 0.9522 | *0.0125* | 0.9367 | *0.0189* | 0.5518 | *0.3349* | 0.7174 | *0.1725* | 0.6068 | *0.2779* | 0.4637 | *0.4315* |
| | *Cxcl16* | 0.9571 | *0.0106* | 0.9427 | *0.0163* | 0.9500 | *0.0133* | 0.9728 | *0.0054* | 0.9517 | *0.0126* | 0.8474 | *0.0699* | 0.7158 | *0.1739* | 0.8449 | *0.0716* | 0.7572 | *0.1382* | 0.6448 | *0.2401* |
| **Inflammatory Milieu** | *Tnf* | 0.9732 | *0.0052* | 0.9753 | *0.0046* | 0.9506 | *0.0131* | 0.9968 | *0.0002* | 0.8830 | *0.0472* | 0.8408 | *0.0744* | 0.6857 | *0.2013* | 0.8157 | *0.0923* | 0.7219 | *0.1685* | 0.6183 | *0.2663* |
| | *Ltb* | 0.9491 | *0.0137* | 0.9427 | *0.0163* | 0.9856 | *0.0021* | 0.9264 | *0.0237* | 0.8991 | *0.0379* | 0.6265 | *0.2581* | 0.9006 | *0.0371* | 0.9733 | *0.0052* | 0.9285 | *0.0227* | 0.8490 | *0.0688* |
| | *Fasl* | 0.7627 | *0.1337* | 0.7329 | *0.1589* | 0.8265 | *0.0845* | 0.6383 | *0.2464* | 0.5849 | *0.3002* | 0.1470 | *0.8135* | 0.9917 | *0.0009* | 0.9443 | *0.0156* | 0.9793 | *0.0036* | 0.9987 | *<0.0001* |
| | *Cd40lg* | 0.8119 | *0.0951* | 0.7653 | *0.1315* | 0.8590 | *0.0622* | 0.6859 | *0.2011* | 0.6545 | *0.2308* | 0.2528 | *0.6816* | 0.9782 | *0.0038* | 0.9491 | *0.0137* | 0.9700 | *0.0062* | 0.9794 | *0.0035* |
| | *Il1b* | 0.7240 | *0.1667* | 0.7024 | *0.1859* | 0.7178 | *0.1721* | 0.7989 | *0.1049* | 0.9562 | *0.0109* | 0.9336 | *0.0203* | 0.4148 | *0.4874* | 0.5906 | *0.2944* | 0.4805 | *0.4126* | 0.3188 | *0.6011* |
| **MHC** | *H2-Aa* | 0.9569 | *0.0107* | 0.9533 | *0.0120* | 0.9902 | *0.0012* | 0.9170 | *0.0283* | 0.8333 | *0.0796* | 0.5554 | *0.3311* | 0.9321 | *0.0210* | 0.9850 | *0.0022* | 0.9502 | *0.0132* | 0.8922 | *0.0418* |
| | *H2-Ab1* | 0.9484 | *0.0139* | 0.9455 | *0.0152* | 0.9822 | *0.0028* | 0.9405 | *0.0173* | 0.9223 | *0.0257* | 0.6740 | *0.2122* | 0.8681 | *0.0564* | 0.9550 | *0.0114* | 0.9011 | *0.0368* | 0.8094 | *0.0969* |
| | *Cd74* | 0.9443 | *0.0156* | 0.9441 | *0.0157* | 0.9833 | *0.0026* | 0.9298 | *0.0221* | 0.8994 | *0.0377* | 0.6303 | *0.2544* | 0.8928 | *0.0414* | 0.9691 | *0.0065* | 0.9224 | *0.0256* | 0.8387 | *0.0759* |
| **Two-way ANOVA** | | R | *p* | R | *p* | R | *p* | R | *p* | R | *p* | R | *p* | R | *p* | R | *p* | R | *p* | R | *p* |

Statistically significant correlations between immune cell markers and indicated gene expression were highlighted in yellow and green for macrophage/DC, neutrophil, and T cell, respectively. R, Pearson R; p, p value.

**Supplemental Table 5.** Pearson correlation coefficient with two-tailed p value analysis for gene expression.

| | **Model** | **U-IRI** | | | | | | **IRI/CL-NX** | | | | | |
|---|---|---|---|---|---|---|---|---|---|---|---|---|---|
| | ***MHC*** | *H2-Aa* | | *H2-Ab1* | | *Cd74* | | *H2-Aa* | | *H2-Ab1* | | *Cd74* | |
| **Inflammatory Milieu** | *Tnf* | 0.9002 | *0.0373* | 0.9307 | *0.0217* | 0.9159 | *0.0289* | 0.9435 | *0.0160* | 0.9446 | *0.0155* | 0.9067 | *0.0338* |
| | *Ltb* | 0.9907 | *0.0011* | 0.9974 | *0.0002* | 0.9990 | *<0.0001* | 0.9659 | *0.0075* | 0.9394 | *0.0177* | 0.9524 | *0.0124* |
| | *Fasl* | 0.8872 | *0.0447* | 0.8071 | *0.0987* | 0.8348 | *0.0786* | 0.7093 | *0.1797* | 0.6792 | *0.2073* | 0.7452 | *0.1484* |
| | *Cd40lg* | 0.9036 | *0.0354* | 0.8376 | *0.0767* | 0.8570 | *0.0635* | 0.8078 | *0.0982* | 0.7816 | *0.1185* | 0.7800 | *0.1197* |
| | *Il1b* | 0.6597 | *0.2258* | 0.7873 | *0.1139* | 0.7493 | *0.1449* | 0.9455 | *0.0151* | 0.9576 | *0.0104* | 0.9216 | *0.0261* |
| **Two-way ANOVA** | | R | *p* | R | *p* | R | *p* | R | *p* | R | *p* | R | *p* |

Statistically significant correlations between major histocompatibility complex (MHC) and inflammatory cytokine expression were highlighted in yellow and green for the U-IRI kidneys and IRI/CL-NX kidneys, respectively. R, Pearson R; p, p value.

**Supplemental Table 6.** Characteristics of participants included in the study.

| **Characteristic** | n (%) or Median (IQR) |
|---|---|
| **Total N** | 10 |
| **Demographics** | |
| Age, years | 58 (27, 64) |
| Female | 5 (50%) |
| Black race | 2 (20%) |
| **Laboratory Features at biopsy** | #VALUE! |
| Creatinine, mg/dl | 4.0 (2.3, 6.6) |
| Blood Urea Nitrogen, mg/dl | 37 (31, 59) |
| **eGFR trend, ml/min** | |
| eGFR before AKI | 75.7 (70.1, 92.7) |
| eGFR at biopsy | 15.6 (9.5, 31.0) |
| eGFR 6-m after | 58.0 (41.3, 83.7) |
| eGFR loss from pre-biopsy to biopsy | -61.3 (-70.7, -36.0) |
| eGFR gain from biopsy to post-biopsy | 33.8 (20.9, 49.1) |
| **Urine tests** | |
| Albuminuria, mg/g | 54 (19, 117) |
| Leukocyte esterase | |
| 1+ | 5 (33%) |
| 2+ | 1 (56%) |
| Granular casts | |
| 1-5/LPF | 3 (38%) |
| >5/LPF | 5 (63%) |

**Supplemental Table 7.** Diagnosis, etiology, and clinical scenario of participants included in the study.

| Case | Diagnosis | Clinical scenario and presumed etiology |
|---|---|---|
| # 0284 | ATN | Severe AS, Contrast use (Cath/CT) for TAVR workup |
| # 0252 | ATN | Vancomycin, Hypotension, Sepsis |
| # 0175 | ATN | Cirrhosis; Sepsis; NSAIDs |
| # 0224 | ATN | Hypotension; Diarrhea |
| # 0222 | Myeloma | Myeloma cast nephropathy |
| # 0101 | ATN | Hypotension; NSAID |
| # 0082 | ATN | Cisplatin |
| # 0022 | ATN | Hypotension; Cirrhosis; Large Volume Paracentesis; Sepsis |
| # 0017 | ATN | Hypotension; Cirrhosis; Sepsis; Diuresis |
| # 0034 | Arterionephrosclerosis | Chronic NSAID use |

Case number represents de-identified study number.

**Supplemental Table 8.** Primer sequences used for quantitative PCR.

| Gene | Forward | Reverse |
|---|---|---|
| *Ccl4* | CCAAGCCAGCTGTGGTATTC | CTGCTCAGTTCAACTCCAAGTC |
| *Ccl5* | CCAATCTTGCAGTCGTGTTTG | AGCAATGACAGGGAAGCTATAC |
| *Ccl7* | CCTGGGAAGCTGTTATCTTCAA | GGTTTCTGTTCAGGCACATTTC |
| *Ccl12* | GCTGTGATCTTCAGGACCATAC | TGGAACTCTCAGCCTAGACAT |
| *Ccr1* | ATACTCTGGAAACACAGACTCAC | CCCACCACTCCAATGATGAA |
| *Cd40lg* | CTGAACTGTGAGGAGATGAGAAG | TGTAGAACGGATGCTGCATTA |
| *Cd74* | GGTCAAGTCACCCTGTGAAG | GGTGTGACATCAGGGAACATAA |
| *Cxcl16* | TGTCCATTCTTTATCAGGTTCCA | AACTCTTCCCATGACCAGTTC |
| *Cxcr6* | CTGGGCTTCTCTTCTGATGC | CGTTTGTTCTCCTGGCTGTTA |
| *Fasl* | TGGCCCATTTAACAGGGAAC | CAACCTCTTCTCCTCCATTAGC |
| *H2-Aa* | GAAACACTGGGAACCTGAGAT | TTCCAGGGTGTGACTCATAAAG |
| *H2-Ab1* | CATCTACAACCGGGAGGAGTA | ACGACATTGGGCTGTTCAAG |
| *Il1b* | TGTGAAATGCCACCTTTTGA | TGTCCTCATCCTGGAAGGTC |
| *Ly6g* | TTGTGGACTCTCACAGAAGC | GTCTTCACGTTGACAGCATTAC |
| *Ltb* | CTCATAGGCGCTTGGATGA | GACGTGGCAGTAGAGGTAATAG |

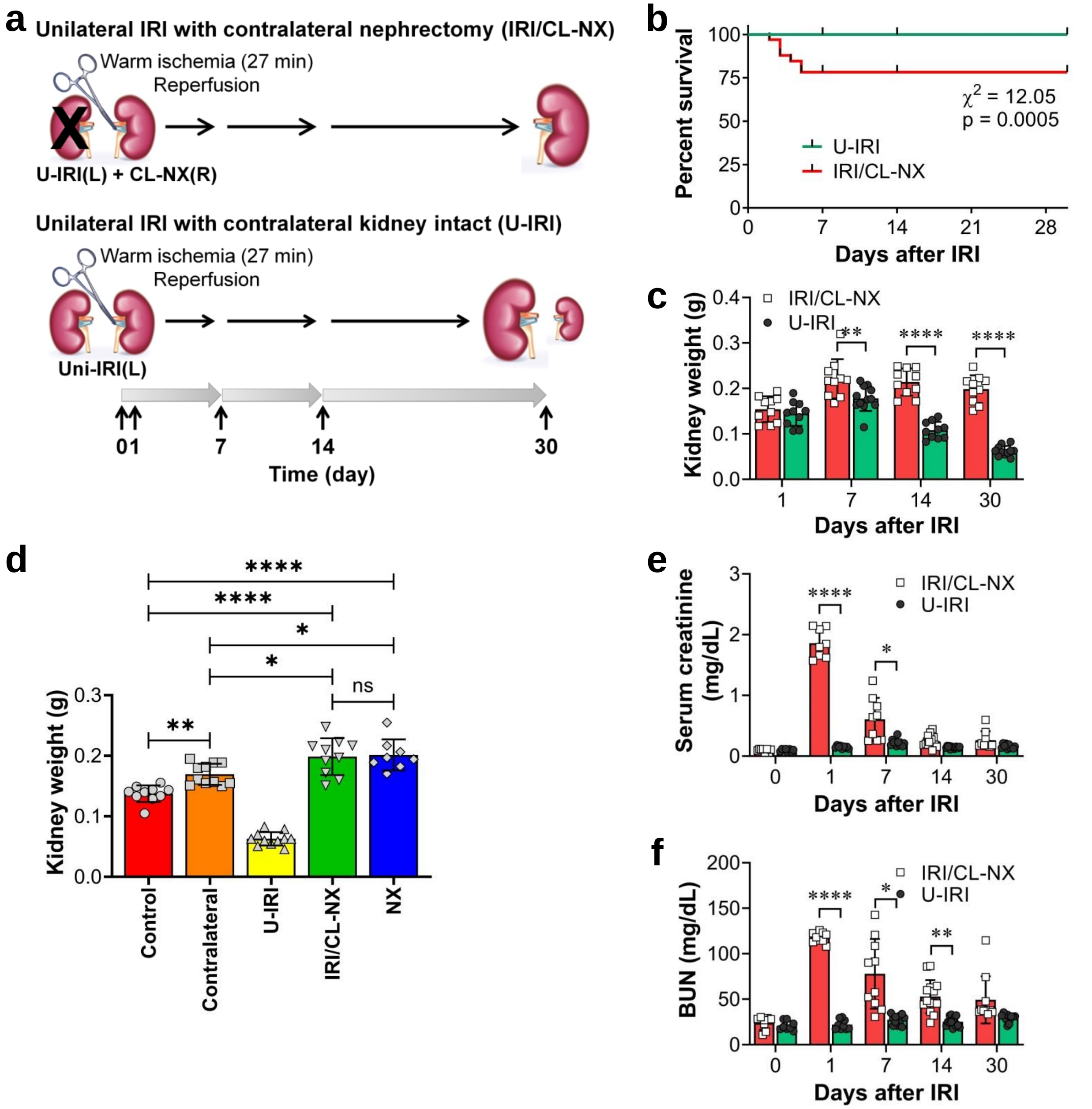


**Supplemental Figure 1.** Models of ischemia/reperfusion injury. **a.** Scheme of U-IRI and IRI/CL-NX mouse models. Kidneys from mice subjected to 27 minutes of unilateral IRI (U-IRI) were compared to kidneys from mice subjected to 27 minutes of IRI with contralateral nephrectomy (IRI/CL-NX). The mice were sacrificed on day 1, 7, 14 and 30 after injury. **b.** The survival rate was determined in both mouse models. Starting n for U-IRI = 49 mice, and for IRI/CL-NX = 99 mice. Approximately 79% of mice survived for 7 days after IRI/CL-NX; whereas 100% of mice survived after U-IRI. **c.** Kidney weights were determined on day 1, 7, 14, and 30 after injury. n=10 kidneys/time point. $p<0.0001$ between models and in time series; **$p<0.01$ and ****$p<0.0001$ in the indicated subgroup analyses. **d.** Kidney weights were determined on day 30 for U-IRI kidneys and their contralateral kidneys, IRI/CL-NX kidneys, kidneys with CL-NX alone (NX), and age-matched healthy kidneys (Control). n = 10 kidneys (8 for NX). $P<0.0001$ among the groups (ANOVA). *p<0.05, **p<0.01, and ****p<0.0001 for the subgroup comparison. ns, not statistically significant. **e** and **f**. Serum creatinine and BUN levels of mice at indicated time points after injury. n=10 mice/time point. p<0.0001 between models and in time series; *p<0.05, **p<0.01, and ***p<0.001 for the subgroup analyses at each time point. Renal function analyses revealed the expected rise in serum creatinine and BUN on day 1 after injury in the mice subjected to IRI/CL-NX followed by significant improvement on day 7 and return to near baseline by days 14-30 as kidney repair occurred. In contrast, serum creatinine and BUN levels showed minimal changes in mice subjected to the U-IRI alone, reflecting the preserved clearance by the uninjured contralateral kidney.

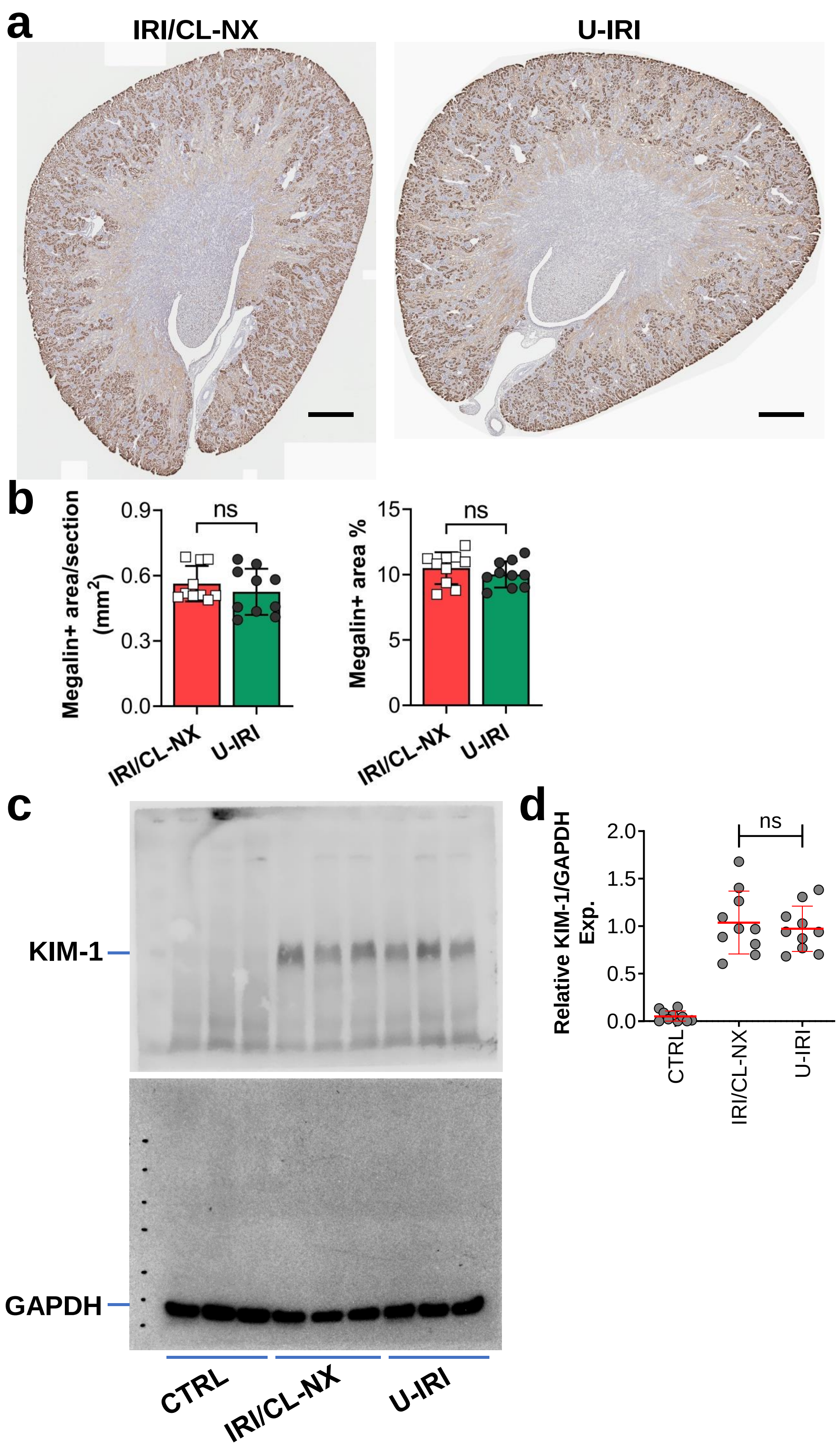

a
IRI/CL-NX
U-IRI
b
Megalin+ area/section (mm2)
0.9
0.6
0.3
0.0
ns
IRI/CL-NX
U-IRI
Megalin+ area %
15
10
5
0
ns
IRI/CL-NX
U-IRI
c
KIM-1
GAPDH
CTRL
IRI/CL-NX
U-IRI
d
Relative KIM-1/GAPDH Exp.
2.0
1.5
1.0
0.5
0.0
ns
CTRL
IRI/CL-NX
U-IRI

**Supplemental Figure 2.** IRI leads to equivalent acute kidney injury between U-IRI and IRI/CL-NX models. **a.** Midline kidney sections underwent IHC staining for megalin (brown staining) on day 1 after IRI. Scale bars, 0.5 mm. **b.** Megalin-positive area as in (a) was quantified for the entire kidney section (left panel) and as a percentage of the section area (right panel). n=10 kidneys/group. ns, not statistically significant. Western blot analysis for KIM-1 protein expression was performed on whole kidney lysates (each lane is from a separate kidney) and (**d**) quantified and normalized to GAPDH. ns, not statistically significant.

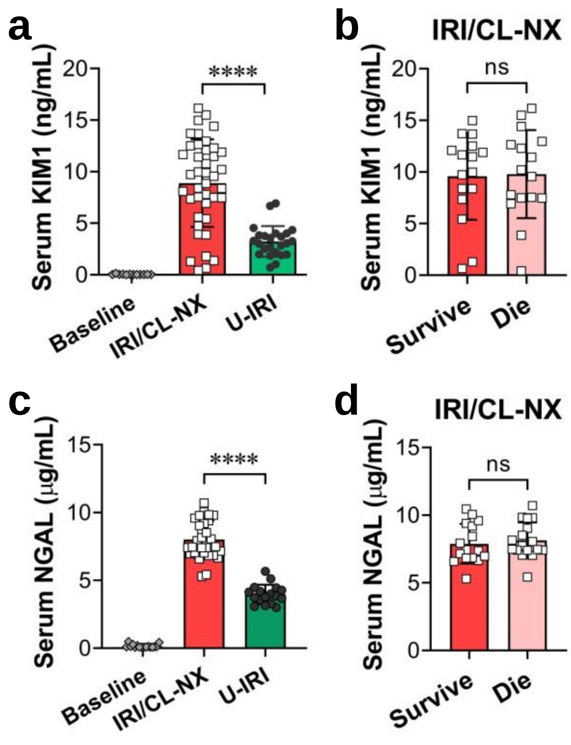


**Supplemental Figure 3.** IRI leads to acute kidney injury in both models. WT mice were subjected to IRI with contralateral nephrectomy (IRI/CL-NX) or unilateral IRI (U-IRI). **a.** Serum KIM1 levels were determined by ELISA at baseline and on day 1 after injury. **b.** Serum Kim1 levels in mice subjected to IRI/CL-NX that either survived (and thus were included in the subsequent analyses) or died prior to day 7. **c.** Serum NGAL levels were determined by ELISA at baseline and on day 1 after injury. **d.** Serum NGAL levels in mice subjected to IRI/CL-NX that either survived (and thus were included in the subsequent analyses) or died prior to day 7.**** $p<0.0001$; ns, not statistically significant.

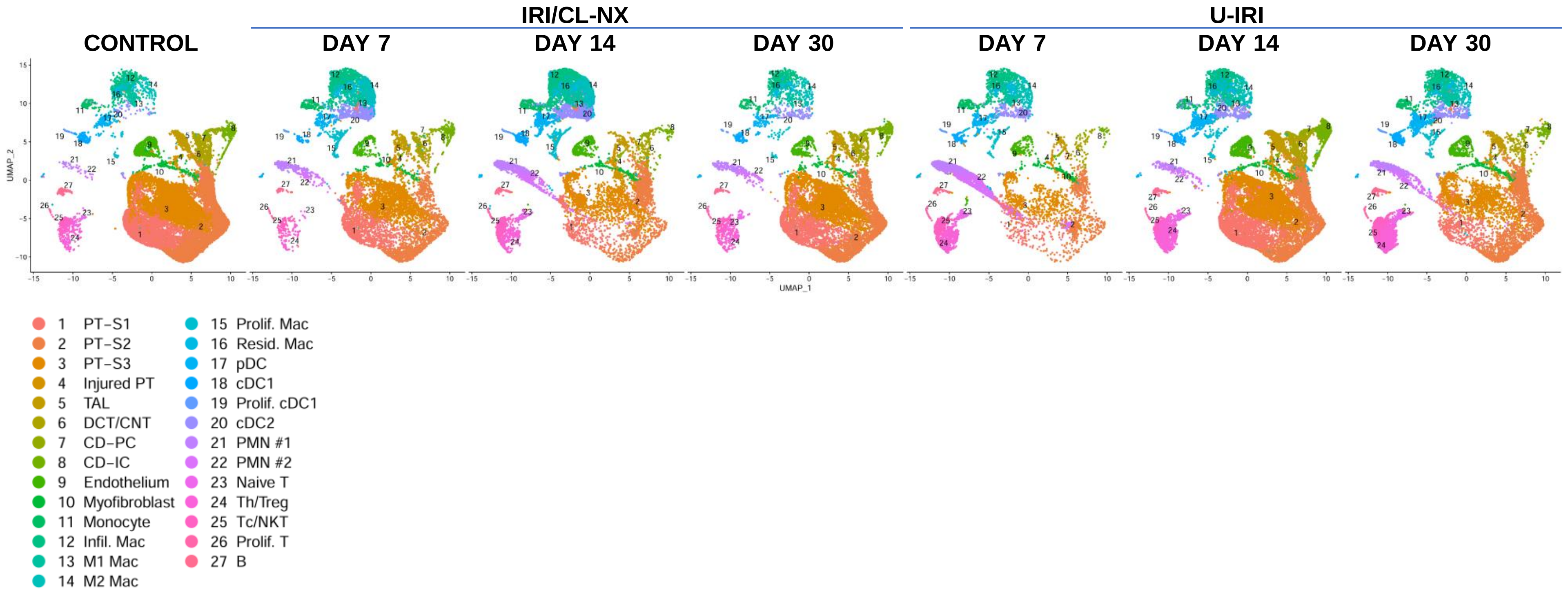


**Supplemental Figure 4.** UMAP projection of cells from U-IRI, IRI/CL-NX, and control healthy kidneys . The cell clusters were identified using the composite data from all cells (Figure 3a), and compared in each model at each time point by kidney cell and immune cell lineage-specific marker expression as shown in (Figure 3b). PT, proximal tubule; TAL, thick ascending limb; DCT, distal convoluted tubule; CNT, connecting tubule; CD-PC, collecting duct-principal cell; CD-IC, collecting duct-intercalated cell; Infil. Mac, infiltrating macrophage; pDC, plasmacytoid dendritic cell; cDC, conventional dendritic cell; PMN, polymorphonuclear neutrophils; Th/Treg, T helper/regulatory T cells; Tc/NKT, cytotoxic T/natural killer T cells; B, B cells.

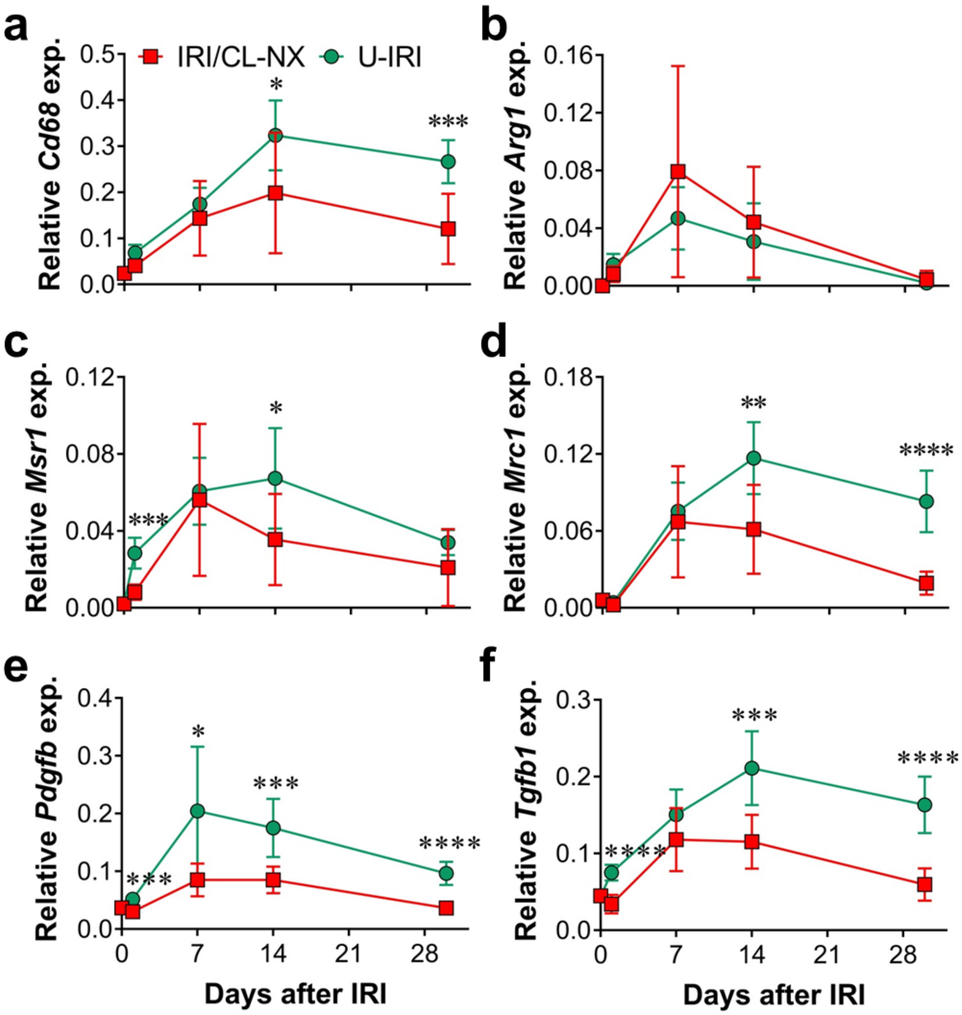


**Supplemental Figure 5.** Macrophages transition from reparative to profibrotic gene expression in the U-IRI kidney 7 days after injury. **a-f.** Quantitative RT-PCR analysis for indicated genes was performed on whole kidney RNA harvested on day 0, 1, 7, 14, and 30 after injury. n=10 kidneys/time point. Two-way ANOVA was summarized in Supplemental Table 1. *p<0.05, **p<0.01, ***p<0.001, and ****p<0.0001 at each time point.

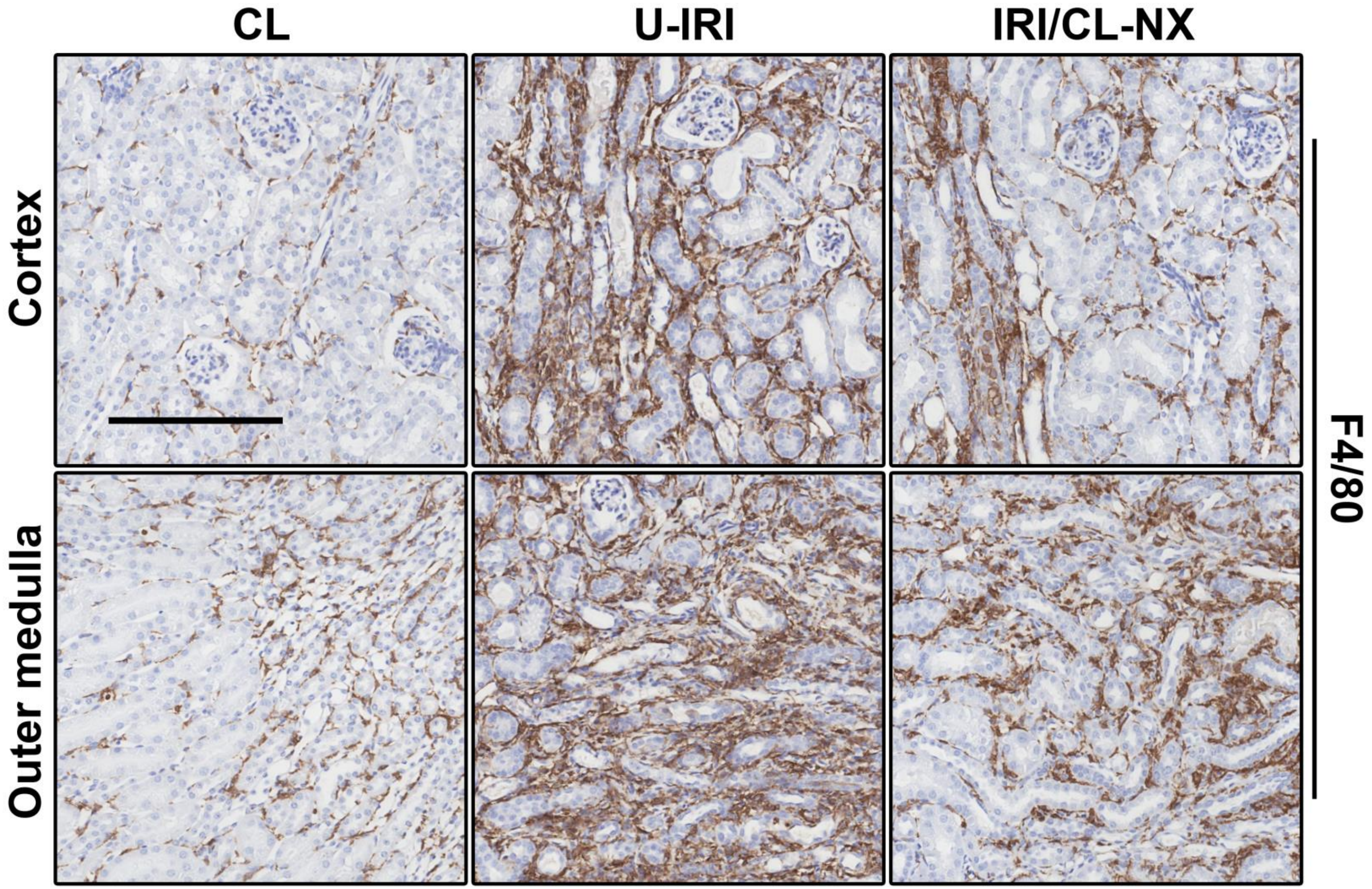


**Supplemental Figure 6.** U-IRI promotes late macrophage accumulation. Contralateral (CL), unilateral IRI (U-IRI) and IRI with contralateral nephrectomy (IRI/CL-NX) kidneys were harvested at 14 days after injury. The kidney sections were immunostained with F4/80. Representative images of kidney section were shown at lower magnification. Scale bars, 200 µm. n=10 kidneys/group.

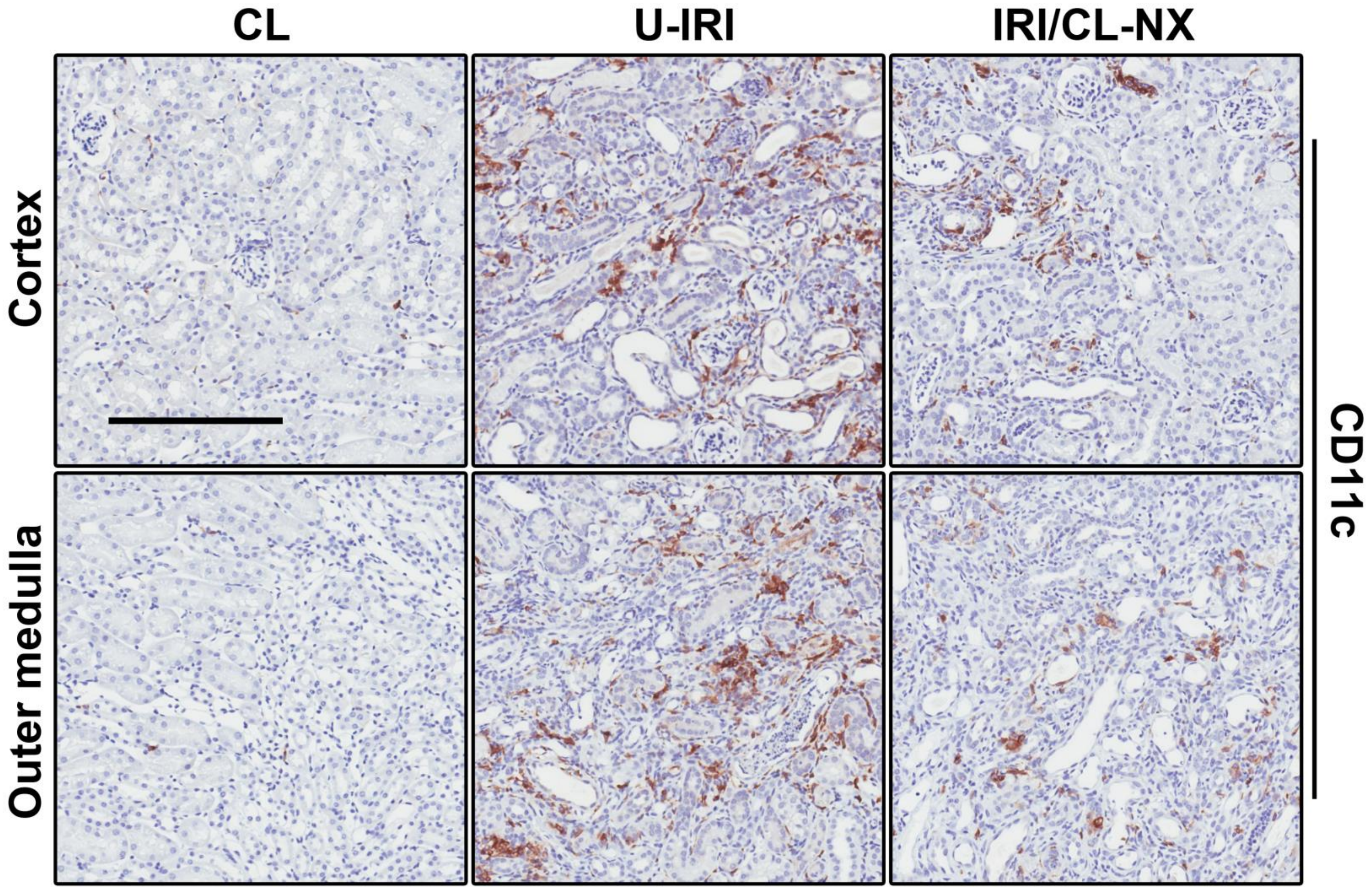


**Supplemental Figure 7.** U-IRI promotes late dendritic cell accumulation. Contralateral (CL), unilateral IRI (U-IRI) and IRI with contralateral nephrectomy (IRI/CL-NX) kidneys were harvested at 14 days after injury. The kidney sections were immunostained with CD11c. Representative images of kidney section were shown at lower magnification. Scale bars, 200 µm. n=10 kidneys/group.

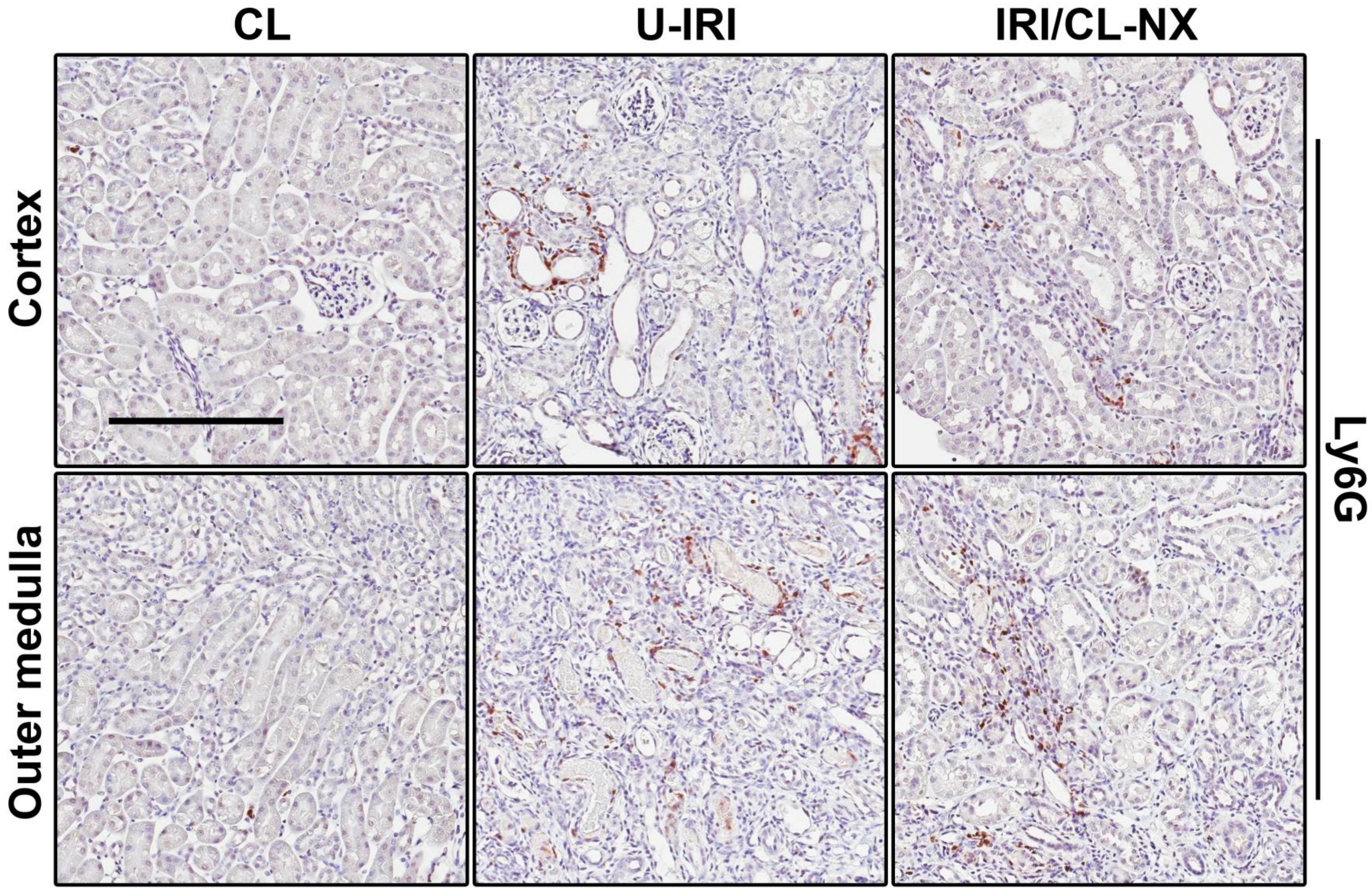


**Supplemental Figure 8.** U-IRI promotes late neutrophil accumulation. Contralateral (CL), unilateral IRI (U-IRI) and IRI with contralateral nephrectomy (IRI/CL-NX) kidneys were harvested at 14 days after injury. The kidney sections were immunostained with Ly6G. Representative images of kidney section were shown at lower magnification. Scale bars, 200 µm. n=10 kidneys/group.

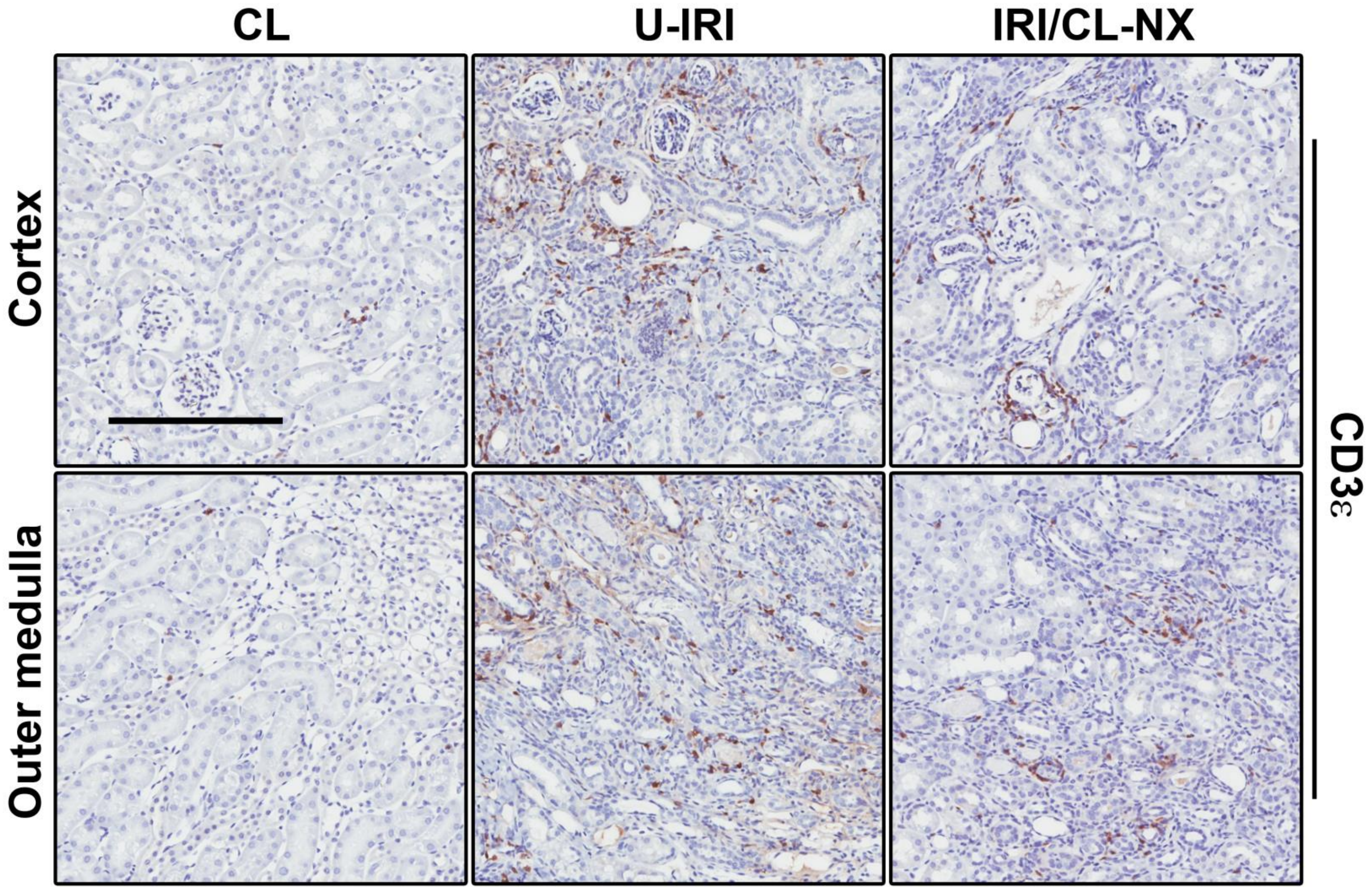


**Supplemental Figure 9.** U-IRI promotes late T cell accumulation. Contralateral (CL), unilateral IRI (U-IRI) and IRI with contralateral nephrectomy (IRI/CL-NX) kidneys were harvested at 14 days after injury. The kidney sections were immunostained with CD3ε. Representative images of kidney section were shown at lower magnification. Scale bars, 200 µm. n=10 kidneys/group.

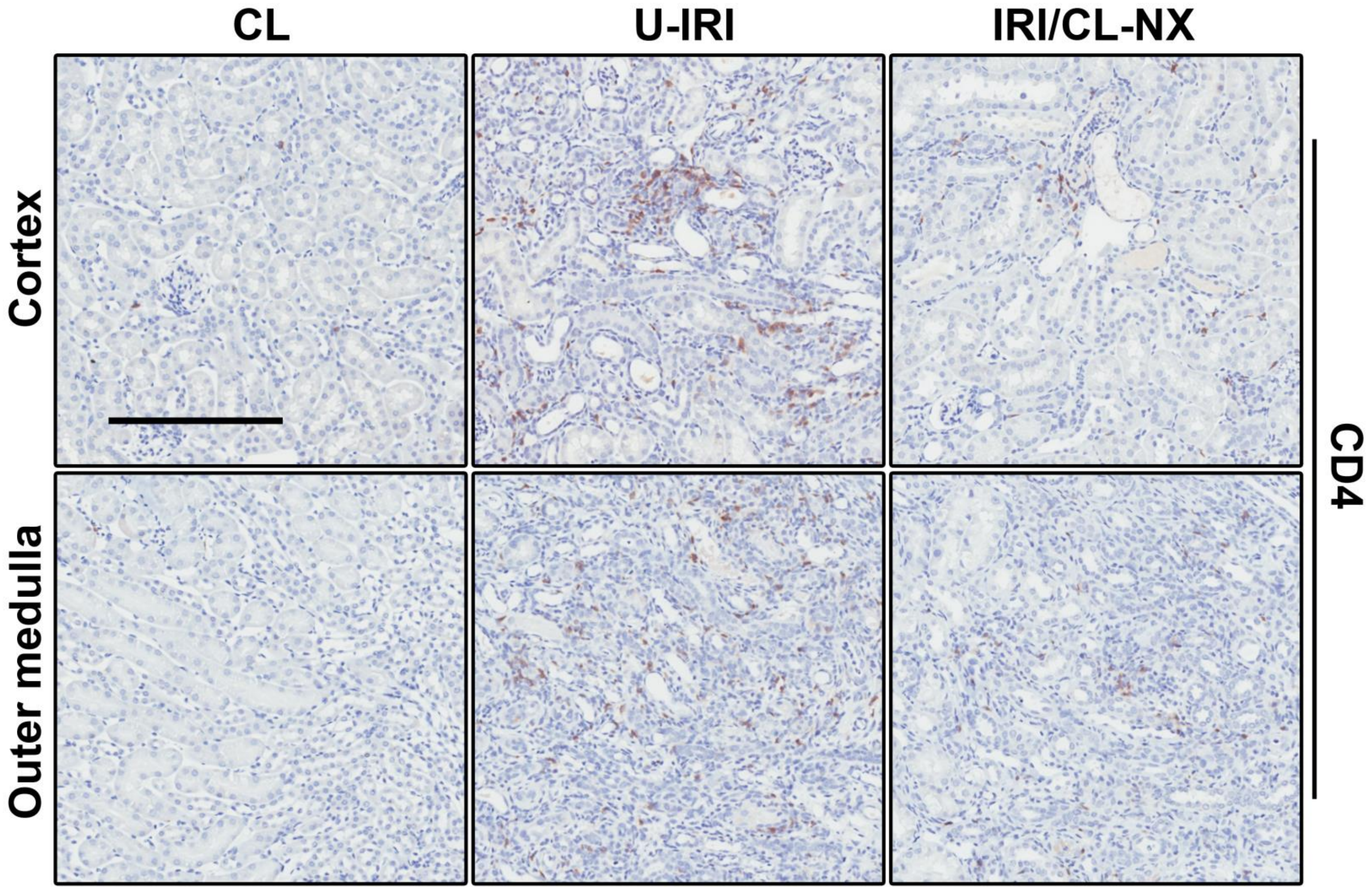


**Supplemental Figure 10.** U-IRI promotes late T helper cell accumulation. Contralateral (CL), unilateral IRI (U-IRI) and IRI with contralateral nephrectomy (IRI/CL-NX) kidneys were harvested at 14 days after injury. The kidney sections were immunostained with CD4. Representative images of kidney section were shown at lower magnification. Scale bars, 200 µm. n=10 kidneys/group.

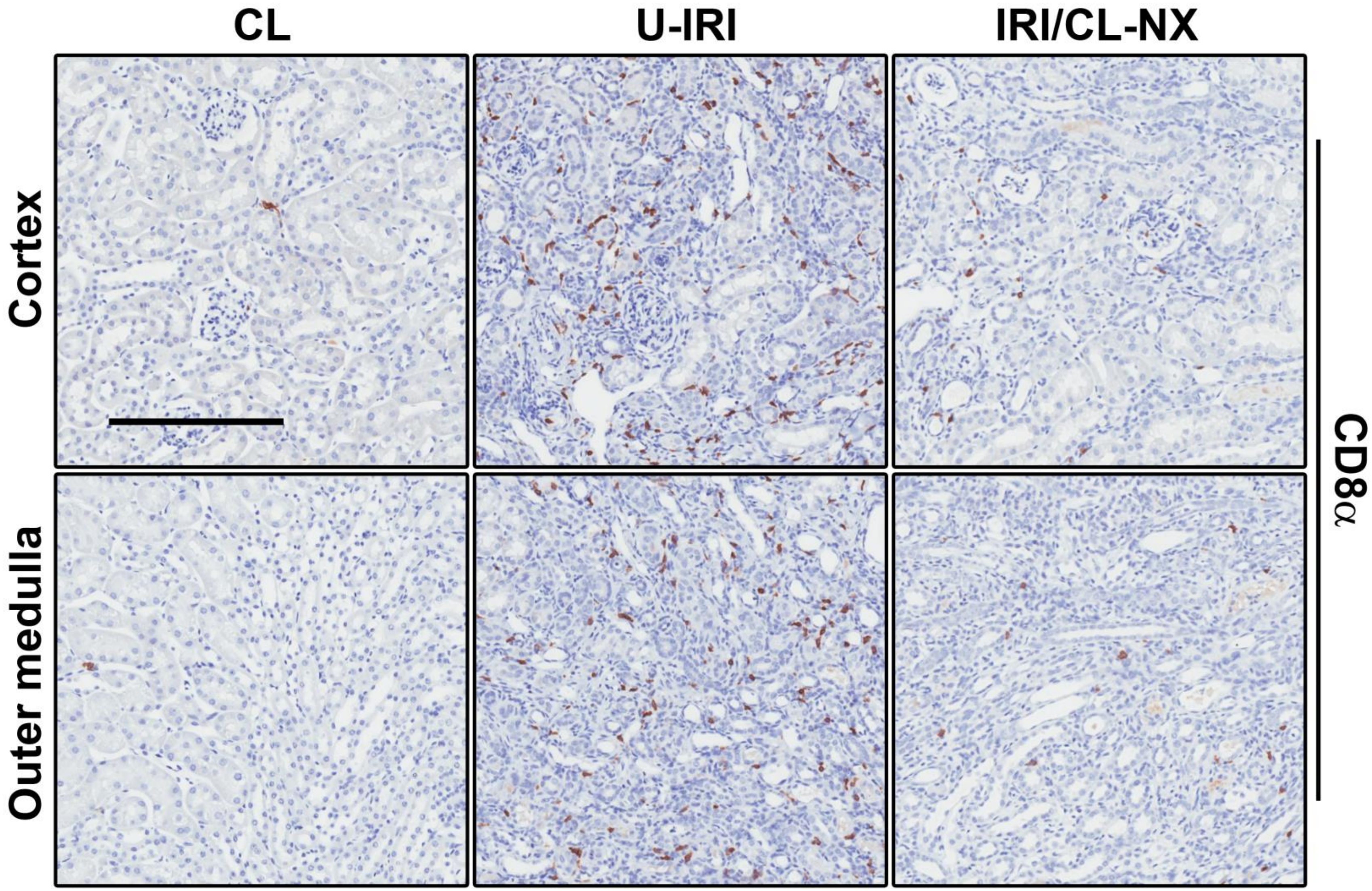


**Supplemental Figure 11.** U-IRI promotes late cytotoxic T cell accumulation. Contralateral (CL), unilateral IRI (U-IRI) and IRI with contralateral nephrectomy (IRI/CL-NX) kidneys were harvested at 14 days after injury. The kidney sections were immunostained with CD8α. Representative images of kidney section were shown at lower magnification. Scale bars, 200 µm. n=10 kidneys/group.

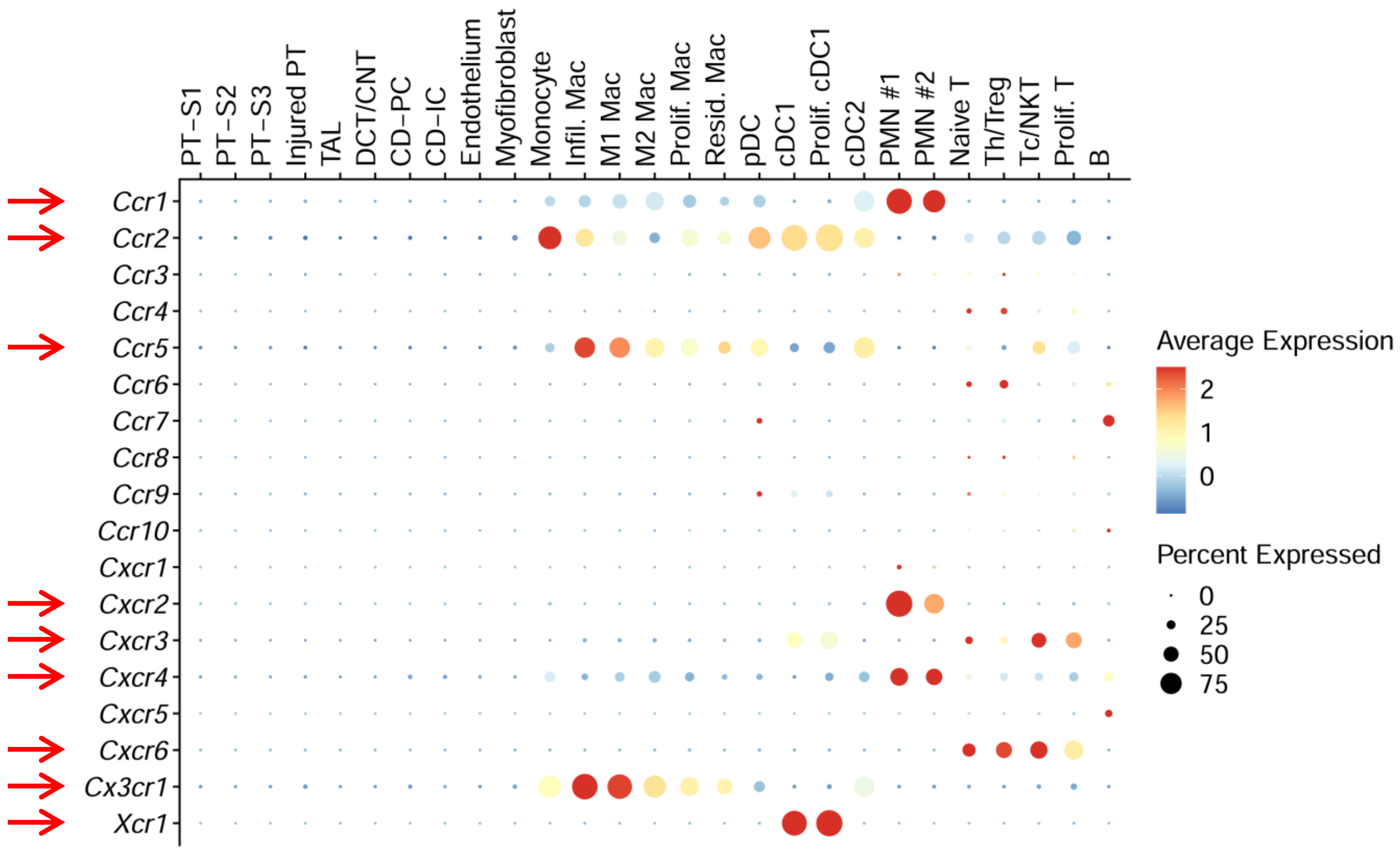


**Supplemental Figure 12.** The distribution and relative expression of chemokine receptors are shown in a dot plot using the integrated dataset.

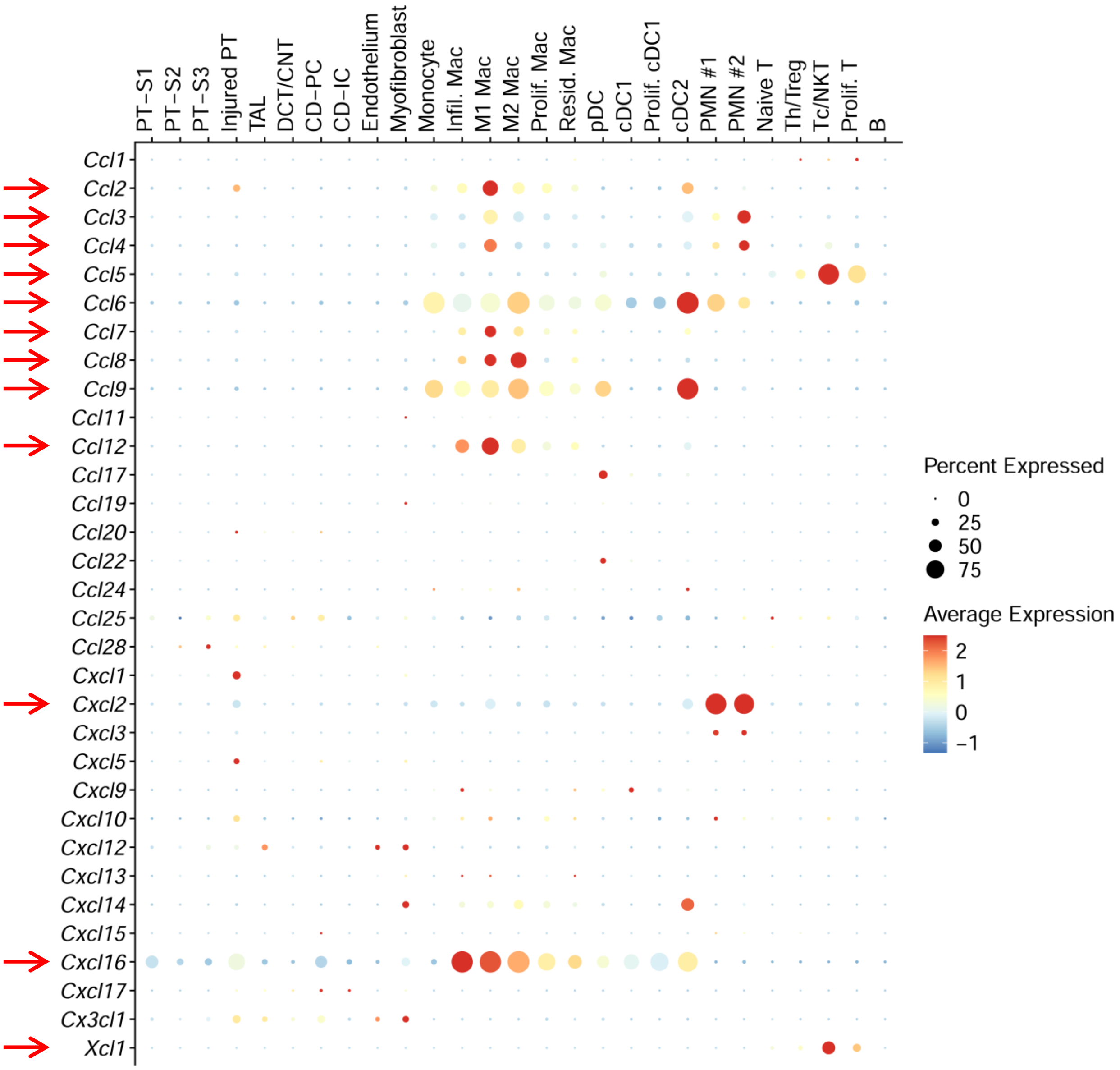


**Supplemental Figure 13.** The distribution and relative expression of corresponding chemokine ligands are shown in a dot plot using the integrated dataset.

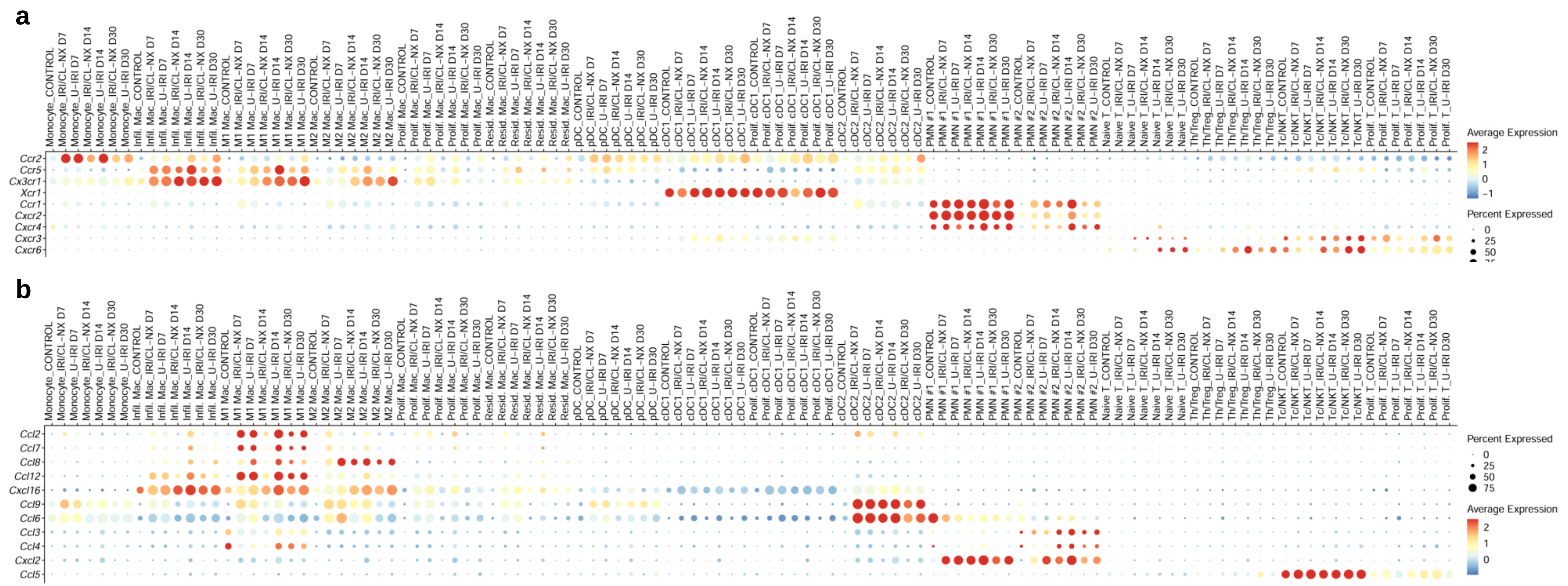


**Supplemental Figure 14.** The distribution and relative expression of key chemokine receptors (**a**) and corresponding chemokine ligands (**b**) in macrophage, dendritic cells, PMNs, and T cells are shown in the dot plots using the split datasets.

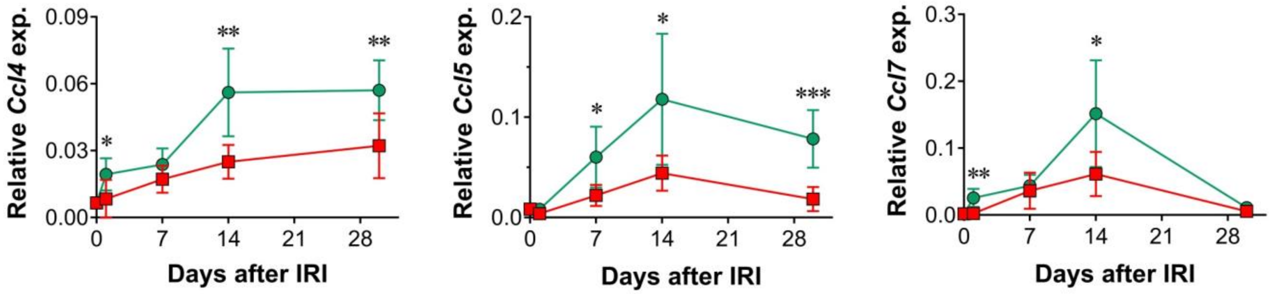


**Supplemental Figure 15.** Chemokine expression kinetics. **a-c.** Quantitative RT-PCR analysis for indicated genes was performed on whole kidney RNA harvested on day 0, 1, 7, 14, and 30 after injury. n=10 kidneys/time point. Two-way ANOVA was summarized in Supplemental Table 1. *p<0.05, **p<0.01, ***p<0.001, and ****p<0.0001 at each time point.

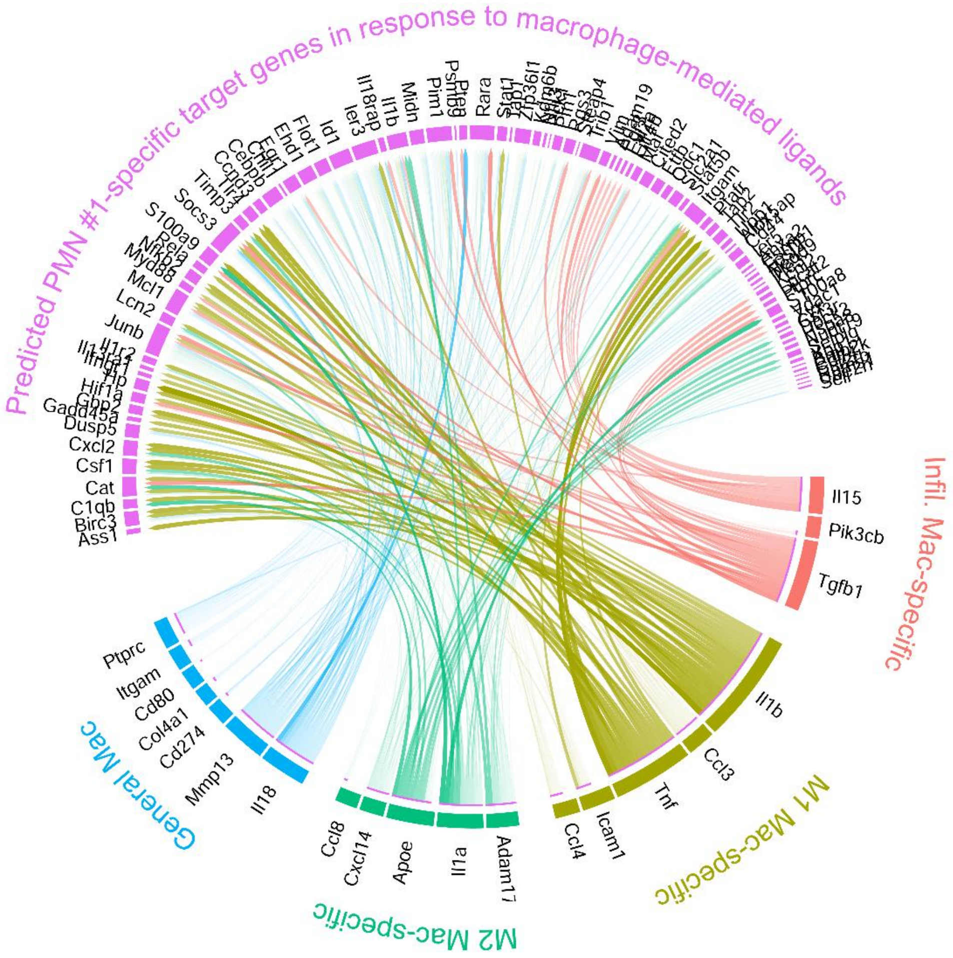


**Supplementary Figure 16.** Ligand-receptor-target interactions between infiltrating, M1, and M2 macrophages with PMN #1. Based on DEG between U-IRI and IRI/CL-NX kidneys on day 14 after injury, the corresponding ligands that were significantly expressed by the infiltrating, M1, and M2 macrophages were identified and linked to their potential target genes that were upregulated by PMN #1 and visualized using a chord diagram.

**a**

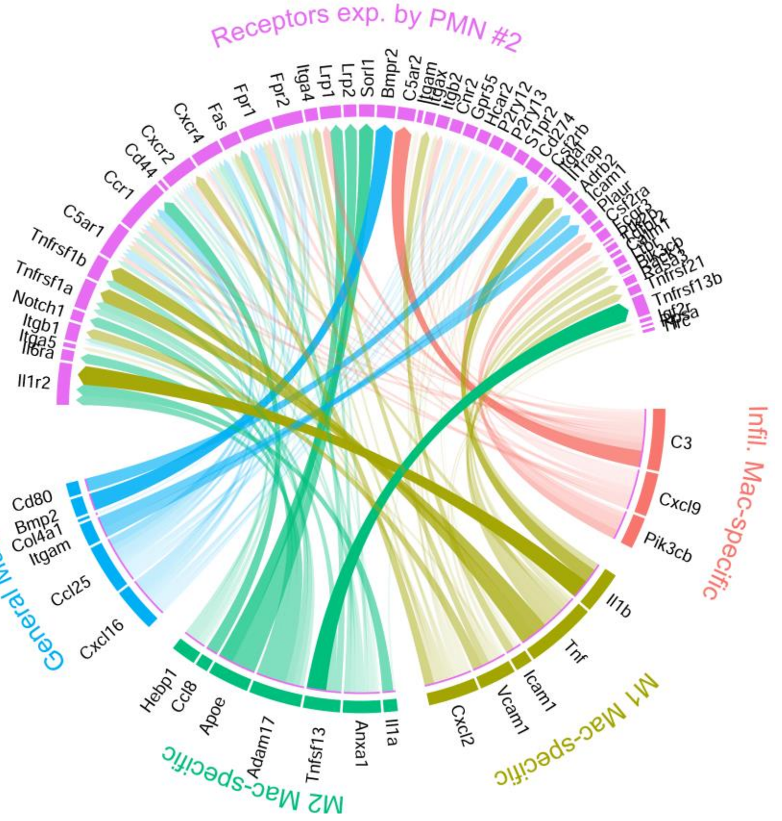
Receptors exp. by PMN #2
Cxcr4
Fas
Fpr1
Fpr2
Itga4
Lrp1
Lrp2
Sorl1
Bmpr2
C5ar2
Itgam
Itgax
Itgb2
Chr2
Gpr55
Hcar2
P2ry12
P2ry13
S1pr2
Cd274
Csf2rb
Il1rap
Adrb2
Icam1
Plaur
Csf2ra
Tnfrsf21
Tnfrsf13b
Igf2r
Cxcr2
Cd44
Ccr1
C5ar1
Tnfrsf1b
Tnfrsf1a
Notch1
Itgb1
Itga5
Il6ra
Il1r2
Infil. Mac-specific
C3
Cxcl9
Pik3cb
General Mac
Cd80
Bmp2
Col4a1
Itgam
Ccl25
Cxcl16
M2 Mac-specific
Hebp1
Ccl8
Apoe
Adam17
Tnfsf13
Anxa1
Il1a
M1 Mac-specific
Cxcl2
Vcam1
Icam1
Tnf
Il1b

**b**

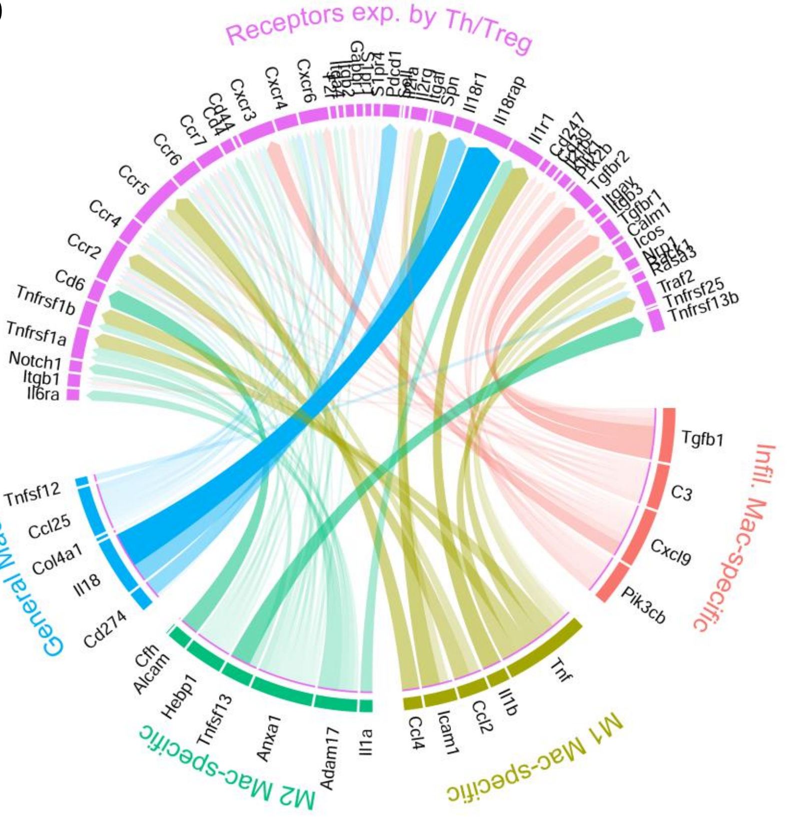
Receptors exp. by Th/Treg
Cxcr6
Cxcr4
Cxcr3
Cd44
Cd4
Ccr7
Ccr6
Ccr5
Ccr4
Ccr2
Cd6
Tnfrsf1b
Tnfrsf1a
Notch1
Itgb1
Il6ra
S1pr4
Pdcd1
Il2rg
Itgal
Spn
Il18r1
Il18rap
Il1r1
Cd247
Tgfbr2
Itgav
Itgb3
Tgfbr1
Calm1
Icos
Nrp1
Rasa3
Traf2
Tnfrsf25
Tnfrsf13b
Infil. Mac-specific
Tgfb1
C3
Cxcl9
Pik3cb
General Mac
Tnfsf12
Ccl25
Col4a1
Il18
Cd274
M2 Mac-specific
Cfh
Alcam
Hebp1
Tnfsf13
Anxa1
Adam17
Il1a
M1 Mac-specific
Ccl4
Icam1
Ccl2
Il1b
Tnf

**Supplementary Figure 17.** Ligand-receptor-target interactions between infiltrating, M1, and M2 macrophages with PMN #2 (**a**) and Cd4+ Th/Treg cells (**b**). Based on DEG between U-IRI and IRI/CL-NX kidneys on day 14 after injury, the corresponding ligands that were significantly expressed by the infiltrating, M1, and M2 macrophages were identified and linked to their corresponding receptors (bottom panel) based on the potential target genes (now shown) for PMN #2 (**a**) and Th/Treg cells (**b**) and visualized using a chord diagram.

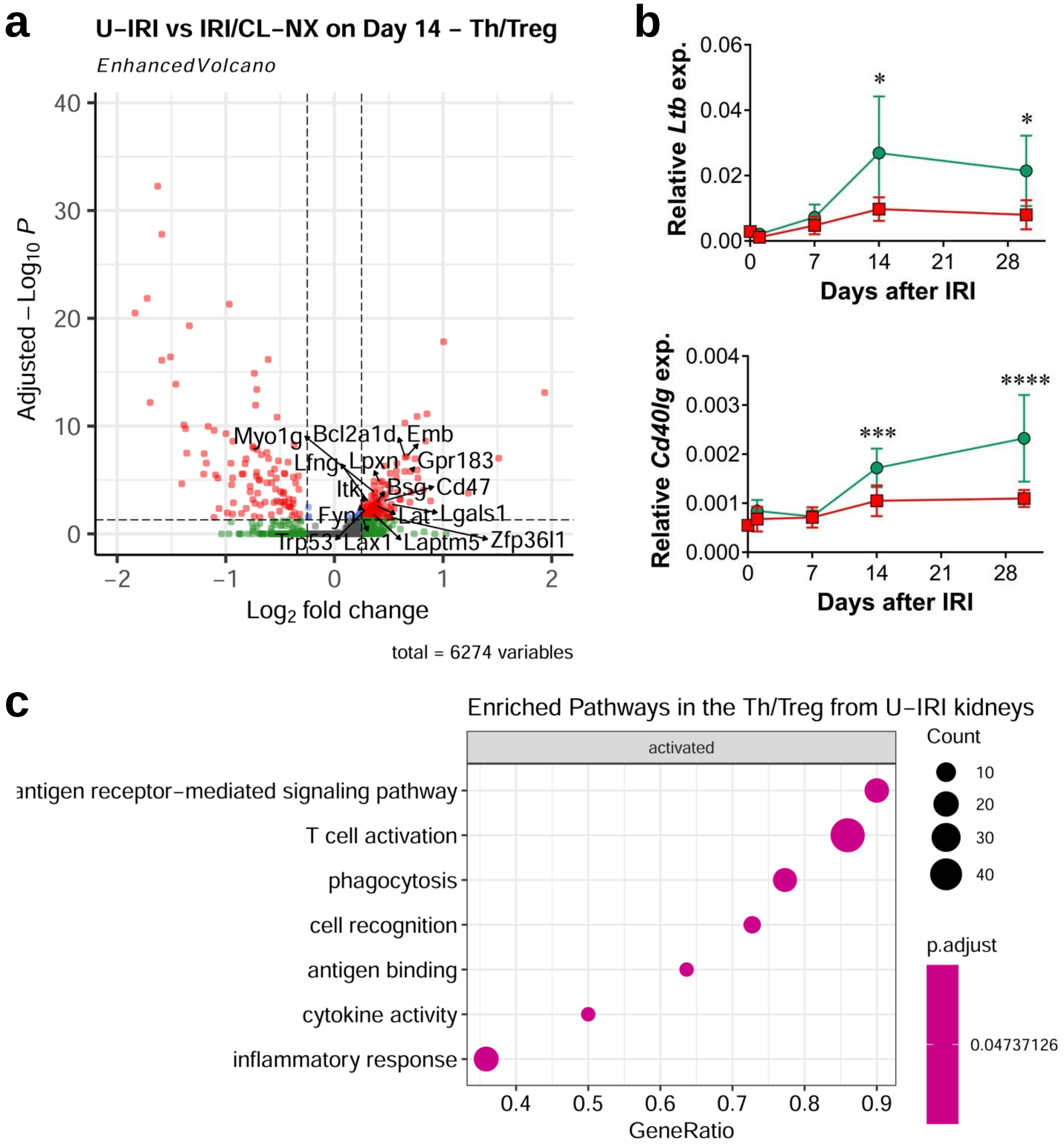


**Supplementary Figure 18. a.** Volcano plot demonstrating differential gene expression in U-IRI compared to IRI/CL-NX derived Cd4+ Th/Treg cells on day 14 after injury. **b.** Quantitative RT-PCR analysis for indicated genes was performed on whole kidney RNA harvested on day 0, 1, 7, 14, and 30 after injury. n=10 kidneys/time point/group. Two-way ANOVA summarized in Supplementary Table 1. *p<0.05, ***p<0.001, ****p<0.0001 at each time point. **c.** Based on DEG between U-IRI and IRI/CL-NX kidneys on day 14 after injury, the top relevant enriched GO terms for Cd4+ Th/Treg cells were visualized in a dot plot.

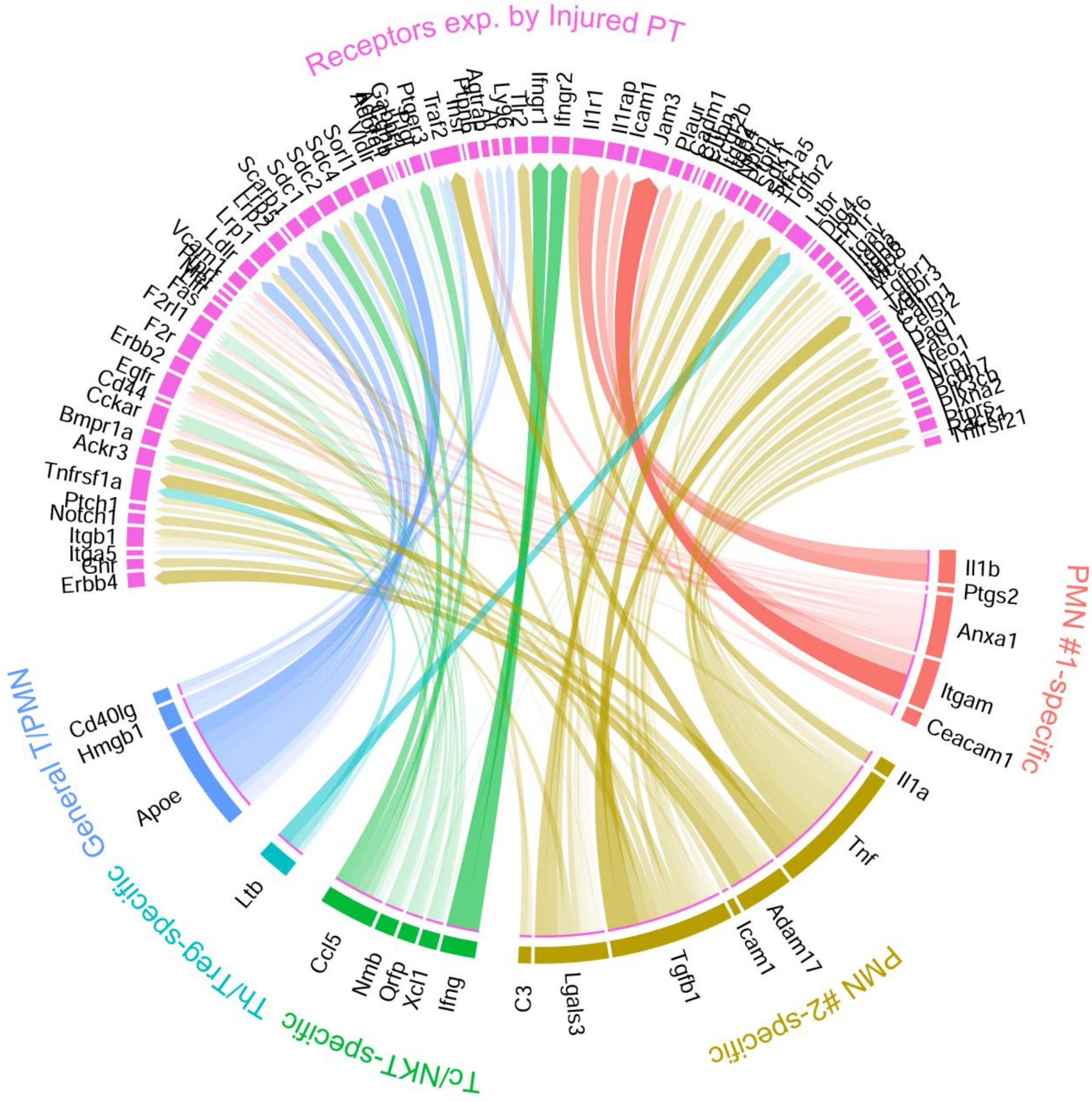


**Supplementary Figure 19.** Ligand-receptor-target interactions between PMNs, Cd8a+ Tc/NKT, and Cd4+ Th/Treg cells with injured PT cells. Based on DEG between U-IRI and IRI/CL-NX kidneys on day 14 after injury, the corresponding ligands that were significantly expressed by the PMNs, Cd8a+ Tc/NKT, and Cd4+ Th/Treg cells were identified and linked to their corresponding receptors based on the potential target genes (Figure 7h) for injured PT cells and visualized using a chord diagram.

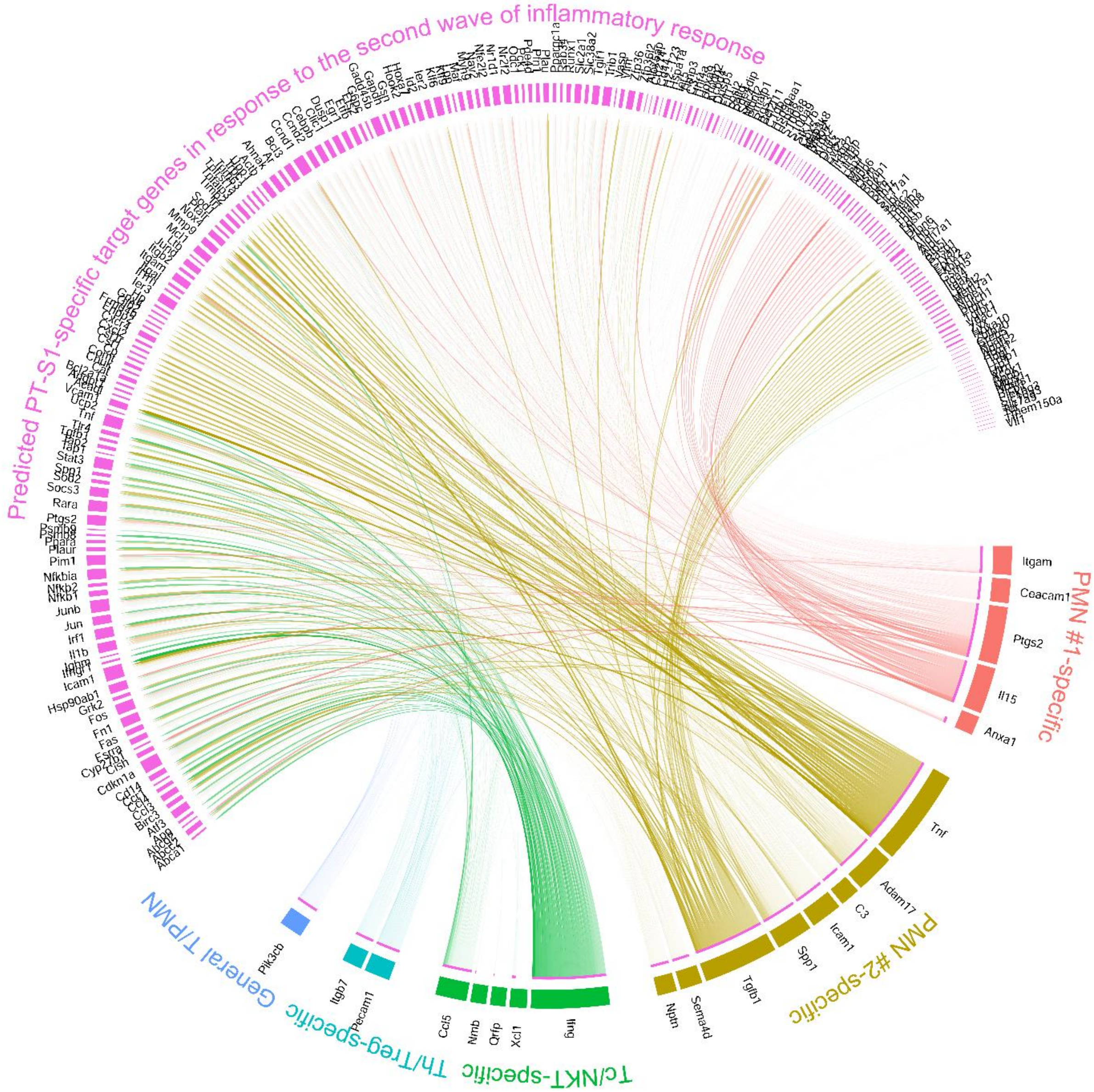


**Supplementary Figure 20.** Ligand-receptor-target interactions between PMNs, Cd8a+ Tc/NKT, and Cd4+ Th/Treg cells with PT-S1 cells. Based on DEG between U-IRI and IRI/CL-NX kidneys on day 14 after injury, the corresponding ligands that were significantly expressed by the PMNs, Cd8a+ Tc/NKT, and Cd4+ Th/Treg cells were identified and linked to their potential target genes for PT-S1 cells and visualized using a chord diagram.

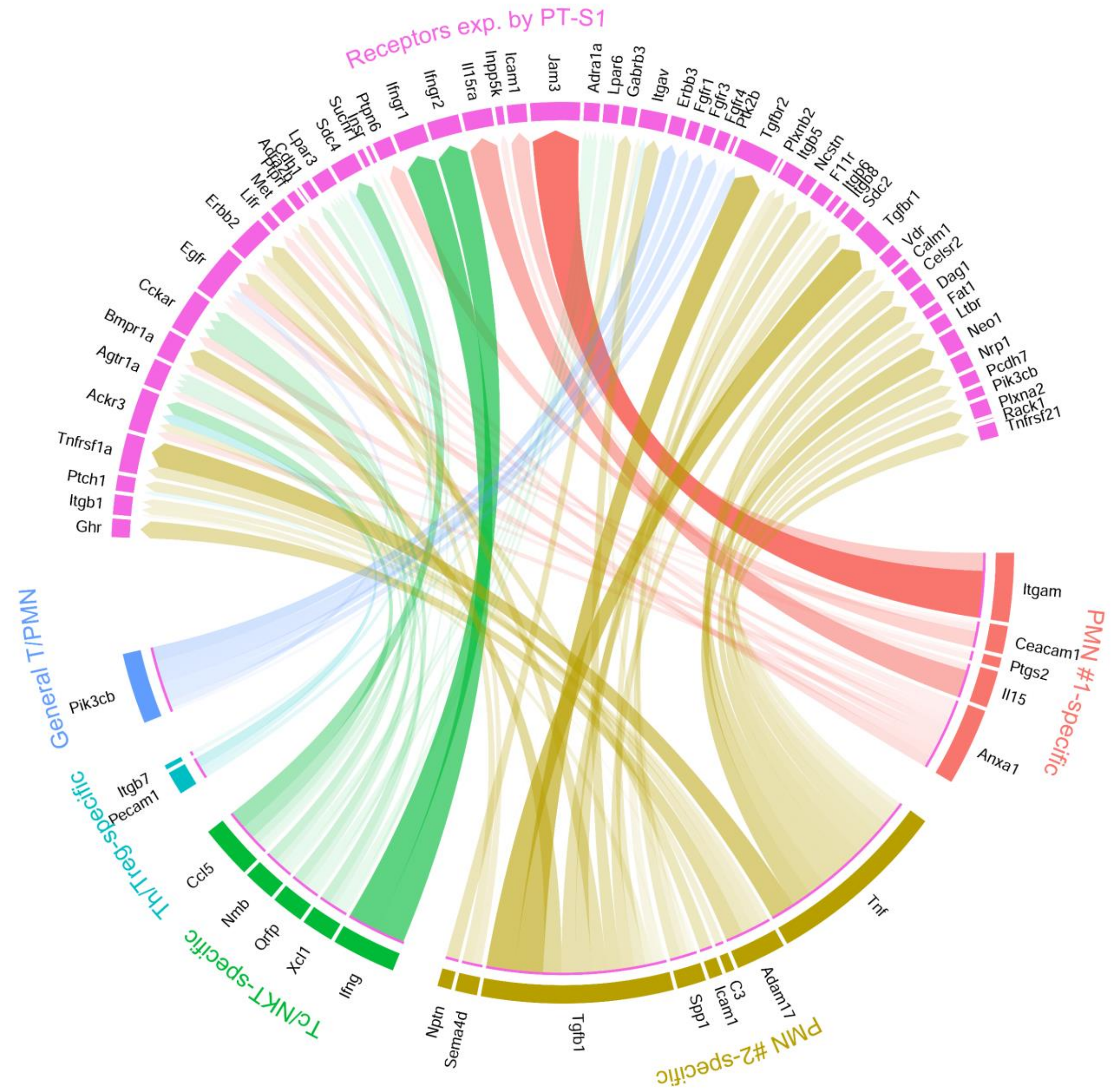


**Supplementary Figure 21.** Ligand-receptor-target interactions between PMNs, Cd8a+ Tc/NKT, and Cd4+ Th/Treg cells with injured PT cells. Based on DEG between U-IRI and IRI/CL-NX kidneys on day 14 after injury, the corresponding ligands that were significantly expressed by the PMNs, Cd8a+ Tc/NKT, and Cd4+ Th/Treg cells were identified and linked to their corresponding receptors based on the potential target genes (Supplemental Figure 20) for PT-S1 cells and visualized using a chord diagram.

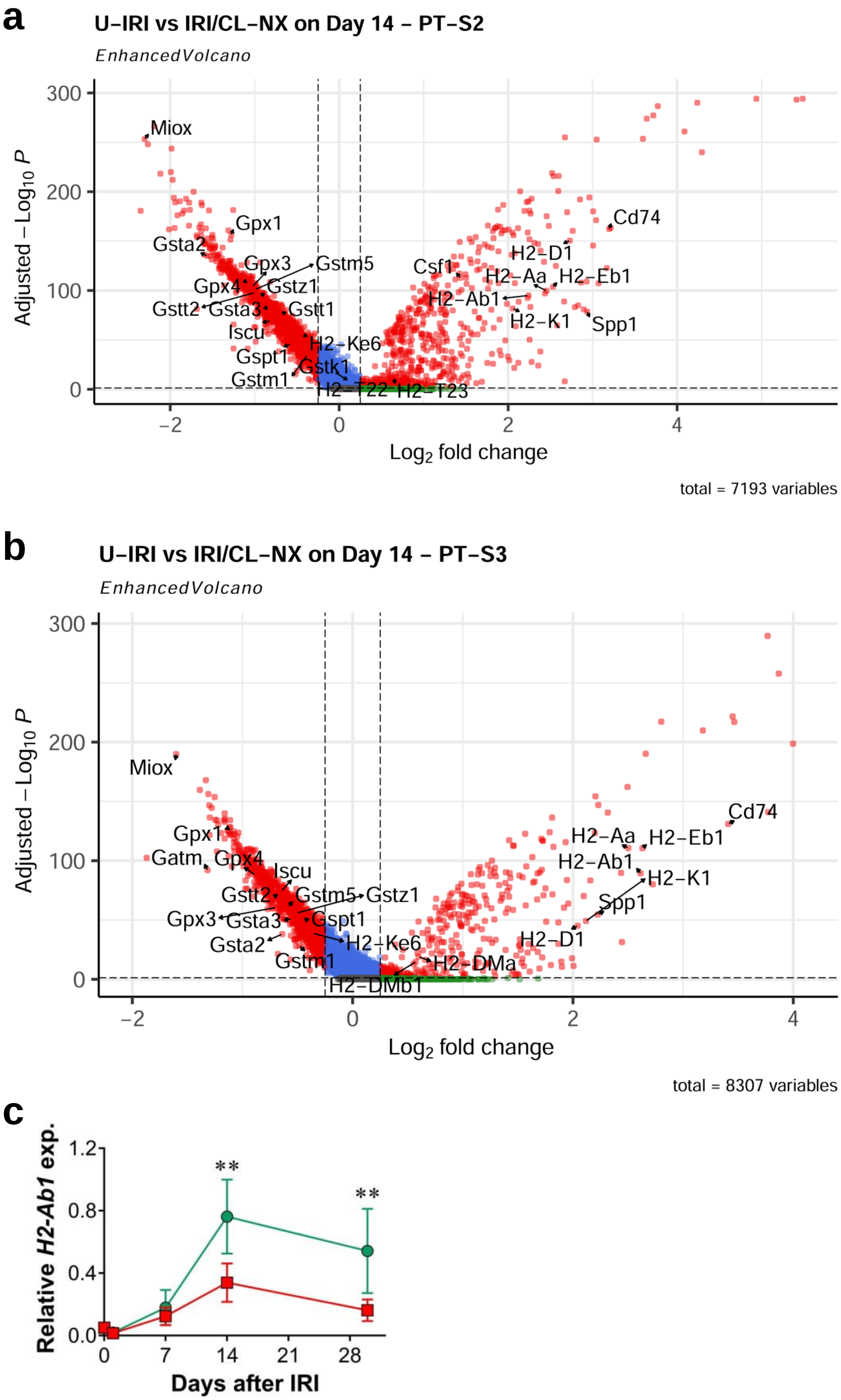
a
U–IRI vs IRI/CL–NX on Day 14 – PT–S2
EnhancedVolcano
Adjusted −Log10 P
Log2 fold change
Miox
Gpx1
Gsta2
Gpx3
Gstm5
Gpx4
Gstz1
Gstt2
Gsta3
Gstt1
Iscu
Gspt1
Gstk1
Gstm1
H2–Ke6
H2–T22
H2–T23
Csf1
H2–D1
H2–Aa
H2–Eb1
H2–Ab1
H2–K1
Spp1
Cd74
total = 7193 variables
b
U–IRI vs IRI/CL–NX on Day 14 – PT–S3
EnhancedVolcano
Adjusted −Log10 P
Log2 fold change
Miox
Gpx1
Gatm
Gpx4
Iscu
Gstt2
Gstm5
Gstz1
Gpx3
Gsta3
Gspt1
Gsta2
H2–Ke6
Gstm1
H2–DMa
H2–DMb1
H2–Aa
H2–Eb1
H2–Ab1
H2–K1
Spp1
H2–D1
Cd74
total = 8307 variables
c
Relative H2-Ab1 exp.
Days after IRI
**
**

**Supplementary Figure 22. a** and **b.** Volcano plots demonstrating differential gene expression in U-IRI compared to IRI/CL-NX derived PT-S2 (**a**) and PT-S3 (**b**) cells on day 14 after injury. **c.** Quantitative RT-PCR analysis for *H2-Ab1* was performed on whole kidney RNA harvested on day 0, 1, 7, 14, and 30 after injury. n=10 kidneys/time point/group. Two-way ANOVA summarized in Supplementary Table 1. **p<0.01 at each time point.

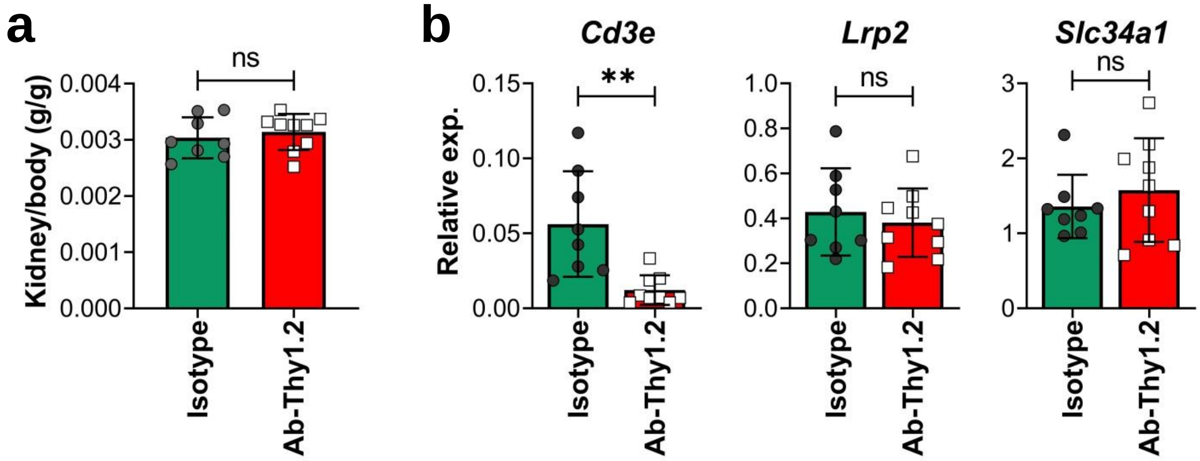


**Supplementary Figure 23.** Depletion of T cells alone did not prevent kidney tubule atrophy. WT mice were treated as described in Methods with either isotype or antibodies (Ab)-against Thy1.2 beginning 5 days after U-IRI and sacrificed on day 30. **a.** Kidney-to-body weight ratios on day 30 following U-IRI. n = 8-9 kidneys/group. ns, not statistically significant. **b.** Quantitative RT-PCR analysis for *Cd3e*, *Lrp2*, and *Slc34a1* was performed on whole kidney RNA from U-IRI mice. N = 8-9 kidneys/group. **p < 0.01; ns, not statistically significant.

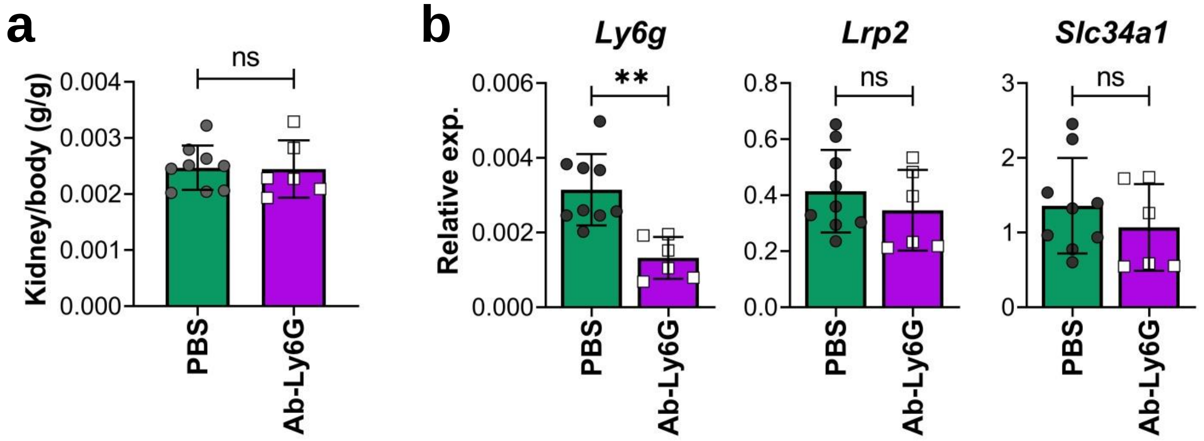


**Supplementary Figure 24.** Depletion of neutrophils alone did not prevent kidney tubule atrophy. WT mice were treated as described in Methods with either PBS or antibodies (Ab)-against Ly6G beginning 5 days after U-IRI and sacrificed on day 30. **a.** Kidney-to-body weight ratios on day 30 following U-IRI. n = 6-9 kidneys/group. ns, not statistically significant. **b.** Quantitative RT-PCR analysis for *Ly6g*, *Lrp2*, and *Slc34a1* was performed on whole kidney RNA from U-IRI mice. N = 6-9 kidneys/group. **$p < 0.01$; ns, not statistically significant.

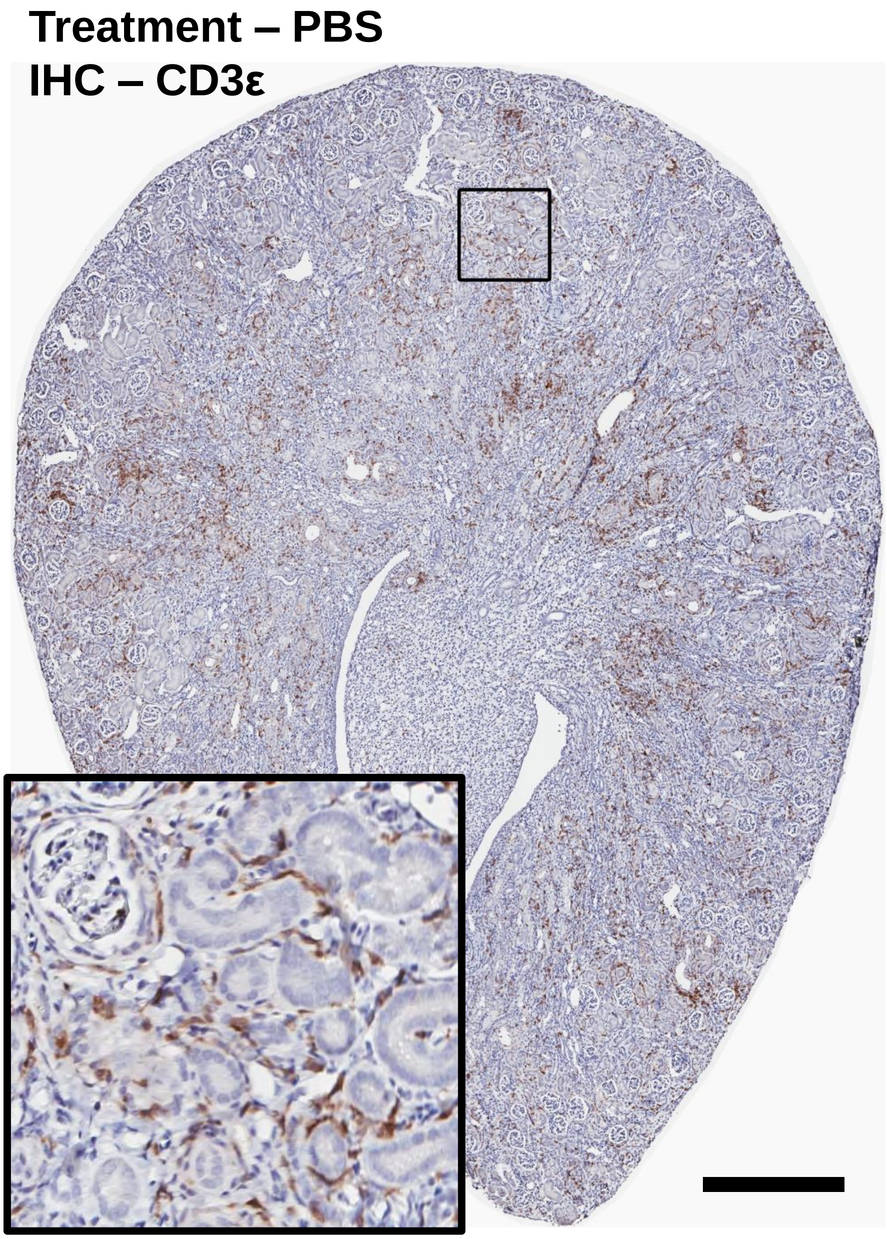


**Supplementary Figure 25.** The kidney sections on day 30 after U-IRI (treated with PBS) were immunostained with CD3ε. Representative image of whole kidney sections was shown and higher magnification image was shown in the box. Scale bar, 400 μm.

**Treatment – Ab-Thy1.2 + Ab-Ly6G**
**IHC – CD3ε**

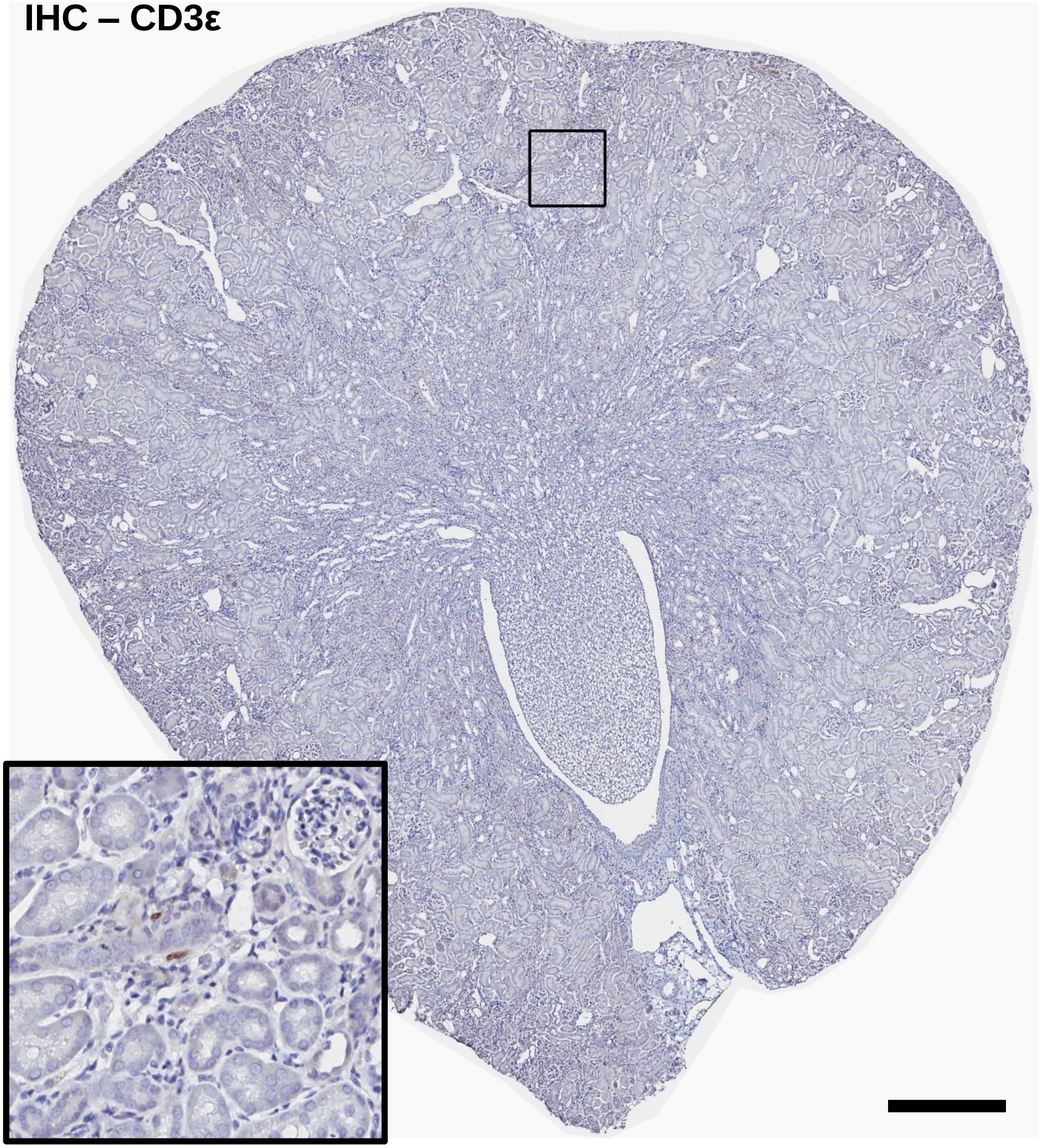

**Supplementary Figure 26.** The kidney sections on day 30 after U-IRI (treated with antibodies (Ab)-against Thy1.2 and Ly6G) were immunostained with CD3ε. Representative image of whole kidney sections was shown and higher magnification image was shown in the box. Scale bar, 400 μm.

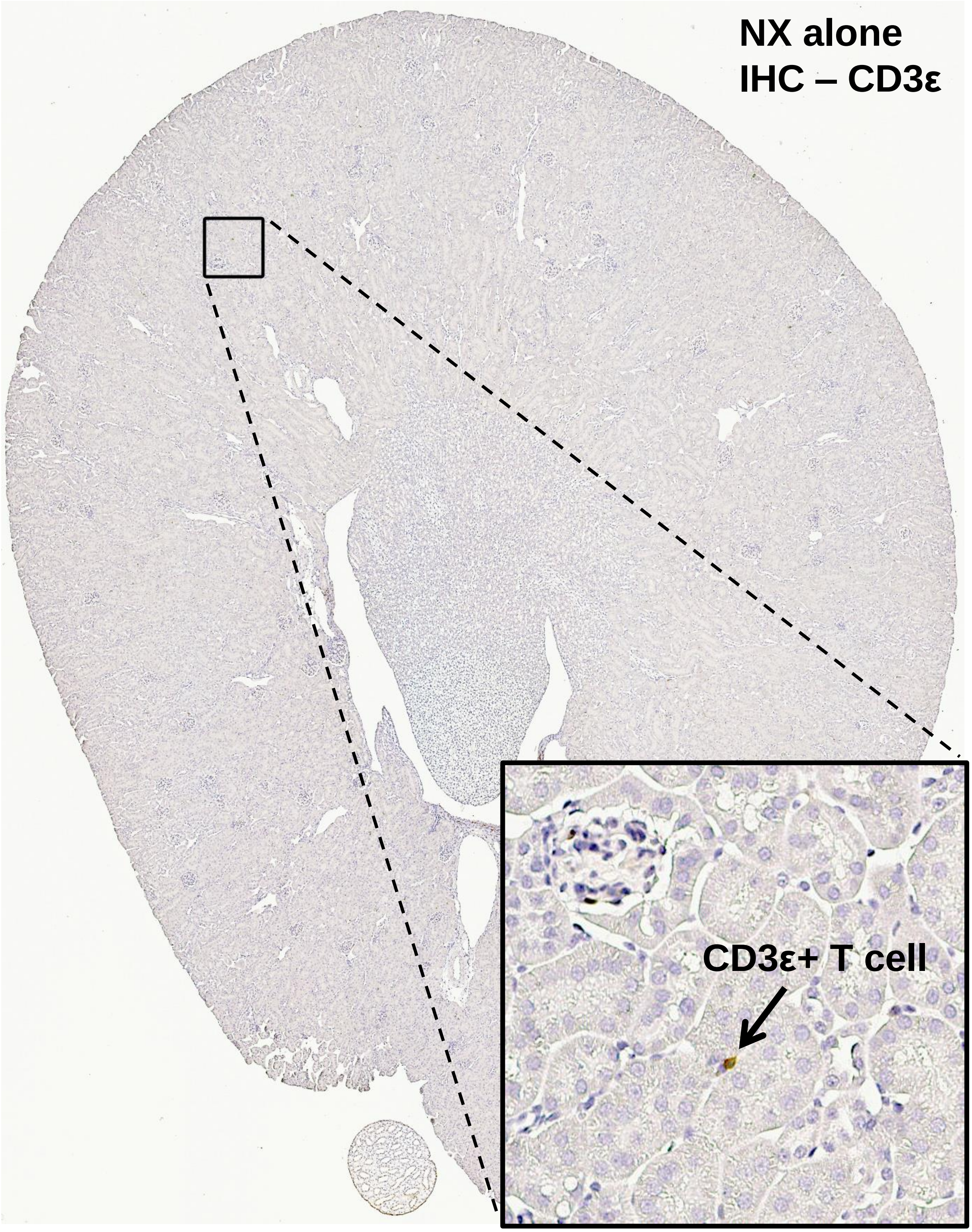


**Supplementary Figure 27.** The kidney sections on day 30 after CL-NX alone were immunostained with CD3ε. Representative image of whole kidney sections was shown and higher magnification image was shown in the box.
